# Pluripotent and differentiated cells display distinct genomic and nuclear landscapes of genome folding variability

**DOI:** 10.64898/2026.11.16.766639

**Authors:** Xiangru Huo, Ye Wang, Liyan Yang, Jiang Min, Jack Huey, René Maehr, Johan H. Gibcus, Frank Alber, Job Dekker

## Abstract

Chromosome folding varies between cells. It is not known whether cell types differ in the extent of folding variability, whether distinct chromosomal regions in the same cell type display different levels of variability, and how folding variability relates to sub-nuclear location and gene expression. We combined two single-cell Hi-C approaches with 3D genome modeling of pseudo-single cells (IGM), DNA FISH, chromosome painting, and single-cell RNA-seq across a five-stage differentiation trajectory from human embryonic stem cells to hepatocyte-like cells. In all cell types folding variability correlates with compartment status, is highest towards the nuclear interior, decreases as a function of radial position in the nucleus, and is the lowest at the periphery. Loci associated with nuclear landmarks such as speckles, and especially the nuclear periphery display more consistent subnuclear positioning in all cell stages. Moreover, pluripotent and differentiated cells display different overall levels of chromosome folding variability: in H1 pluripotent cells the entire genome displays higher folding and positional variability. In contrast, in differentiated cells folding variability is generally lower along the genome and the number of loci that become consistently positioned at the nuclear periphery increases sharply. The two distinct patterns of folding variability were correlated with cell-to-cell gene expression variation: In pluripotent cells gene expression variation is large, pervasive and seen throughout the genome and the nucleus. In differentiated cells cell-to-cell variation in gene expression is much lower and restricted mostly to genes located near the center of the nucleus, consistent with the restricted pattern of chromosome folding variation. These findings reveal two distinct genomic states: pluripotent cells maintain a variably folded, spatially permissive genome and transcriptional plasticity, whereas differentiated cells mature into a spatially deterministic, more consistently folded organization that correlates with more restricted transcriptional programs.

## Introduction

The spatial arrangement of the genome within the interphase cell nucleus is determined by multiple processes^1–4^. First, in general active and inactive chromatin domains, ranging in size from tens of kb to several Mb, are spatially segregated through a process referred to as compartmentalization. This occurs both along single individual chromosomes, and between different chromosomes. This process is related to phase separation that is driven through attractions between chromatin domains of similar type, and/or repulsion of domains of different type^5,6^. Second, chromatin domains associate with several different nuclear bodies and sub-nuclear structures and such associations are correlated with formation of chromosomal compartments^7^. For instance, inactive and closed chromatin domains are often associated with the nuclear periphery (e.g., Lamin Associated Domains (reviewed in van Steensel^8^), while a subset of highly expressed GC-rich regions associate with nuclear speckles^9,10^. Third, the cohesin complex actively generates chromatin loops across the genome^11,12^. This process is fast (with in vitro chromatin extruded at several kb per second^13^; and in vivo somewhat slower, 0.1-0.5 kb per second^14^) and produces dynamically positioned loops that can be up to several hundred kb in size. This process also generates more stable loops between loci where extrusion is stalled, i.e., at CTCF-bound sites^15–17^. However, even these more stable loops are not permanent and have an estimated lifetime of up to 30 minutes^14,18^. The final spatial arrangement of the genome is the result of the combined activity of all these processes acting simultaneously.

While these processes act in most if not all cells, chromosome folding and nuclear organization is cell type specific, and changes when cells change fate or differentiation state. The reason for this is that different regions of the genome are active or inactive in different cell types, leading to changes in genome-wide profiles of compartmentalization, and to differences in which specific loci are associated with nuclear structures such as the nuclear lamina. Similarly, patterns of cohesin-mediated loop extrusion are determined to a significant extent by cis-elements whose activities are cell-type specific. For instance, cohesin loading, pausing, blocking and unloading are regulated by elements such as enhancers, promoters, 3’ ends of genes, and sites bound by CTCF respectively, leading to cell type-specific cohesin “traffic patterns” and resulting chromatin loops^19–22^. These observations suggest that chromosome folding principles are conserved across cell types and that nuclear and genome organization only differ between different cell types because they differ in their epigenomic patterns and activity of cis-elements that in turn guide these principles.

The polymer nature of very long chromosomes, and the mechanisms of their folding are predicted to naturally lead to large variation in their spatial disposition, even for otherwise identical cells. One reason is that the nucleus organizes at least in part through self-assembly, driven by the different mechanisms described above, which will produce stochastic and heterogeneous conformations such as compartments as emergent properties (see Misteli^1^ for an extensive discussion of these phenomena). For instance, compartmentalization through phase separation will lead to stochastic higher order assemblies: while in all cells active domains will be positioned near other active domains, which specific pairings occur will vary^23^. Besides this biophysical basis for heterogeneity in chromosome folding, molecular processes such as loop extrusion, transcription and chromatin remodeling will add further variations in genome organization over time in single cells. These processes move segments of chromatin at minute time scales and thereby create variation within single cells.

Heterogeneity and stochasticity in chromosome folding and nuclear organization have been observed experimentally. For instance, high throughput fluorescence in situ hybridization has shown that chromosome conformation, and locus positioning shows large variation between cells of the same type^24–26^. Similarly, single cell Hi-C showed that cells differ in patterns of inter-chromosomal interactions, and compartmental interactions^27–31^. Modeling has shown that population-based Hi-C interaction maps can be explained as the sum of many single cell interaction maps that differ in precise patterns of inter- and intra-chromosomal interactions^32–35^.

Given that the general processes that fold chromatin appear to apply to all cells, and that these activities produce stochastic and heterogeneous chromosome structures we were interested to explore whether chromosome folding variability is also similar for different cell types and, within a cell type, across the genome. In addition, we were interested in whether and how chromosome folding variability would be related to cell-to-cell variation in gene expression. Previous studies have shown that gene expression across the genome can be variable between cells and vary over time within cells^36–40^. This led us to ask the following questions: 1: Is chromosome folding variability between cells constant along the genome, or are specific regions more variable than others? 2: Does any such variability in folding along the genome depend on cell state? For instance, does folding variability change as cells progress from pluripotency to lineage commitment? And 3: Is chromosome folding variability related to gene expression variability, and does this relationship change as cells change fate (**Fig. 1a**)? We find that embryonic stem (ES) cells and differentiated cells differ in chromosome folding variability, in correlation with differences in gene expression heterogeneity. Single cell 3D genome modeling indicates that nuclear organization in ES cells and differentiated cells differ in the extent to which loci become anchored to nuclear landmarks such as the nuclear lamina and speckles giving rise to two modes of interphase nuclear organization characterized by distinct chromatin folding variability landscapes.

**Fig 1.**
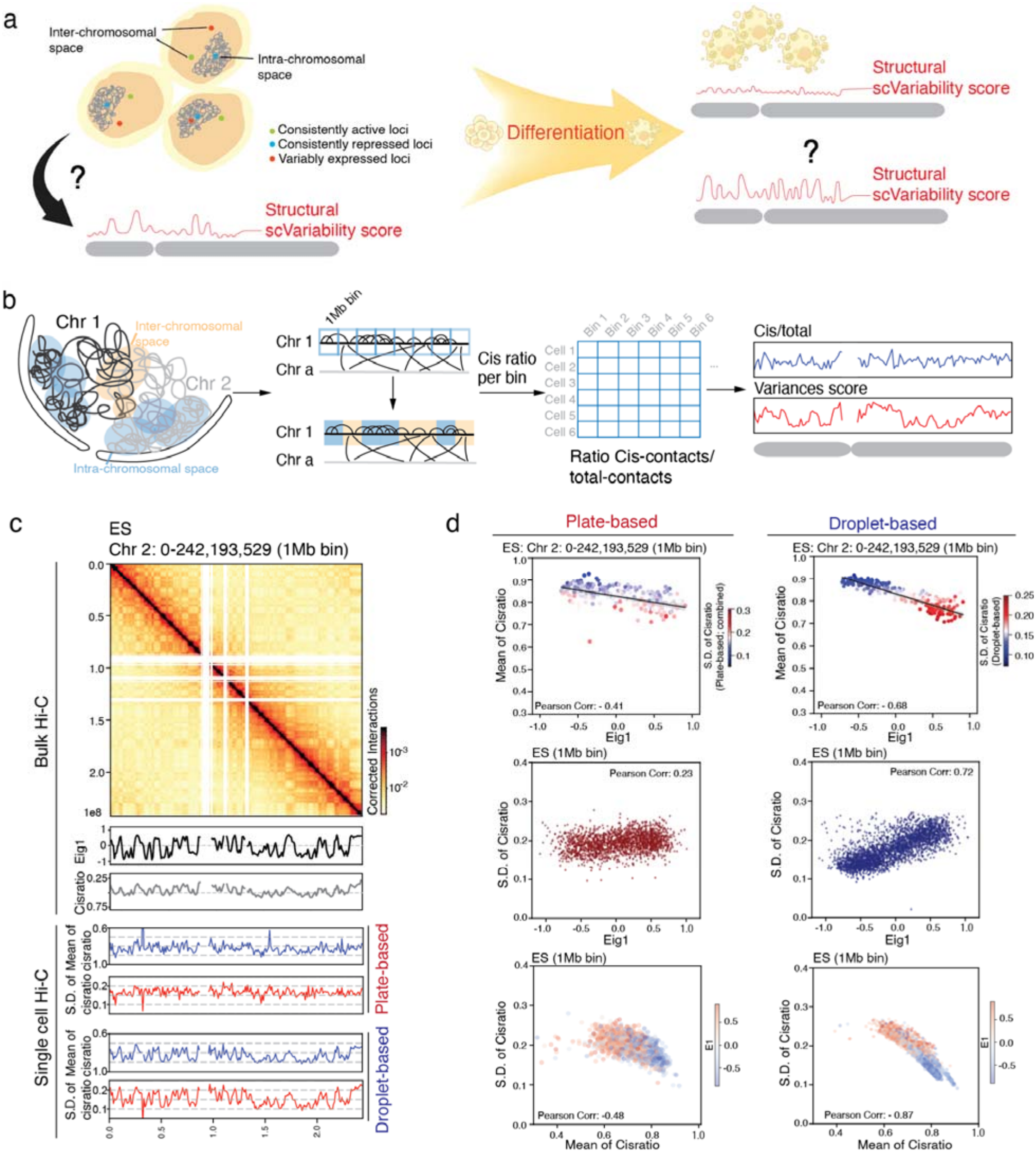
Single-cell Hi-C and integrative analysis reveal cell-to-cell variability in genome folding. **(a)** Schematic illustrating the central question: How variable is genome folding and genomic loci positioning between single cells and can this information be linked to features of genome function? Does this change during differentiation? **(b)** Conceptual overview of how cis-to-total contact ratio per genomic bin is calculated in single cells as a measurement of distribution of genomic loci with respect to chromosome territory. Variation in this ratio across cells reflects genome-folding variability at each locus. Yellow shading marks inter-chromosomal space and blue shading marks intrachromosome space. In this schematic, “Chr A” represents any autosome other than Chr 1, which is the target chromosome used here to illustrate the cis-to-total contact ratio calculation. **(c)** Genome folding variability and compartmentalization in ES cells along chromosome 2. Top: Bulk Hi-C contact map (0–242 Mb, 1 Mb binning) showing corrected interaction frequencies. Middle: Bulk-derived compartment eigenvector (Eig1) and cis-to-total contact ratio profiles. Bottom: Plate-based and droplet-based single-cell Hi-C-derived profiles showing the mean cis-to-total contact ratio per bin (blue) and standard deviation (S.D.) of the cis-to-total contact ratio across cells (red). (Plate scHi-C uses integrated ES batch 1 and batch 2; plate scHi-C and droplet Hi-C uses filtered contacts distance > 0.3kb). **(d)** Relationships among the mean of cis-to-total contact ratio, compartment eigenvector, and S.D. of cis-to-total contact ratio. Across 1-Mb bins on chromosome 2 in ES cells (left: plate-based; right: droplet-based), Top row plot-compartment eigenvector (Eig1) versus mean of cis-to-total contact ratio, with the dots colored by the S.D. of the cis-to-total contact ratio (color scale: blue = low variability, red = high variability). Middle row plot-compartment eigenvector (Eig1, positive values refer to A compartments, negative values refer to B compartments) and S.D. of cis-to-total contact ratio. Bottom row plot-the mean of cis-to-total contact ratio and S.D. of cis-to-total contact ratio, with dots colored by compartment eigenvector (Eig1) (color scale: blue = compartment B, red = compartment A).(Plate scHi-C uses integrated ES batch 1 and batch 2; plate scHi-C and droplet Hi-C uses filtered contacts distance > 0.3kb).

## Results

### Detecting variability in chromosome folding in single cells

We implemented two variants of the single cell Hi-C procedure. First, we sorted fixed single cells in wells of a 96-well plate that were preloaded with a lysis buffer containing Nesprin-coated beads. Cells were then lysed and the nuclei were retained on beads. The supernatant containing the cytoplasm was removed, and could be used to identify RNAs from that cell using a standard Smart-seq3 protocol^41^ (Methods, Supplement). Here we focus on analyzing chromosome organization in remaining nuclei using an adapted single cell Hi-C protocol (Methods). Briefly, chromatin is first fragmented with NlaIII, DNA is re-ligated, and then prepared for sequencing using Tn5 tagmentation. The latter step adds Illumina sequencing adapters, and nucleus-specific barcodes are added during the subsequent PCR amplification. Second, we adapted the droplet Hi-C procedure developed by the Ren lab^31^. Fixed cells were lysed, fragmented with NlaII, MboII, and DpnII, and DNA was re-ligated. Tn5 tagmentation was performed to add ATAC adapters. Next, ~5,000 single nuclei were loaded into a single 10X genomics microfluidic chip. Single nuclei were captured in individual droplets, each containing cell-specific PCR primers. Then DNA fragments were recovered from droplets and an Illumina sequencing library was constructed. To validate the single-cell Hi-C assays, we benchmarked them using exponential and synchronized HAP1 cell cultures, demonstrating that both methods can accurately distinguish interphase from mitotic cells based on characteristic differences in interaction distance distributions and compartmentalization strength (**Supplemental Note 1, 2; Extended Data Fig. 1a-f**).

Chromosome territories (CTs), as observed by chromosome painting, are reflected in Hi-C data by a strong enrichment of intra-chromosomal interactions relative to inter-chromosomal interactions^42,43^. We leveraged this principle to quantify whether individual genomic loci are embedded within, or looped out of their CT in single cells, a phenomenon that is related to gene expression^44,45^. Further, we found that in human embryonic stem cells territoriality is low and chromosomes mingle extensively^44^. Territoriality increases as cells differentiate, making territoriality a quantifiable feature that may vary across cell states and between cells.

Specifically, for each genomic 1Mb bin we calculated the ratio of cis-chromosomal interactions to total interactions (cis-to-total contact), which serves as a proxy for the spatial positioning of that locus relative to its CT. A high cis-to-total contact ratio indicates that a locus predominantly interacts with regions on the same chromosome, consistent with localization within intra-chromosomal space and embedding inside the CT. A low cis-to-total contact ratio reflects increased inter-chromosomal interactions, indicating positioning in the inter-chromosomal space and looping out of the CT. We computed this metric for all genomic bins in each single cell, yielding a genome-wide cis-to-total interaction profile per cell. By aggregating these profiles across cells, each bin is characterized by a distribution of cis-to-total values, from which we derived (i) the mean cis-to-total contact ratio, representing the population-average spatial positioning of the locus relative to its CT (with active regions expected to show lower cis-to-total contact ratio to reflect higher inter-chromosomal space localization, and (ii) the standard deviation (S.D.) of the cis-to-total contact ratio, capturing cell-to-cell variability in locus positioning relative to its CT (**Fig. 1b**). This framework enables us to assess both the population-average organization and single-cell variability of genome folding with respect to CT positioning for each bin (locus) along the genome.

To evaluate this computational framework, we calculated S.D. of cis-to-total contact ratios in interphase cells from an exponentially growing HAP1 culture, and a synchronized culture enriched in prometaphase-arrested HAP1 cells respectively. We observed that compared to interphase, prometaphase cells exhibited a narrower S.D. distribution across genomic bins, reflecting more uniform folding variability across the genome as chromosomes condensed to form rod-shaped structure in mitosis cells. (**Extended Data Fig. 1g**).

### Annotation of variably and consistently folded genomic regions in H1-hESC

We implemented single-cell Hi-C on human embryonic cells (H1-hESCs), collecting ~600 cells from two batches using the plate-based procedure and ~5,000 cells using the droplet-based Hi-C procedure^31^. After filtering for cells with more than 20,000 contacts and selecting interphase cells based on the short-range-to-mitotic interaction ratio, we retained around 300 plate-based and 1,500 droplet-based single cells (**Extended Data Fig. 1h**). To remove false-positive contacts arising from un-ligated fragments, we filtered out short-range interactions using a library-specific distance cutoff (**Supplementary Note 2, supplemental Fig. 1**). Next, we calculated cis-to-total contact ratio for all genomic bins across all chromosomes, and measured S.D. and mean of cis-to-total contact ratio across single cells to annotate variability of genome folding as we described above (**Fig. 1c, d**). The annotations were reproducible across the two single cell Hi-C procedures (**Extended Data Fig. 1i**).

We compared the mean and S.D. of the cis-to-total contact ratio annotated from scHi-C with the eigenvalues derived from bulk Hi-C compartment analysis. As expected, the mean cis-to-total contact ratio was negatively correlated with the compartment’s eigenvalue. The active region (compartment A) shows lower cis-to-total contact ratio, consistent with active expressed genes preferentially looping out of their CT, while repressed regions (compartment B) showed higher cis-to-total contact ratio, meaning they remained mostly embedding within their CT on average (**Fig. 1d**)^45,46^.

Cell-to-cell folding variability was also non-uniform across the genome. Active regions (compartment A) exhibited higher cell-to-cell folding variability in the cis-to-total contact ratio, whereas repressed or closed chromatin regions (compartment B) exhibited greater consistency in genome folding. (**Fig. 1d**). Because the cis-to-total contact ratio is a bounded proportion, its raw variability (S.D.) is partly constrained by its own mean — bins with a mean ratio near 0 or 1 have limited numerical range in which to vary, and consequently show a lower S.D. independent of any true underlying biological heterogeneity. Since compartment B bins have a higher mean cis-to-total ratio closer to this boundary, we considered whether the apparent A-versus-B difference in variability could simply reflect this mean-driven effect rather than genuine biology. To test this, first, we modeled each bin’s expected standard deviation as a function of its mean cis-to-total contact ratio (see Methods) and used the residual between observed and expected standard deviation (resid_SD) as a mean-corrected variability score. This corrected analysis confirmed the original finding: compartment A bins retained significantly higher resid_SD than compartment B bins (**Extended Data Fig. 1j**). Furthermore, we applied a more stringent statistical workflow in which we used a hierarchical statistical model to decouple the biological variability and sampling noise, followed by removing the effect of mean from biological variance, and filter the contact depth matched cells (Supplemental Note 3,**supplemental Fig. 2-4**). These results all indicate that the elevated cell-to-cell variability of active chromatin is not an artifact of sampling noise, sequence reads depth or its intermediate mean cis-to-total contact ratio.

**Figure 2:**
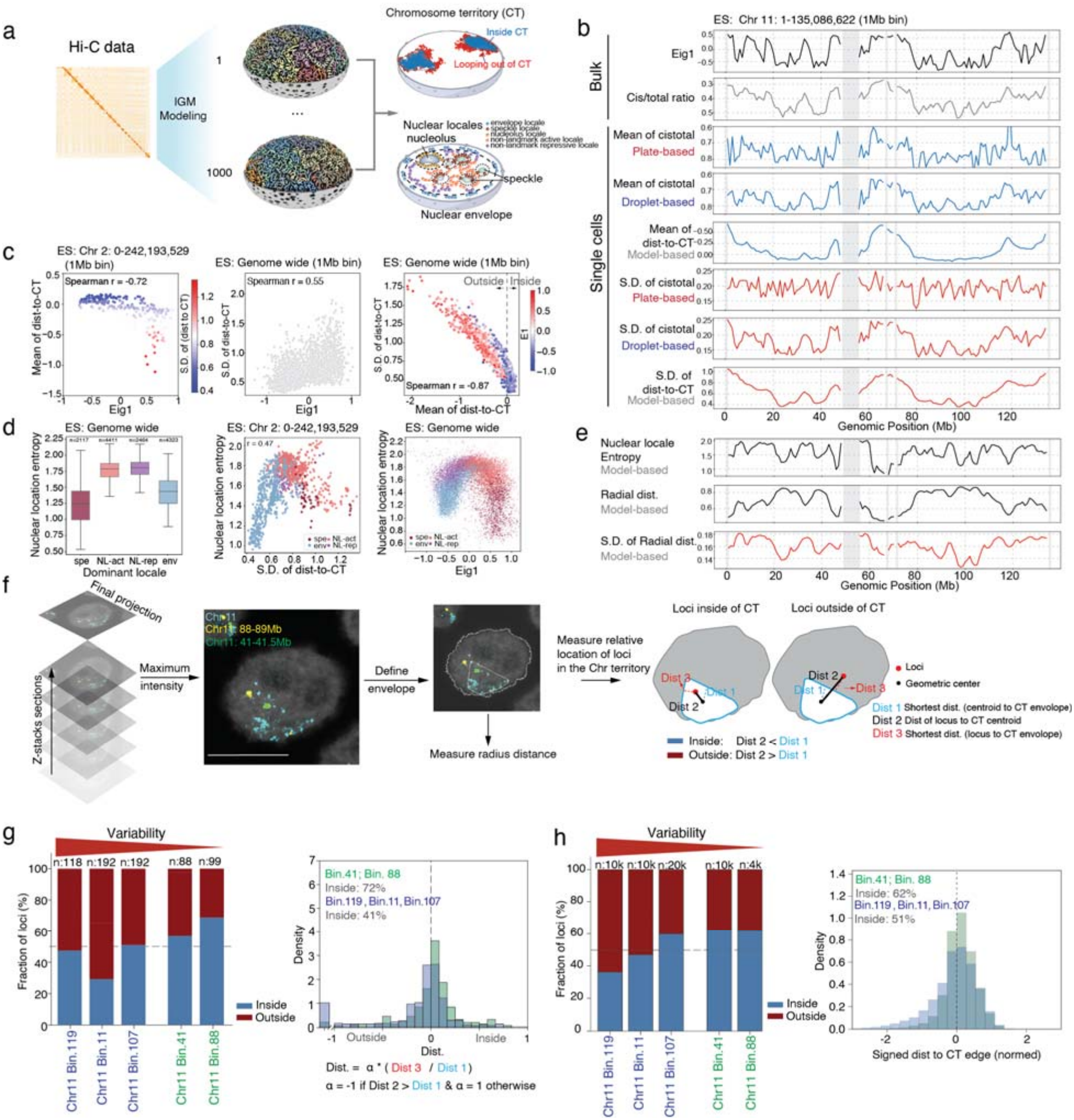
Integrative genome modeling and DNA FISH validate genome folding variability and spatial positioning in single cells. **(a)** Integrative genome modeling (IGM) framework. Left: Ensemble modeling scheme for generating 1,000 diploid genome structures (200-kb resolution) constrained by Hi-C data. Right, top: Example chromosomes showing the core chromosome territory (blue), defined by the convex hull of loci with the highest cis-to-total contact ratios, and genomic regions extending beyond the core territory boundary (red). Regions outside the boundary surface have negative signed CT-boundary distances (dist-to-CT) and are considered to loop out of the chromosome territory. Right, bottom: Schematic representation of chromatin colored according to their primary nuclear locale assignments. Nuclear speckles, nucleoli, and the nuclear envelope are shown together with chromatin regions assigned to speckle (spe), nucleolar (nuc), envelope (env), non-landmark active (NL-active), or non-landmark repressive (NL-rep) locales. **(b)** Multi-platform genome folding profiles. Chromosome 11 tracks comparing bulk Hi-C (Eig1 and cis-to-total ratio) with mean and S.D. of cis-to-total ratios derived from plate-based scHi-C (red fonts), droplet-based scHi-C (blue fonts), and IGM models with mean and S.D. of dist-to-CT (grey fonts). (Plate scHi-C uses integrated ES batch 1 and batch 2; plate scHi-C and droplet Hi-C uses filtered contacts distance > 0.3kb). **(c)** Compartment status vs. structural variability. Chr 2 scatter plots in ES cells relating bulk Eig1, mean of dist-to-CT, and S.D. of dist-to-CT (variability). Left: Eig1 vs. mean of dist-to-CT, colored by S.D. (blue = low, red = high). Middle: Eig1 vs. S.D. of dist-to-CT. Right: Mean of dist-to-CT vs. S.D. of dist-to-CT, colored by Eig1 (blue = B compartment, red = A compartment). **(d)** Nuclear locale entropy dynamics. Per-bin (200 Kb) autosomal analyses colored by dominant locale (P argmax. None of the genomic loci were canonically nucleolar. We therefore excluded the nucleolar locale from subsequent entropy analyses). Left: nuclear location entropy boxplots across dominant locales. Boxes show median and interquartile range; whiskers extend to 1.5×IQR; outliers plotted as gray points. *Middle:* nuclear location entropy vs. S.D. of dist-to-CT for chromosome 2. *Right:* nuclear location entropy vs. bulk Hi-C Eig1. Dots in scatter plots are colored by the dominant nuclear locale class assigned per bin. At all three panels, bins are restricted to autosomes (chr1–22), excluding centromere and repetitive regions that lack a valid bulk Hi-C E1 value. **(e)** Chr 11 locus positioning tracks. Aligned profiles of model-derived nuclear location entropy (top), mean ensemble radial distance (middle; 0 = nuclear center, 1 = envelope) and S.D. of ensemble radial distance (bottom). Centromeric and unmapped regions are masked in gray. **(f)** Workflow for DNA FISH imaging and quantitative spatial analysis. Left: Z-stack sections of nuclei were acquired and processed by maximum intensity projection to generate final images. Middle: Example projection showing loci on chromosome 11 (bin 88–89 Mb, yellow; bin 41–41.5 Mb, cyan). Right: Chromosome territory boundaries and nuclear envelope were defined (white dashed line). For each locus, two measurements were obtained:1) relative radial position within the nucleus; and 2) relative positioning to CT, using three distances -- Dist 1: shortest distance from centroid of CT to the CT envelope. Dist 2: distance from centroid of CT to the locus. If the distance from locus to centroid of CT < shortest radius distance, then define the locus lies inside the CT; if the distance from locus to centroid of CT > longest radius distance, then defined locus lies outside the CT; otherwise, the locus lies on the edge. Scale bar: 10μm. **(g)**. *Left:* the relative distribution of loci to CT from DNA FISH. *Right:* Histograms of Dist. from DNA FISH measurements, where the Dist. = a*(shortest distance from locus to CT envelope / shortest radius distance of this CT). a = 1, if the locus inside the CT; a = −1 if the locus outside the CT. **(h)**. *Left:* the relative distance distribution of loci to CT in the IGM model. *Right:* Dist. from IGM structures, where Dist. = a × (shortest distance from locus to CT envelope / shortest centroid-to-envelope distance of this CT). a = 1 if the locus is inside the CT; a = −1 if the locus is outside the CT. CT of each chromosome copy per structure is defined as the 3D convex hull of the top 30% cis-to-total ratio beads; distances are computed directly on the raw hull. Both homolog copies of each locus are pooled as independent samples across all 1000 structures.

### Folding variability inferred from pseudo-single cell 3D genome models

To independently assess genome folding variability in single cells using a complementary computational approach, we used the Integrative Genome Modeling (IGM) software^33^ to generate 1,000 pseudo-single-cell 3D genome structures from previously generated H1-hESC bulk Hi-C data^44^ (**Fig. 2a**). Genome structures were generated at 200 kb resolution, with each 200 kb interval represented as a sphere of 95 nm radius, such that the total chromatin volume matches the nuclear chromatin density. To determine cis or trans contacts from pseudo-single cell models, contacts between spheres were defined when their surfaces were within 237.5 nm (2.5 times the sphere radius).

We next asked whether model-derived measurements could recapitulate the genome folding variability observed directly in scHi-C. Because the cis-to-total contact ratio serves as a proxy for a locus’s position relative to its CT (**Fig. 1b**), the availability of explicit 3D genome structures allowed us to directly quantify the extent to which genomic regions are embedded within or looped out of their CT cores using a geometric measure. For each chromosome copy in each genome structure, we defined the CT core as the 30% chromatin bins with the highest cis-to-total contact ratios, with the 30% threshold selected based on agreement with DNA FISH measurements (Methods). The CT boundary was then defined by the convex hull enclosing these core bins. For each genomic bin, we then measured the signed distance to the core CT boundary (dist-CT), assigning positive values to bins embedded within the CT and negative values to bins looping outside of it (**Fig. 2a, right top**). Across the 1,000 structures, the mean signed distance represented the population-average position of each locus relative to its CT boundary, whereas its standard deviation (S.D.) quantified cell-to-cell variability in this positioning. This provided a model-based geometric measure of locus-level folding variability, complementary to the S.D.of the cis-to-total contact ratio measured directly in scHi-C. The IGM-derived folding variability significantly correlated with those obtained with scHi-C, with the stronger concordance observed for droplet-based scHi-C (Spearman’s r= 0.44) (**Fig. 2b, Extended Data Fig. 2a**). Consistent with scHi-C data, model-derived folding variability was highly locus-specific and fluctuated substantially along the genome, with highly variable regions preferentially located in gene-dense A compartment domains, whereas gene-poor B compartment regions exhibited greater folding consistency (**Fig. 2b-c**).

### Folding variability is associated with heterogeneous subnuclear positioning

The 3D genome models allowed us to relate genome-folding variability, quantified as the standard deviation of the distance to the CT boundary (S.D. of dist-to-CT), to subnuclear localization, measured by the mean radial position (RAD) of each genomic region within the nucleus. More consistently folded regions were preferentially localized towards the nuclear periphery, whereas more variably folded regions were enriched in the nuclear interior (**Extended Data Fig. 2b the left panel**). Moreover, the standard deviation of a region’s radial position correlates with the genome folding variability (**Extended Data Fig. 2b the right panel)**. These results suggest that folding variability is coupled to broader positional plasticity within the nucleus, with variably folded regions exploring a wider range of both chromosome-territory and radial nuclear positions across cells. We therefore extended this analysis beyond radial position to examine chromatin localization relative to nuclear landmarks. To this end we integrated TSA-seq measurements for the nuclear speckle marker SON and the nucleolus marker POL1RE to predict the positions of nuclear speckles and nucleoli within each pseudo-single-cell model, following our previously established approach^47,48^. Together with the nuclear envelope, these landmarks allowed us to assign each genomic region in each pseudo-single cell model to a defined “nuclear locale” based on its relative proximity to the nuclear envelope, nuclear speckles, or nucleoli^49^. Distance thresholds were selected to maximize agreement with SON TSA-seq and POLIRE-TSA-seq data (Methods). Genomic regions not assigned to any of these landmarks were further classified according to their pseudo-single cell compartment identity as non-landmark active (NL-act) or non-landmark repressive (NL-rep) (Methods) (**Fig. 2a, Extended Data Fig. 2c**).

Because the same genomic region may occupy different locales across single cells, we characterized its variation in nuclear locations in the cell population using a locale-occupancy profile, defined as the fraction of instances a locus is assigned to each locale in the cell population (**Extended Data Fig. 2c lower panels)**. To quantify the cell-to-cell variability of a genomic region’s nuclear location, we then calculated the entropy of its locale occupancy profile. Low locale entropy indicates a consistent preference for a specific nuclear locale, whereas high entropy reflects a more heterogeneous distribution across locales in different cells and, consequently, greater positional variability across cells (Methods) (**Extended Data Fig. 2c**).

Genomic regions dominantly localized in the nuclear envelope or speckle locales possessed the lowest locale entropy, indicating comparatively consistent positioning at these landmarks across cells, whereas non-landmark-active and repressive regions showed substantially higher local entropy (**Fig. 2d**). Notably, speckle-associated regions nevertheless displayed a broader distribution of locale entropy and greater folding variability (S.D. of dist-to-CT) than envelope-associated regions. Thus envelope-associated regions appear more spatially constrained in both their folding and spatial positioning across cells. In contrast, regions predominantly assigned to non-landmark active (NL-act) or repressive (NL-rep) locales exhibited the highest locale entropy together with elevated folding variability, highlighting their inherent spatial uncertainty (**Fig. 2d-e**).

The relationship between CT-based folding variability and cell-to-cell variability in subnuclear positioning differed markedly among nuclear locales. For genomic regions dominantly found at the envelope, the locale entropy positively correlated with CT-based folding variability (ρ(ENV)=0.42), indicating that loci with less consistent envelope localization also varied more in their degree of CT embedding or looping out (S.D dist-to-CT) (**Extended Data Fig 2c,f**). In contrast, regions dominantly assigned to the non-landmark-active (NL-act) and repressive (NL-rep) locales showed the highest overall locale entropy but negative correlations with CT-based folding variability (ρ(NL-active)= −0.41 and ρ(NL-active)= −0.17). Thus, loci that sample a broader range of nuclear locales did not necessarily vary more in how far they extended beyond their chromosome territories. Speckle-associated regions showed relatively low locale entropy but no significant correlation with CT-based folding variability, suggesting that loci can maintain a consistent speckle-associated environment while varying substantially in the extent to which they can loop out from their chromosome territory. Together, these locale-specific relationships reveal distinct modes of coupling between CT-based folding and positional variability: subnuclear positioning and CT-based folding dynamics are positively coupled at the nuclear periphery, inversely related in non-landmark regions, and largely uncoupled near nuclear speckles.

### Detection of variable positioning of loci by microscopy

We validated folding variability by imaging. We selected eight loci on chromosome 2 and 11 that displayed a range of variabilities in cis-to-total contacts based on scHi-C data (**Extended Data Fig. 2f**). We examined their relative position within the chromosome territory (CT) using DNA FISH, by (i) measuring the distances from each locus to the centroid and to envelope of its corresponding CT; (ii) comparing the locus– centroid distance to the shortest centroid-to-boundary distances to determine whether the locus is positioned toward the interior, or exterior of the CT; and (iii) normalizing the locus–centroid distance by the shortest centroid-to-boundary distance of that territory, assigning positive values to loci located inside the CT and negative values to loci positioned outside the CT (**Fig. 2g**). Consistent with our scHi-C-based annotation, the variably folded regions displayed a broader distribution across cells - frequently positioned either inside or outside of the CT envelopes, while the consistently folded regions with lower variability were more frequently confined within the CT (**Fig. 2g, Extended Data Fig. 2g**). The spatial localization patterns measured by DNA FISH across all loci were also consistent with those inferred from the IGM models analyzed in the same way as illustrated in **Fig. 2f** (see Methods for details). Variably folded regions displayed a greater propensity to loop out of their chromosome territory and engage in interchromosomal interactions (**Fig 2h, Extended Data Fig. 2h**).

Together, these analyses demonstrate that genome-wide folding variability can be robustly annotated from scHi-C data and by 3D modeling of pseudo-single cells derived from bulk Hi-C data, and validated by DNA FISH. Variably folded regions, as defined here based on the S.D. of dis-to-CT are preferentially associated with transcriptionally active, gene-rich domains in the A compartment located away from the periphery.

### Cell-to-cell variability of genome folding changes as cells differentiate

To investigate the functional relevance of folding variability in a defined biological context, we measured cell-to-cell variability of genome folding across a well-characterized embryonic differentiation trajectory^44,50^. This trajectory spans five sequential stages, from human embryonic stem cells (H1-hESC) through definitive endoderm (DE), hepatoblasts (HB), immature hepatocyte-like cells (iHLC) and hepatocyte-like cells (HLC) (**Fig. 3a**). To ensure that our estimates of folding variability were not confounded by cellular heterogeneity, we confirmed the homogeneity of cell populations at each stage using both immunofluorescence (IF) staining and single-cell RNA-seq (**Fig. 3b, Extended Data Fig. 3a-c**).

**Figure 3:**
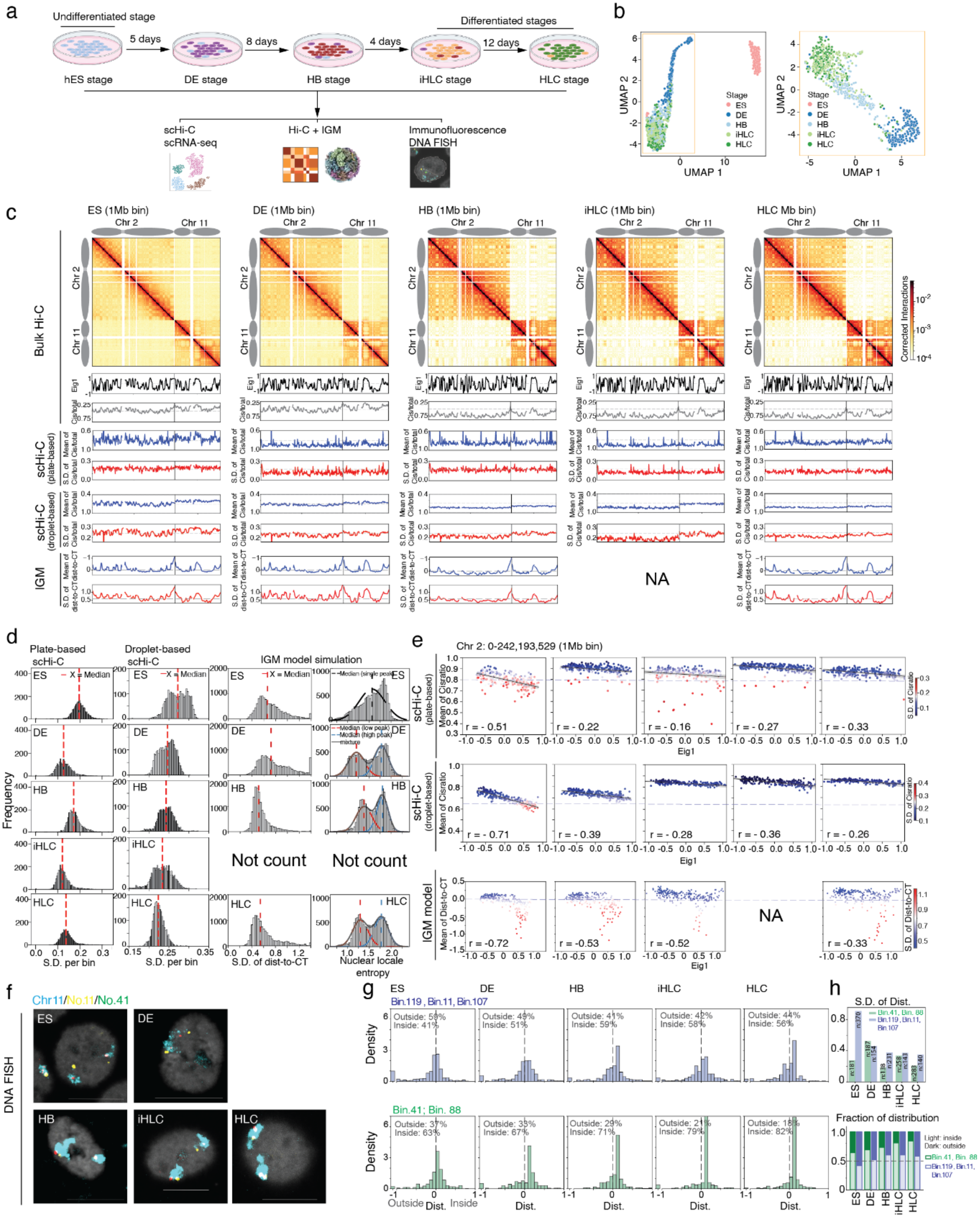
Genome folding variability progressively stabilizes during differentiation from pluripotent to lineage-specific states. **(a)** Experimental workflow integrating scHi-C, scRNA-seq, IGM modeling, immunofluorescence, and DNA FISH across five endoderm differentiation stages (ES, DE, HB, iHLC, HLC). **(b)** Uniform Manifold Approximation and Projection (UMAP) projections of scRNA-seq transcriptional trajectories for all five stages (left) and with ES omitted (right). **(c)** Chromosome-scale genome folding dynamics. Top: Bulk Hi-C contact heatmaps (Chr 2 and 11), eigenvector tracks (Eig1), and cis-to-total contact ratios. Middle: Plate/droplet scHi-C profiles showing mean cis-to-total ratio (blue, territory strength) and S.D. of cis-to-total contact ratio tracks (red, folding variability). (Plate scHi-C uses ES batch 1). Bottom: IGM profiles showing mean dist-to-CT (blue, territory strength) and S.D. of dist-to-CT (red, folding variability) tracks. **(d)** Genome folding variability and nuclear spatial positioning. Histograms of cis-to-total ratio S.D. (left/middle), dist-to-CT S.D. and nuclear locale entropy per genomic bin (right) across platforms and models. (Plate scHi-C uses ES batch 1). The red dashed line marks the median S.D. at each stage in the cis-to-total ratio S.D. plots. Histograms in the last column show the per-region nuclear location entropy distribution across five stages, computed over autosomal 200-kb regions (centromeric regions excluded). For each stage, the distribution was modeled with a Gaussian mixture: ES was fit with a single Gaussian, whereas the other stages were fit with a two-component mixture. Fitted component densities (dashed lines; red, lower-entropy peak; blue, higher-entropy peak) and their sum (grey) are overlaid and scaled to counts; dotted vertical lines mark each component mean. The emergence of a distinct lower-entropy population during differentiation reflects the progressive partitioning of the genome into stably positioned nuclear-landmark–associated regions. **(e)** Compartment vs. territory strength. Scatter plots of mean cis-to-total ratio (plate/droplet based scHi-C) or mean dist-to-CT (IGM) vs. Eig1 along Chr 2 across stages, colored by S.D. (blue = low variability, red = high variability). (Plate scHi-C uses batch 1 of 5 stages dataset and filtered contacts distance > 0.3kb; droplet Hi-C uses filtered contacts distance > 0.4kb of 5 stages dataset). **(f)** DNA FISH validation. Representative projection images of Chr 11 loci (yellow: 11–12 Mb; green: 41–41.5 Mb) relative to the chromosome territory (cyan) across differentiation. Scale bar: 10 μm. **(g)** Locus positioning dynamics. Histograms of normalized envelope distances (Dist) for variable vs. consistent Chr 11 loci relative to the territory edge (positive = inside, negative = outside). Histograms of Dist. from DNA FISH measurements, where the Dist. = a*(shortest distance from locus to CT envelope / shortest radius distance of this CT). a = 1, if the locus inside the CT; a = −1 if the locus outside the CT. **(h)** Quantitative analysis of distribution and localization changes for variably and consistently positioned loci relative to chromosome territories (CTs) across ES cells differentiation. Blue and green bars represent relatively variable and consistent loci, respectively. Top: Standard deviation (S.D.) of locus distances. Bottom: Fraction of loci located inside (light shades) versus outside (dark shades) of the CT across stages.

To systematically capture genome conformational changes during differentiation, we used previously generated bulk Hi-C data^44^ (asynchronous cells) and single-cell Hi-C data generated here (plate-based and droplet-based) across all five stages, where only interphase cells were retained for downstream scHi-C analysis (**Extended Data Fig. 4a**). The bulk Hi-C data were used to generate pseudo-single-cell 3D structures for four cell stages except for iHLC, which had lower sequencing depth and not enough inter-chromosomal contacts to build accurate models. In bulk Hi-C contact maps, we observed a global strengthening of the intra-chromosomal plaid pattern representing increased compartmentalization strength, and a marked decrease in inter-chromosomal interactions from ES to differentiated stages, reflecting the establishment of more defined chromosome territories during differentiation^44^ (**Fig. 3c, Extended Data Fig. 4b**). Population-level analysis of changes in genome organization detected with bulk Hi-C during this differentiation process are described in Huo et al^44^.

Single-cell Hi-C and IGM analyses revealed a progressive decrease in the cell-to-cell variability of both cis-to-total contact-based and CT-based folding measures and an increase in the average per-bin cis-to-total contact ratio in scHi-C and dist-to-CT in the models across the differentiation trajectory. This decrease in variability was genome-wide rather than confined to specific regions (**Fig. 3c, d, e**). However, regional heterogeneity persisted, with some distinct loci retaining higher variability than others (**Fig. 3c, Fig. 3e**, see below). Further, although the entire genome becomes more consistently folded as cells differentiate, in all cell stages loci in the A compartment remain more variably folded than loci in the B compartment (**Fig. 3c, 3e**). This difference persisted after mean-correction (resid_SD; see Methods) and removal of technical noise: A compartment loci retained significantly higher resid_SD than compartment B loci across all differentiation stages (**Extended Data Fig. 4c, supplemental note 3**), while genome-wide cis-to-total contact-based and CT-based folding variability decreased as cells differentiate (**Supplemental Note 3**). Notably, the locale entropy distribution became increasingly bimodal with progressive differentiation, indicating that a subset of genomic regions progressively adopt a more consistent subnuclear localization across cells. As shown below, these regions predominantly correspond to loci that become preferentially localized at the nuclear envelope (**Fig. 3d**).

To validate our scHi-C folding variability annotations during differentiation, we again used DNA FISH to track the nuclear positions of the same 5 selected loci on Chr. 11 shown in **Fig. 2g** as cells differentiate from ES cells to HLCs. Notably, the relative localization of these loci with respect to their CT from imaging closely reflected their folding variability changes from scHi-C and IGM pseudo single cell models (**Fig. 3f-h, Extended Data Fig. 4d**). Overall, the relative differences between more consistently (bins 41, 88) and more variably folded regions (bins 119,11,107) were maintained throughout differentiation (**Extended Data Fig. 4d)**. At the same time, genomic regions became globally less variable, as measured by both contact-based folding variability (S.D. of cis-to-ratio) and CT-based folding variability (S.D. of dist-to-CT) (**Extended Data Fig. 4d, Fig. 3h top panel)**. In parallel, loci became increasingly embedded within their chromosome territories, as reflected by increased mean cis-to-ratio and mean dist-to-CT values (**Extended Data Fig. 4d)**. This progressive increase in territorial embedding is consistent with our independent imaging, which showed an increased frequency of loci residing within their chromosome territories during differentiation (**Fig. 3g, and 3h bottom panel)**.

Given the high folding variability and weak compartmentalization observed in ES cells, we next investigated the underlying potential causes of this reduced compartmental segregation. Specifically, we considered three possible contributing factors: First, low structural similarities across single cells, whereby cell-to-cell differences in the epigenome state give rise to heterogeneous compartment profiles that weaken the population-average signal; Second: a high proportion of cells in S phase within the ES population; and third, intrinsically weaker compartmental segregation in individual cells due to reduced preferential interaction strength among regions of the same compartment. To first evaluate the impact of cell cycle phase, we sorted ES cells into G1, S and G2 populations and performed bulk Hi-C on each of them (**Extended Data Fig. 4e**). The Hi-C contact maps showed that G1-phase cells exhibited weaker compartmentalization than S-phase cells ((**Extended Data Fig. 4f, g**), thereby excluding the possibility that an overrepresentation of S-phase cells explains the decreased compartmentalization observed in bulk Hi-C data (**Fig. 3c**). Next, we examined the strength of compartmental segregation at the single-cell level using immunofluorescence staining of active and inactive chromatin marks - H3K27ac and H3K9me3, respectively-across the differentiation stages (**Extended Data Fig. 4h**), as described previously^51^. In ES cells, we observed greater overlap between H3K9me3 and H3K27ac signals, suggesting weaker spatial separation between active and inactive chromatin compartments. Furthermore, as cells progressed along the differentiation trajectory, H3K9me3 signals shifted from the nuclear interior to the periphery, while the H3K27ac signal fraction showed the opposite trend ((**Extended Data Fig. 4h right panels**). Taken together with the elevated folding variability in ES cells, these findings suggest that weak compartmental segregation within individual cells contributes to the weaker compartmentalization observed in ES population-based Hi-C data, possibly due to lower affinities between chromatin domains of similar type, which has been detected by liquid chromatin Hi-C in H1-hESCs^44^.

### Changes in chromosome territoriality correlate with changes in stage-specific folding variability

To explore the mechanisms underlying the progressive decrease in cell-to-cell genome-folding variability during differentiation, we examined how changes in nuclear morphology and chromosome organization contribute to this trend. First, we used imaging to measure the dimensions of nuclei in ES cells and in their differentiated progeny (**Extended Data Table 1**). We find that nuclear shape changed progressively from a nearly spherical morphology in pluripotent ES cells to a more flattened and polarized morphology in differentiated stages. These measured dimensions were incorporated as geometric constraints when generating ensembles of pseudo-single-cell 3D genome structures with IGM (**Fig. 4a**).

**Figure 4:**
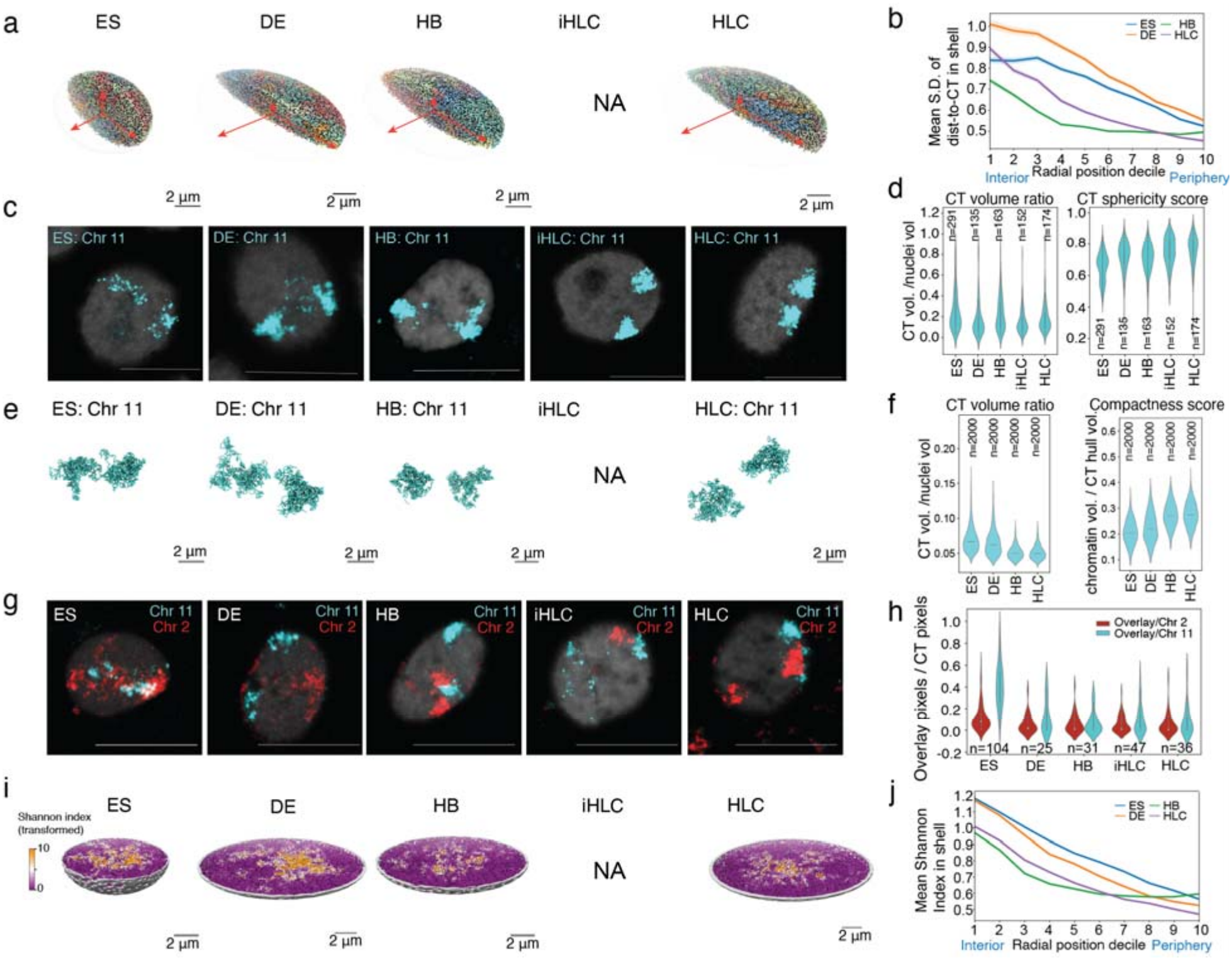
Chromosome territory compaction, and peripheral localization contribute to folding variability decreasing during differentiation. **(a)** Representative nuclear shapes reconstructed from models across four stages. Red arrows indicate the three axes of the coordination. **(b)** Genome-wide relationship between cis-ratio variability and radial position across the five stages. For each cell type, genomic bins were assigned to 10 radial shells of approximately equal bin count, defined by deciles of the rank-transformed normalized radial position (shell 1, lowest radial value; shell 10, highest, chrX, chrY, and centromeric bins excluded). The x-axis indicates the radial shell, and the y-axis shows the per-shell mean of S.D. of dist-to-CT. The shaded bands denote the 95% confidence interval (mean ± 1.96 × SEM); shells containing fewer than 200 bins were excluded. (**c)** Representative chromosome 11 territory structures visualized by DNA FISH staining across five stages. Scale bar: 10μm. **(d)** Violin plots showing quantified chromosome territory properties for chromosome 11 measured from DNA FISH (c). Left: ratio of CT volume to total nuclear volume; right: sphericity scores of the CT in DNA FISH. **(e)** Representative chromosome 11 territory structures visualized in models across four stages. Scale bar: 2μm. **(f)** Violin plots showing quantified chromosome territory properties for chromosome 11 measured from 3D models. Left: ratio of CT volume to total nuclear volume, where CT volume is defined as the volume of the convex hull enclosing all chromatin beads belonging to the same chromosome copy; right: Compactness score, defined as the summed volume of all chromatin beads in the same chromosome copy divided by the corresponding CT volume. **(g)** Representative picture from maximum-intensity projections of DNA FISH images for chromosome 11 (cyan) and chromosome 2 (red) across the differentiation stages. Scale bar: 10μm. **(h)** Violin plot showing the fraction of overlapping pixels between chromosome 2 and chromosome 11 territories in DNA FISH across five stages. **(i)** Representative half-spherical visualization of chromosome structure from model across four stages of differentiation. Chromatin regions are colored according to their chromosome-intermingling entropy (Shannon index score S). For visualization, values are transformed by a power-law (S6) prior to mapping onto a purple-to-orange color scale, with purple and orange indicating low and high intermingling entropy, respectively. **(j)** Genome-wide relationship between chromosome-intermingling entropy and radial position across the five differentiation stages. For each cell type, genomic bins were assigned to 10 radial shells of approximately equal bin count, defined by deciles of the rank-transformed normalized radial position (shell 1, lowest radial value; shell 10, highest, chrX, chrY, and centromeric bins excluded). The x-axis indicates the radial shell, and the y-axis shows the per-shell mean Shannon index score. The shaded bands denote the 95% confidence interval (mean ± 1.96 × SEM); shells containing fewer than 200 bins were excluded.

We then quantified CT-based folding variability (S.D. of dist-to-CT) together with locale entropy as a function of radial subnuclear position across all four cell states (**Fig. 4b**). This analysis revealed three major patterns of folding variability dynamics: (i) Across all four cell states, loci in the nuclear interior generally displayed higher CT-based folding variability than those near the periphery. (ii) Loci in cells of earliest differentiation stages (ES and DE cells) displayed higher folding variability across almost all radial shells compared to later stages (HB, HLC). However, at later differentiation stages (HB, HLC), folding variability decreased most strongly in the outer radial shells, indicating that differentiation-associated stabilization is most pronounced toward the nuclear periphery. (iii) In ES cells, folding variability peaks at intermediate radial positions between the nuclear center and periphery. In differentiated cells, however, the peak in folding variability shifted to the nuclear center. We characterize these highly variable regions in greater detail below.

Both Hi-C and scHi-C data showed a marked decrease in trans contacts and a corresponding increase in cis contacts as ES cells differentiate into HLCs (**Fig. 3c, 3e**). This shift suggests that chromosomes become more territorial as cells differentiate. To test this directly, we performed chromosome painting to co-stain cells with probes for chromosomes 2 and 11. We find that in ES cells chromosome territories are relatively large and diffuse and often appear fragmented into multiple regions (**Fig. 4c, g**). By contrast, in DE cells and subsequent differentiated states, chromosome territories become more compact and stain more brightly. This was confirmed by quantifying the chromosome territory volume normalized for nuclear volume, which shows a decrease during differentiation (**Fig. 4d, Extended Data Fig. 5a-b**), together with an increase in sphericity of the territory (**Fig. 4d, Extended Data Fig. 5a-b**, right panel, see Methods). IGM-derived pseudo-single-cell structures showed the same trend across the four cell states: chromosomes in ES cells are more extended, and become progressively more compact from the DE stage onwards (**Fig. 4e, f, Extended Data Fig. 5a-b**). Increased territoriality is expected to reduce the extent of interchromosomal mixing. We therefore quantified the overlap between chromosomes 2 and 11 using chromosome painting. Consistent with this expectation, the two chromosomes showed substantially greater overlap in ES cells than in DE cells and subsequent differentiated states (**Fig. 4g, h**).

The IGM derived pseudo-single-cell structures allowed us to extend this analysis to all chromosome pairs, and to determine where in the nucleus inter-chromosomal intermingling is most pronounced (**Fig. 4i, j**). To quantify inter-chromosomal intermingling, we computed a chromosome-intermingling entropy that measures the diversity of chromosome identities within the local neighborhood surrounding each genomic region (**Methods**). This analysis revealed a progressive reduction in inter-chromosomal intermingling during differentiation (**Fig. 4j**), consistent with the chromosome-painting experiments. Across all cell states, chromosomal intermingling was highest in the nuclear interior and progressively decreased towards the nuclear periphery.

These results confirm that chromosomes are extensively intermingled in ES cells and become progressively more territorial during differentiation. Furthermore, we note that chromosome folding variability, expressed as S.D. of dist-to-CT, follows the same decreasing trend where local inter-chromosomal intermingling decreased as the nuclear radial localization increased (**Fig 4b, j**), suggesting that regions occupying compositionally mixed nuclear environments, i.e., containing loci from multiple chromosomes, show greater cell-to-cell variation in folding.

Together, these results show that the decrease in genome folding variability during differentiation is accompanied by coordinated changes in (i) nuclear morphology, (ii) chromosome territory compaction, and (iii) reduced inter-chromosomal intermingling. As shown below, these changes are also accompanied by a progressive relocalization of chromatin towards the nuclear periphery.

### Folding variability is driven in part by the balance of peripheral tethering and access to interchromosomal space

To determine which experimental restraints encode the observed locus-specific patterns of folding variability, we systematically generated IGM ensembles using individual or combined subsets of the input data (**Supplementary Note 4, Supplemental Figure 5a**). Models restrained only by Hi-C derived trans interactions already closely reproduced the locus-specific pattern of CT-based folding variability observed in the fully restrained reference model (r = 0.91), although their overall variability was slightly lower (median S.D., 0.58) (**Supplemental Fig. 5a the left chart, Supplemental Fig. 5b, the third left model**). This contrasts with cis-only models and is consistent with trans restraints directly constraining loci in interchromosomal space and therefore specifying to what extent specific regions loop out beyond their chromosome territories. Strikingly, models restrained only by LaminB-DamID-derived probabilities of nuclear envelope association also closely reproduced both the overall magnitude of folding variability (median S.D., 0.64), and the locus-specific variability pattern observed in the reference model (r = 0.86), despite the absence of any chromatin-chromatin interaction data (**Supplemental Fig. 5a the left chart, Supplemental Fig. 5b, the second left model**). Combining trans and LaminB-DamID restraints provided little additional improvement, suggesting that the two data types encode largely overlapping information about locus-specific CT embedding. Together, these results link folding variability to the balance between peripheral tethering and access to interchromosomal space and suggest that changes in lamina-association propensity can reshape folding variability across the genome.

### Regions gaining nuclear landmark association show reduced folding variability during differentiation

We used nuclear locale assignments to track how chromatin regions change their locations relative to nuclear landmarks during differentiation. Differentiation was accompanied by marked changes in the relative chromatin distribution across nuclear locales, most pronounced at the nuclear speckle and nuclear envelope locales. Upon exit from pluripotency, the fraction of chromatin regions assigned to the nuclear envelope locale increased from 27.2% in H1-hESC to more than 34.7% in HLCs (**Fig. 5c**), whereas the fraction assigned to the speckle locale decreased from 18.0% to 14.6%. Locale occupancy also decreased modestly in the non-landmark-active locale while the fraction of chromatin in non-landmark repressive locales increased (**Fig. 5c**). Overall, the fraction of chromatin in repressive locales (envelop, non-landmark-repressive and nucleoli) increased from 51.0% in H1-hESC to 70.8% in HLC cells, while active chromatin (non-landmark-active, speckle) decreased (**Fig. 5c**).

**Fig 5.**
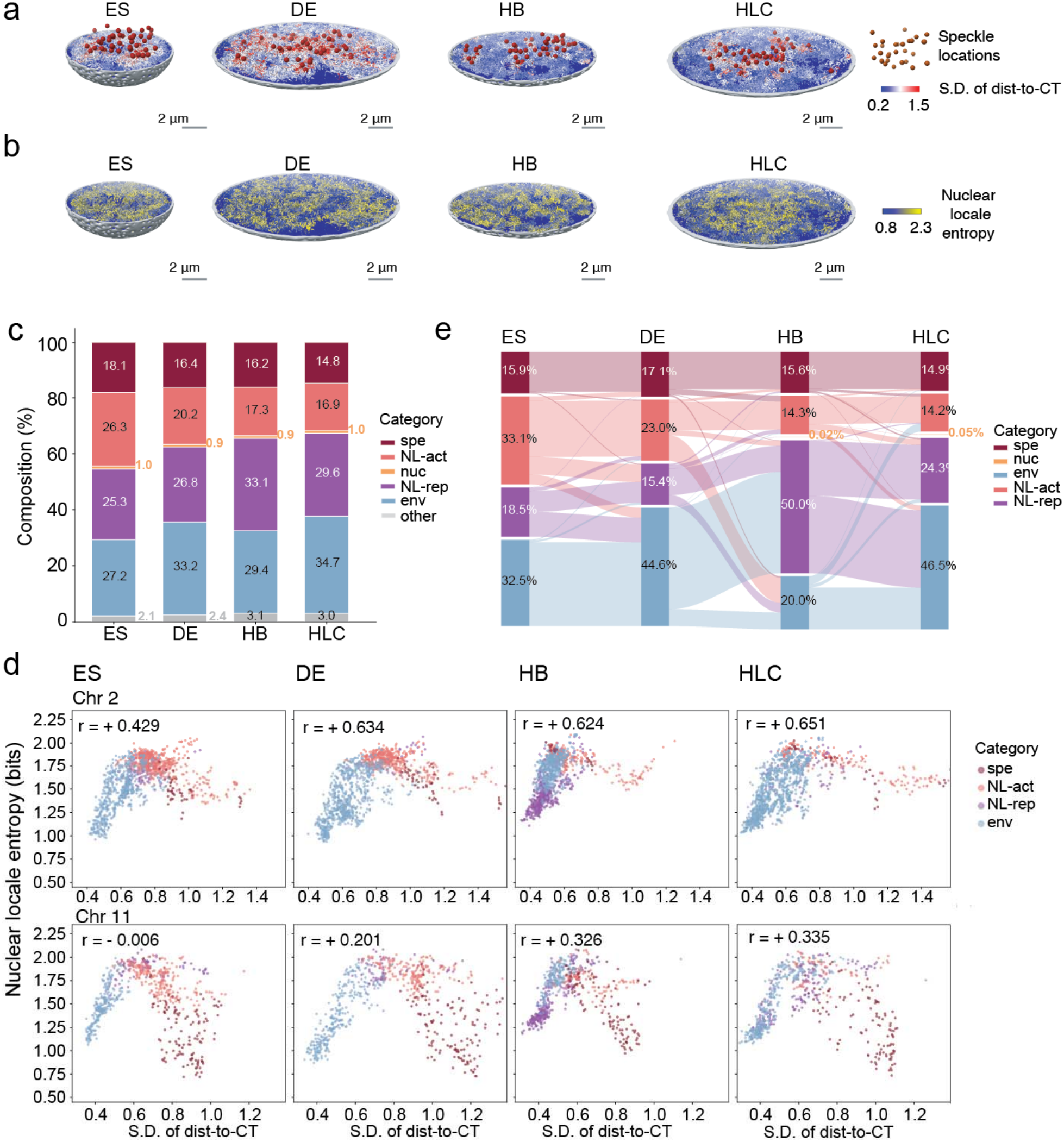
Relocation of genomic regions toward nuclear landmarks contributes to folding consistency during differentiation. **(a)** Representative hemispherical average feature map of S.D. of dist.-to-CT in IGM models across four differentiation stages. Each 200-kb chromatin bead is colored by the S.D. of dist-to-CT (red = high variability, blue = low variability). Dark red dots mark the positions of speckle cluster centers. **(b)** Representative hemispherical single nuclear feature map of nuclear location entropy on IGM structure across five stages. Each 200-kb chromatin bead is colored by its nuclear location entropy over the five nuclear locale classes; both homolog copies of a haploid bin share the same value. Blue represents low entropy and yellow represents high entropy. **(c)** Nuclear locale composition across four differentiation stages. For each stage, locale assignments were pooled across all 1,000 structures and both homologues locus allele copies of all autosomal, non-centromeric 200-kb genomic regions. Based on their single-cell subnuclear locations or ABratio, each locus instance was classified into into one of five primary nuclear locale categories: speckle-associated (spe), nucleolus-associated (nuc), envelope-associated (env) nonlandmark-active (NL-act) or non-landmark repressive (NL-rep) (Methods). **(d)** Genome-wide correlation scatter plot of S.D. of dist-to-CT (x-axis) versus nuclear location entropy (y-axis) in each of the four differentiation stages. Each dot is one genomic region, colored by its dominant locale, defined as the argmax over the five normalized nuclear-landmark association probabilities determined from the IGM-derived 3D genome structures; Pearson r and Spearman ρ are calculated between S.D. of dist-to-CT and entropy. Regions were restricted to autosomes and non-centromeric bins. Top: chr2; bottom: chr11. **(e)** Sankey-style representation of the dominant (canonical) nuclear locale distributions across differentiation stages. For each genomic region at each stage, the dominant locale was defined as the locale with the highest frequency in its locale-occupancy profile. Each of the four columns represents one cell stage. Nodes within each column indicate the fraction of genomic regions assigned to each dominant locale (spe, nuc, env, NL-act, NL-rep) and ribbons visualize the changes in these fractions between consecutive differentiation stages.

We used locale entropy as a measure of cell-to-cell variability in subnuclear positioning, with higher entropy indicating that a genomic region occupies a broader range of nuclear locales across cells. During differentiation, locale entropy developed a pronounced bimodal distribution (**Fig. 3d**). Some regions retained high local entropy whereas an increasing subpopulation with substantially reduced locale entropy emerged, composed predominantly of envelope- and non-landmark-repressive regions. The reduction in locale entropy within this subpopulation was not driven by decreasing entropy within individual locale classes. Regions preferentially associated with repressive locales, including the envelope, non-landmark-repressive locales, maintained similar entropy across differentiation, whereas speckle- and NL-active regions showed a modest increase in entropy (**Extended Data Fig. 6d**). Thus, the emergence of a low-entropy subpopulation in differentiated cells (DE, HB, HLC) primarily reflects a redistribution of genomic regions toward intrinsically lower-entropy locales, particularly the nuclear envelope, rather than a general reduction in entropy within each locale class.

Indeed, exit from pluripotency was accompanied by a major redistribution of dominant nuclear locales, defined as the locale occupied most frequently by a locus across cells. A substantial fraction of genomic regions that were predominantly associated with the non-landmark active locale in ES cells were most frequently observed in the non-landmark repressive or envelope locale in DE cells. Moreover, a substantial fraction of regions most frequently associated with the non-landmark-repressive locale in ES cells shifted to the envelope locale, contributing to the emergence of a lower-entropy population because envelope-associated regions generally exhibit lower locale entropy in DE and HLC cells (**Fig. 5d, e, Extended Data Fig. 6e**). Together, these results imply that a fraction of genomic regions with variable nuclear localization in ES cells gain positional consistency in differentiated cells primarily through stable association with repressive nuclear landmarks, such as the envelope.

Although the combined fraction of regions predominantly assigned to repressive locales (ENV and NL-rep) increased progressively during differentiation, their distribution between these two environments was not strictly monotonic. At the HB stage, dominant assignments shifted transiently from the envelope toward NL-rep, accompanied by increased locale entropy specifically among envelope-associated regions. Because the dominant locale is determined by the highest occupancy frequency, higher entropy makes assignments more sensitive to modest shifts between competing locales. Thus, the HB pattern likely reflects increased cell-to-cell sharing of repressive chromatin between peripheral and interior repressive environments rather than a loss of envelope association. The enlarged nucleoli observed in HB cells^44^ are consistent with such increased redistribution between lamina-associated and interior repressive compartments.

To evaluate whether differentiation-associated changes in folding variability are coupled to changes in subnuclear positioning, we compared folding variability against locale entropy across all differentiation stages (**Fig. 5abd, Extended Data Fig. 6c**). The correlation between these metrics strengthened progressively during differentiation, increasing from Spearman’s (r= 0.156) in ES cells to (r=0.316) in DE cells, (r=0.346) in HB cells and (r = 0.361) in HLCs (**Fig. 5d, Extended Data Fig. 6c, e**). The locale-specific relationships observed in ES cells were largely preserved during differentiation, although their absolute value increased. For example, with exception of HB cells, regions in the envelope locale showed an increasingly strong positive correlation, from (r=0.41) in ES cells, to (r=0.44) in DE cells and (r=0.69) in HLC cells (**Fig. 5d, Extended Data Fig. 6e**). In contrast, non-landmark active regions retained a negative correlation, although it weakened modestly from (r=-0.41) in ES cells to (r=-0.35) in DE cells, (r=- 0.24) in HB cells and (r=-0.38) in HLC cells.

Similar to ES cells, CT-based folding variability in differentiated cells is concentrated mostly in transcriptionally active interior chromatin, including near speckles (**Extended Data Fig, 6a-b**), where loci vary substantially in their distance to the chromosome territory boundary, while remaining consistently positioned near speckles with relatively low locale entropy. Among the top 10% most variably folded regions, the fraction with speckle as the dominant locale increased during differentiation, despite an overall decrease in speckle-dominant regions genome-wide (**Extended Data Fig. 6b**). This indicates that, after differentiation, folding variability is increasingly concentrated at active locales, in particular speckles.

### Folding variability correlates with expression heterogeneity

Given that gene expression can be variable between cells and across the genome, we examined how genome folding variability in pluripotent and differentiated cells is related to variability of gene expression. We used scRNA-seq data obtained for each of the 5 cell types, and then analyzed the relation to folding variability at two levels: (i) we quantified how gene expression patterns differ between single cells for each of the different cell types; (ii) within the same cell type, we determined how gene expression variability fluctuated across the genome in single cells.

We first examined the genome-wide expression pattern variability. ES cells displayed greater cell-to-cell transcriptional diversity compared to cells at more differentiated stages. As differentiation progressed, the number of genes expressed per cell markedly decreased, with only 10-15% of genes detected in fully differentiated cells compared to the starting ESC stage (**Extended Data Fig. 7a, b**), consistent with a previous report^52^. At the transcriptome-wide level, principal component 1 in PCA revealed that ES cells exhibited broader dispersion in gene expression profiles, whereas differentiation led to progressively more homogeneous expression patterns (**Fig. 6a, Extended Data Fig. 7c**).

**Fig 6.**
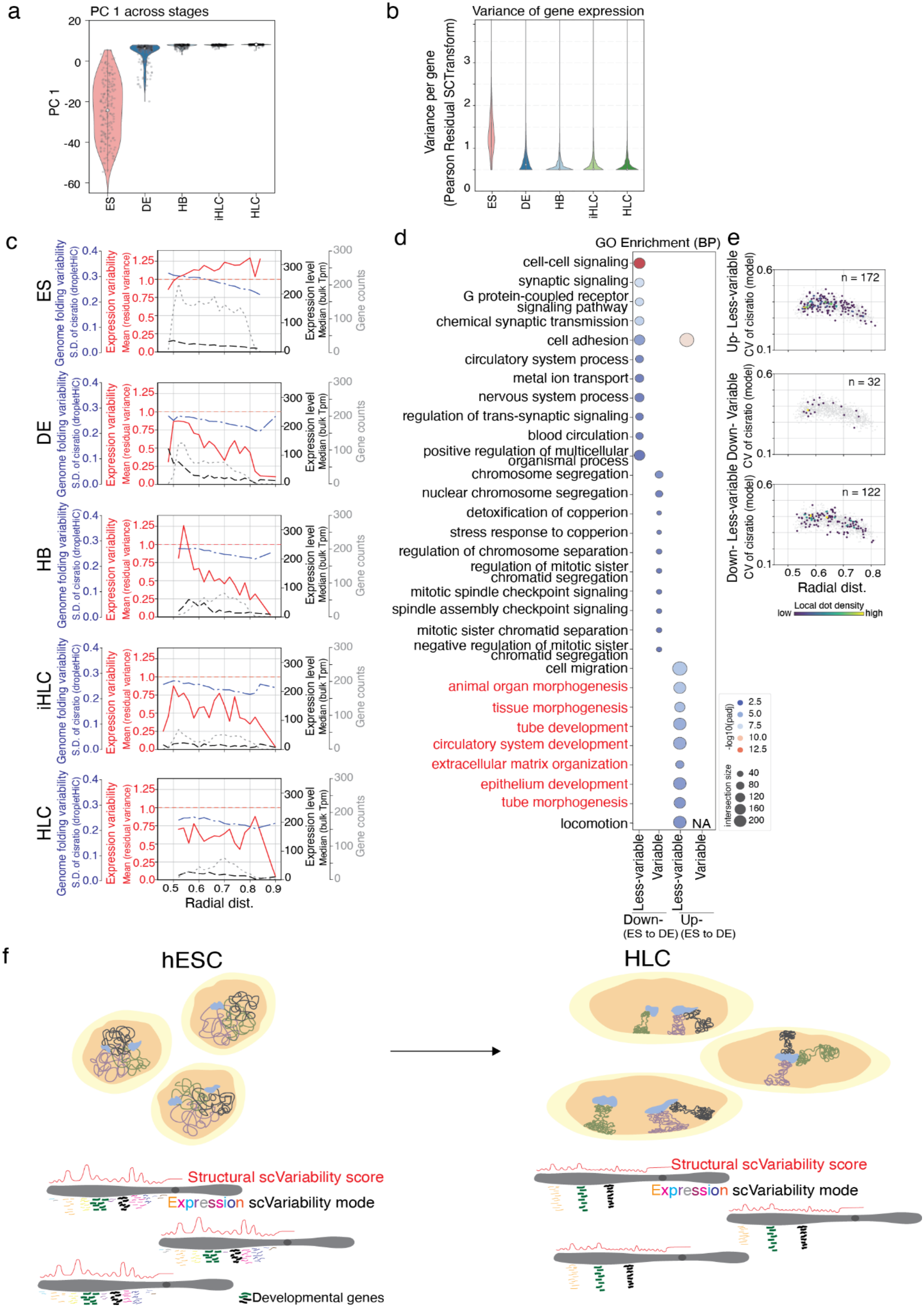
Folding variability underlies expression heterogeneity and genomic priming potential. **(a)** Violin plots showing the principal component 1 from scRNA-seq, illustrating transcriptional separation of cell states during early differentiation. **(b)** Violin plots showing the variance of Pearson residuals per gene across all cells within the same stage. For each gene, Pearson residuals represent standardized deviations of observed expression from the model-predicted mean, scaled by the expected variance. **(c)** Line plots show relationships between radial distance with structural variability (blue, aggregated structural variability score from models), gene expression variability (red, mean of gene’s residual variance per radial distance window), gene expression level (black, bulk Tpm) and gene counts (grey). **(d)** Bubble plots showing GO term enrichments across four gene categories: genes up-regulated or down-regulated from ES to DE; and genes variably expressed, or non-variably expressed in ES. Red text marks developmental gene-related GO terms. Enrichments were detected in only three of these four conditions. **(e)** Scatter plots showing the relationships between radial distance (distance of each genomic bin from nuclear center) and S.D. of cis-to-total contact ratios contact ratio per genomic bin (1Mb bins) in droplet Hi-C. Points corresponding to genes in GO terms identified in panel **e** are highlighted and colored by kernel density. **(f)** Schematic illustration summarizing structural sc-variability changes and associated with transcriptional changes across differentiation.

We correlated genome-wide fluctuation in folding variability with gene expression variability along the genome. To this end, we quantified transcriptional variation for each gene across single cells by calculating the variance of Pearson Residual, derived from regularized negative binomial regression. (**Extended Data Fig. 7d, e**, see Methods)^53^. The results showed overall genes in ES cells exhibited markedly higher residual variation scores than those in more differentiated stages, consistent with PCA analysis (**Fig. 6b**), indicating greater expression heterogeneity at the gene level in pluripotent cells.

Second, to integrate expression heterogeneity with genome folding and nuclear position, we filtered out genes with dropout ratio > 0.85 and TPM <1 (**Extended Data Fig. 7f, Methods**), and then for each genomic bin of 1Mb we calculated the mean of the residual variation for all genes contained in the bin. As described above (**Fig. 4**), we also calculated for each of these bins the folding variability (S.D. of cis-to-total contacts ratio in droplet scHi-C), and the average radial position across the 1,000 pseudo-single cells, for each of the 5 cell states. This allowed us to plot the average gene expression variation as a function of radial position and compare this to the folding variability (**Fig. 6c**).

A variance of the Pearson residual of gene expression larger than 1 indicates that the variation of expression of this gene is larger than expected from technical sampling noise. Notably, loci containing such truly variable expression (Mean of Pearson residual variance > 1) were almost exclusively found in ES cells, with negligible representation in the other four stages (**Fig 6c**). In ES cells high expression variability was observed throughout the nucleus, with a small set of speckle-associated regions located at the nuclear center showing relatively higher expression level and lower expression variability (**Fig 6c**). We note that there are also genes with low expression variability located throughout the nucleus (**Extended Data Fig. 7g**, see below). These observations show that ES cells retain a loose and globally variable genome organization, in which most genes are expressed more stochastically genome-wide across cells. In differentiated cells, in contrast, gene expression variability (red line in **Fig 6c**) is generally lower than the threshold for true biological variability across the nucleus and genome, with most fluctuations remaining within single-cell RNA-seq technical variance (Pearson residual variance < 1), except for a small number of highly expressed loci in the center of the nucleus, e.g., in DE cells. Overall, folding variability and gene expression variability are both lower in differentiated cells.

We next explored how genes with high or low expression variability in ES cells change expression during differentiation. First, we find that both types of genes can become up- or down-regulated as ES cells differentiate into DE cells (**Extended Data Fig. 7h**). Also, the difference in fold change in expression between high variable and low variable genes was modest (**Extended Data Fig. 7h**). We then performed Gene Ontology (GO) enrichment analysis to identify functional categories associated with high and low expression variability genes in ES cells that are up- or down-regulated during differentiation (**Fig. 6d**). Two observations stand out: First, highly variable genes in ES cells that are upregulated during differentiation do not show any remarkable GO term enrichments. Downregulated variable genes were enriched for mitosis and chromosomal segregation process (**Fig. 6d**), which is as expected. In contrast, genes with low gene expression variability in ES cells and that are up-regulated during differentiation are enriched in GO terms associated with development and morphogenesis. Genes that are differentially expressed and associated with the enriched GO terms in **Fig. 6d** are localized throughout the nucleus, and not preferably in the center near speckles (**Fig. 6e**). This includes the genes with low expression variability in ES cells that are up-regulated in DE cells and that are enriched in GO categories related to development.

Together the results show that ES cells and differentiated cells differ globally in chromosome folding variability and gene expression variability (**Fig. 6f**). Within the highly variable ES genome, some regions show relatively lower variability: First, genes near speckles are consistently positioned in speckle locale and relatively stably expressed. Second, a set of genes located throughout the nucleus are also relatively stably expressed and these are enriched in developmental genes that become up-regulated during differentiation.

## Discussion

Heterogeneity in chromosome folding is an expected property of genomes that emerges because of the nature of self-assembly of chromosome conformations driven by biophysical and molecular processes such as phase separation, loop extrusion, and tethering of loci to sub-nuclear structures and bodies^1,2^. Our combined genomic and modeling approach provide important new insights into the sources of genome folding heterogeneity by mapping variability along the length of the genome, throughout the 3D space of the nucleus, and during cell differentiation. Our study of chromosome folding heterogeneity of embryonic stem cells and their differentiated progeny identified two modes of chromosome folding variability: a highly heterogeneous state in embryonic stem cells, and a more consistently folded state in differentiated cells. We find that both states show fluctuations in the level of chromosome folding variability along the genome and throughout the nucleus, with loci not associated with structural landmarks such as the nuclear lamina being relatively more variable. These two states are related to gene expression variation between single cells, with stem cells displaying genome-wide expression variation, while differentiated cells expressing more restricted sets of genes and in a more consistent manner.

Previous studies have shown that ES cells have chromatin features that are distinct from differentiated cells. First, chromatin is more mobile as detected by life cell imaging^54–57^. This suggests that loci are less constrained, e.g., through weaker interactions with each other and with nuclear landmarks such as the nuclear lamina. Second, ES cells display so-called bivalent domains marked by both active (e.g., H3K4me3) and inactive (e.g., H3K27Me3) histone modifications^58–61^. Third, chromosome paints in mouse ES cells showed extensive intermingling of different chromosomes^62^, which is not typical for differentiated cells where chromosomes form distinct territories. Our imaging and Hi-C data are consistent with increased intermingling. In all, the ES cell genome appears less constrained, and this may lead to a larger variation in chromosome folding between cells.

This structural fluidity of the genome in ES cells is coupled with transcriptional activity of more genes than observed in differentiated cells, and larger expression heterogeneity between cells. It has been proposed that this state is important for pluripotency by producing a nuclear environment that is permissive for rapid, stochastic, and reversible transcriptional changes^63,64^. Such a state likely facilitates exploration of multiple regulatory configurations and rapid repositioning between transcriptionally active hubs and repressive compartments. These observations align with the concept that 3D genome plasticity provides a structural substrate for dynamic transcriptional changes^54,65,66^ and support the notion that spatial genome organization is both a marker and mechanism of pluripotency^67,68^.

Our scHi-C data and single cell 3D genome models show that while folding variability is high in general in ES cells, defined types of loci display lower variability in the types of locales they occupy. Interestingly, these are annotated as either speckle associated or peripheral lamina associated. Possibly, association with such nuclear landmarks for these loci is positioning these loci more consistently across single cells. As discussed below, as cells differentiate these associations become more frequent and consistent in the population. Also, despite widespread transcriptional heterogeneity in ES cells, some genes are less variably expressed across cells. These include highly expressed genes located in active regions of the genome that are speckle-associated, as well as some developmental regulators that become further up-regulated during the first step of differentiation. The relative expression consistency of these genes, together with their more consistent residence at specific sub-nuclear locales, may help preserve the core cellular state while maintaining the transcriptional potential required for subsequent lineage specification, especially during the closest developmental transitions.

As ES differentiate, we find that chromosome folding becomes more consistent: First, variability in locus positioning with respect to their chromosome’s territory is reduced. This is correlated with an increase in territoriality as detected by imaging, Hi-C, and IGM models. Chromosomes become less intermingled with each other. Finally, the locale entropy decreases, which indicates that loci are found in more similar locales across cells. While the overall level of folding heterogeneity decreases, the decreases are strongest for loci associated with nuclear landmarks, especially the nuclear lamina. Loci not associated with landmarks and located more centrally in the nucleus display relatively more variability as detected by locale entropy and locus positioning with respect to their territory. At the same time gene expression profiles become more restricted, with fewer genes expressed. The smaller set of genes show reduced cell-to-cell variation in expression.

While chromosome folding in differentiated cells becomes more consistent, it is important to emphasize that chromosome and nuclear organization is still stochastic. Our metrics for folding variability, S.D. of the cis-to-total contact ratio and locale entropy, measure specific aspects of variability, and do not represent all types of heterogeneity. For instance, a low locale entropy indicates that a given locus is consistently positioned at the lamina near other loci that are at the nuclear periphery. However, which specific sets of loci are found interacting with others or are close together at the lamina in each cell is still highly variable, as seen by imaging (e.g., Kind2013^69^; Misteli2020^1^). Similarly, while the S.D. of the cis-to-total interaction may be lower in differentiated cells, which loci along a chromosome are spatially interacting within a territory can still be highly variable. Here, we chose these two metrics because they represent known features of genome regulation (active genes often loop out of their territory), and nuclear locales are related to chromatin state and transcription, and because they can be extracted from scHi-C data and 3D models.

What determines the level of chromosome folding variability in ES cells and in differentiated cells? Our results show that association of loci with nuclear landmarks lowers locale entropy. This is true both in differentiated cells and in ES cells. In ES cells however, the association with landmarks appears much less consistent, especially for the nuclear periphery. This suggests that ES cells and differentiated cells differ in the molecular composition of either the landmarks or the chromatin or both. Consistent with this hypothesis, in a separate study we found that nuclear speckles change morphology as ES cells differentiated toward HLCs^44^. Also, during this differentiation process Lamin B is down regulated, and lamin A/C becomes highly expressed. Possibly, these changes increase the affinities of active and inactive loci with speckles and the nuclear lamina respectively, leading to more consistent positioning of these loci across the cell population. Similarly, our Hi-C and immunofluorescence data show that compartmentalization in ES cells is weaker and active and inactive domains are often co-localized while in differentiated cells they are more separate, consistent with earlier observations^44,70,71^. This suggests that the homotypic affinities between chromatin domains of the same type are weaker in ES cells than in differentiated cells, leading to less compartmentalized nucleus, and higher local entropy. The molecular determinants for compartmental chromatin interactions are poorly understood, but they must differ in ES and differentiated cells. Consistently, in a separate study we find that histone modifications change as cells differentiate, and especially H3K9Me2 levels increase during hepatocyte differentiation at a time when compartmentalization strength increases and peripheral chromatin becomes more consistently positioned^44^. Such modifications in histone modification may drive chromatin interactions either directly or through recruitment of complexes that bind and possibly bridge these modifications.

We conclude that while ES cells and differentiated cells employ common chromosome folding mechanisms such as association of loci with nuclear landmarks and compartmentalization of active and inactive chromatin, these processes are implemented in quantitatively different ways. In ES cells associations with nuclear landmarks and interactions between chromatin domains of similar type are all weaker. This results in a highly heterogeneous genome conformation in ES cells, and a more spatially deterministic, but still stochastic state in differentiated cells. Study of ES cell differentiation provides an experimental system to identify the molecular and biophysical determinants of these fundamental chromosome folding mechanisms.

## Methods

### Cell culture

#### HAP1 cells culture

HAP1 cells were cultured in RPMI-1640 medium supplemented with pyruvate (Gibco, 10569010) supplemented with 10% fetal bovine serum (Gibco, 16000044) and 1% penicillin-streptomycin (Gibco, 15140) at 37 °C in 5% CO_2_.

#### HAP1 mitotic synchronization time course

Prometaphase synchronization was performed as previously described^72^ with some modifications. HAP1 cells were seeded at 2,500/cm^2^ 16 hours before synchronization. Cells were treated with 0.5uM Palbociclib (Selleckchem, PD-0332991) in cell culture medium for 24 hours to arrest cells in G1 phase, and then cells were washed with 1XDPBS. Standard medium was added to release cells from the G1 arrest for 8 hours. Next, media was replaced with the same media containing 100ng/ml nocodazole (Sigma, M1404) for 9.5 hours to arrest cells in prometaphase. Mitotic cells were detected by Hoechst (Thermo Fisher, H1399) staining. Floating and attached cells were then harvested for single cell sorting and scHi-C.

#### H1-hESC cells culture

The H1-hESCs (H1, WiCell, WA01, lot no. WB35186) were cultured in mTeSR1 media (StemCell Technologies, 85850) under feeder-free conditions on Matrigel H1-hESC-qualified matrix (Corning, 354277)-coated plates at 37_°C and 5% CO2. H1-hESCs were fed daily with fresh mTeSR1 media and passaged every 4–5_days using ReLeSR reagent (StemCell Technologies, 05872). Cells were dissociated into single cells with TrypLE Express (Thermo Fisher, 12604013).

#### H1-hESC to HLC differentiation

The differentiation was performed using standard differentiation protocol in 4DN^50^. Briefly, cultured H1-hESCs were detached by TrypLE (Life Technologies) and 1 million cells /1 ml cell density was seeded into plate or well coated with GF(−) Matrigel (0.125 mg/ml). DE cells were induced using the STEMdiffTM Definitive Endoderm Kit (StemCell Technologies) kit. Briefly, cells were washed with DMEM/F12 on the second day, and then medium was changed with STEMdiff Definitive Endoderm Basal medium with supplements A (MR) and supplemental B (CJ) for day 2 and then supplement B (CJ) only for day 3 and day 4 (total 4 days). Matured DE can be detected with immunofluorescence by SOX17 (R&D Systems, AF1924) and FOXA2 (EMD Millipore, 07-633). For hepatic specification (hepatoblasts), DE were detached by TrypLE, and seeded at a density of 1.5 million cells/1 ml density into GF(−) Matrigel (0.125 mg/mL) coated plates or wells with differentiation medium [DMEM/F12, 10% KOSR, 1% Glutamine, 1% NEAA, 1% Penicillin/Streptomycin, 1% DMSO (Sigma)] containing 100ng/ml of HGF (Peprotech), and 10μM Rock Inhibitor Y-27632. Fresh differentiation medium supplemented with HGF was added daily for 7 days. Matured hepatoblasts cells can be detected by TBX3 (Santa Cruz Biotechnology, sc-166623) and AFP (Santa Cruz Biotechnology, sc8399) staining. To induce immature hepatocyte-like cells to differentiate, differentiation basal medium was supplemented with 10^-7^ M dexamethasone and media was changed everyday for 3 days. Differentiation was confirmed by positive staining for AFP. Final maturation into HLCs was induced in William’s E Media, 10% FBS, 1% Penicillin/Streptomycin, 1.8% DMSO, 1 ug/ml insulin, supplemented with 4.8 ug/ml hydrocortisone. Cells were maintained in this maturation medium for an additional 12 days with media changes every 2-3 days. Mature HLCs were characterized by the expression of HNF4a (Abcam, ab41898) and ALB.

### Single Cell Hi-C Library preparation

#### Plate-based Single cell Hi-C and RNA-seq dual-omic library

##### Cell collection and fixation

Cells were dissociated into single cells and then approximately 1×10^6^ cells were collected to perform double crosslinking with Formaldehyde (FA) and disuccinimidyl glutarate (DSG) as previously described ^1^ with few modifications. Briefly, collected cells were first fixed with 1%FA (Fisher, cat. no. BP531-500) for 10 mins at room temperature, followed by quenching with glycine (128mM final concentration, Sigma, cat. no. G8898-1KG) for 5 mins at room temperature, followed by placing the quenched samples on ice for an additional 15 mins. FA-fixed cells were centrifuged at 1000Xg for 8 mins and resuspended in 990ul DPBS, followed by addition of 10ul DSG (300 mM stock, final concentration 3 mM, ThermoScientific, cat. no. 20593) and 5 ul DRAQ5 dye (to facilitate sorting) for 40 mins at room temperature. The cross-linked reaction was quenched with glycine (400mM final concentration) for 5 mins at room temperature and cells were centrifuged at 2000Xg for 12 mins and then resuspended in 1ml DPBS to be ready for sorting.

##### FACS sorting and lysis

SA suspension of capture beads was prepared prepared by adding 2µl primary antibody anti-Nesprin,10µl Dynabeads™ M-280 Sheep Anti-Rabbit IgG beads into 100ul 0.2% BSA (Millipore, A7906-50G), followed by incubation on a rotator at room temperature for 30 mins. Beads were washed twice with 0.2% BSA in DPBS and resuspended in 40ul DPBS (ThermoFisher, 14190250). 96 well plates with each well containing 5µl of lysis buffer per well (10µM HEPES-KOH (pH 7.5); 2.5 µM MgCl2; 10 µM KCl; 0.1% SDS; 1mM DTT; 2U Rnase inhibitor (Invitrogen, 10777019); 40ul beads suspension) were used to sort a single cell into each well. For asynchronized samples in the protocol benchmarking assay, the gate for DRAQ5 (Thermofish, 62251) was set from G1 to G2 populations, while for the pro-metaphase arrested cell sample the gate was set for the G2 peak. For differentiation samples, DRAQ5 gate setting is the same as the asynchronized sample above. The plates were centrifuged at 700Xg for 30 seconds after sorting. Plates were then incubated at 4°C on a ThermoMixer with constant shaking at 900rpm for 30min followed by briefly spinning at 700Xg for 30 seconds. Plates were placed on the magnet for 3 mins at room temperature. The supernatant (containing cytoplasm) can be transferred to a new 96-well plate for smart-seq3 experiments after de-crosslinking. The remaining magnetic beads with attached nuclei were used to perform single cell Hi-C.

##### Single-nuclei Hi-C

**1.Digestion**: Single nuclei attached to magnetic beads in each well were resuspended in 4 μl/well of 0.3% SDS and incubated at 37°C for 30 min, at 900 rpm; followed by addition 1.1 μl of 10% Triton X-100 (final concentration 1.8%) and incubated on ThermoMixer at 37°C for 30 min, 900 rpm to quench SDS. Then 1.9 ul per well digestion solution is added [0.7 µl 10× rCutSmart; 0.2 μl of 10 U/μl NlaIII (NEB) + 1 µl H2O] is added into each well followed by incubation on a ThermoMixer at 37°C for 1hr, 900 rpm. The restriction enzyme is inactivated by incubation at 65°C for 20 min in a PCR machine. **2. Ligation:** 18ul per well ligation solution [5 µll 5× T4 ligase buffer; 0.25 μl BSA (10mg/mL) + 1 μl T4 DNA ligase (Life Tech. 1U/µl); 2.5 μl 10% Triton-X 100 + 9.25 µl H2O] is added into each well followed by incubation on a ThermoMixer at 16 °C for 1 hr, 900 rpm. **3. Reversal of crosslinks**: 2 µl per well proteinase K (0.5 mg/ml) is added into each well followed by incubation at 65°C for 12-16 hours in a PCR machine. **4. DNA purification:** DNA is purified by using 0.9X AMpure beads by adding 24 µl beads to each well, mixing, and incubating for 10 mins at room temperature. The plate was then placed on a magnetic stand till the beads accumulated at the magnetic side and then the supernatant was discared. The beads were washed with 80% ethanol once. After complete evaporation of the ethanol, 5 µL nuclease free water was added to dissolve the DNA for 5 mins. The liquid containing the dissolved DNA was transferred to wells of a new plate. **5. Tn5 insert adaptor and library amplification:** 5 µL 2XTn5 buffer [40mM Tris-HCl pH7.5; 10mM MgCl2] containing Tn5 enzyme [illumina 20034197, 0.0002 µL\well] was added to each well, followed by incubation at 37°C for 40 mins in a PCR machine. Then, the plate was immediately put on a cold block to slow down the Tn5 reaction. 0.9X AMpure beads were used to purify DNA fragments as described at point 4 above. 9.5 µL PCR buffer [Kapa HiFi HotStart 2×mix; DNA forward primer with final concentration as 0.5 µM (containing i5 index to indicate plate order)] was added directly after all ethanol had evaporated. Then 96 different reverse barcodes were added (one for each well) with final concentration of 0.5 µM (containing i7 index to indicate well position). The plate was placed into a PCR machine the following program was run [72°C, 3 mins; 98°C, 3 mins; {98°C, 15 sec; 67°C, 20 sec; 72°C, 5 mins} 22 cycles; 72°C, 5 mins]. **6**.

###### Library purification

Amplified libraries were collected from each of the 96 wells into 1.7ml LoBinding tubes and DNA was purified using AMpure XP beads twice [0.7X first time; 0.6X second time]. The final libraries were sequenced (PE150) on an illumina HiSeq or Novaseq instrument.

#### Droplet-based Single cell Hi-C library preparation

Droplet-based single-cell Hi-C was performed exactly as described by the published Droplet Hi-C protocol (https://www.protocols.io/view/droplet-hi-c-for-fast-and-scalable-profiling-of-ch-261ge5qwyg47/v1). For each sample, ~6,000 nuclei were targeted for recovery. Following library construction, an additional 0.8× SPRIselect cleanup step was performed to remove DNA fragments shorter than 200 bp. Final libraries were sequenced on an Illumina NextSeq 2000 using a 300-cycle kit, with sequencing configured as Y150, I8, U16, Y150 (Read 1: 150 cycles; Index 1: 8 cycles; Index 2: 16 cycles; Read 2: 150 cycles).

#### Single cell RNA-seq (Smart-seq)

Single cell RNA-seq was performed using the Smart-seq 3 protocol^41^. H1-hESC differentiation samples were dissociated and subjected to FACS. Single cells (singletons) were identified and isolated based on forward and side scatter parameters (FSC/SSC) to discriminate against cell clumps, doublets, and debris before sorting into individual reaction wells. Cells were sorted in 96-well plates and reverse transcription, cDNA amplification, tagmentation and library amplification was performed following the Smart-seq 3 protocol^41^ with little modifications. cDNA was PCR amplified with 27 cycles and tagmented fragments were amplified 20 cycles. Library size selection was performed using two rounds of AMpure bead selection (0.9X + 0.6X). The final libraries were sequenced by using a PE150 protocol on an illumina HiSeq or Novaseq instrument.

#### H1-hESC sorting for different cell cycle stages

H1-hESC cells (1st replicate: 90 million; 2nd replicate: 170 million cells) were crosslinked using the FA+DSG fixation protocol described previously with little modifications^71^. Briefly, collected cells were first fixed with 1% FA (Fisher, cat. no. BP531-500) for 10 mins at room temperature, and then the reaction was quenched with glycine (128 mM final concentration, Sigma, cat. no. G8898-1KG) for 5 mins at room temperature, followed by placing the quenched samples on ice for an additional 15 mins. FA-fixed cells were centrifuged at 1000Xg for 8 mins and resuspended in 990ul DPBS, followed by addition of 10ul DSG (final concentration 3mM, ThermoScientific, cat. no. 20593) for 40 mins at room temperature. DSG-fixed cells were quenched with glycine (400mM final concentration) for 5 mins at room temperature and centrifuged at 2000Xg for 12 mins. Fixed cell pellet was resuspended in 9.3 ml DPBS + 100 µL10% Sanponin (final concentration 0.1%) + 100µL RNase (10mg/ml stock) + 500µL Proteinase Inhibitor (20X stock) for 35 mins in RT (keep in dark) and then centrifuged at 1500xg for 10 mins. 50 million cells were then resuspended in 5 ml DPBS containing 1% BSA with PI (final concentration: 50 µg/ml) stained for 35 mins in dark. Stained samples were submitted to the Umass FACS core facility to perform cell cycle stage sorting. Sorting gates were set as G1, S, G2/M to collect cells at different cell cycle stages. Collected cells were centrifuged at 1000Xg for 15 mins and stored at −80°C.

#### Bulk Hi-C

Double crosslinked cell pellets were thawed and resuspended in Hi-C lysis buffer (Hi-C3.0 protocol^71^). Chromatin was digested with two enzymes: DpnII (NEB R0543M) and DdeI (NEB R0543M) at 37 °C overnight. The overhanging DNA ends were filled in using biotin-14-dATP (LifeTech, 19524016) at 23 °C for 4 hours, and ligated with T4 DNA ligase ^3^ (Life Technologies, 15224090) at 16 °C for 4 hours. Proteinase K (ThermoFisher, 25530031) was added twice, and samples were incubated at 65 °C overnight to remove cross-linked proteins. Ligated DNA was purified using 1X Ampure beads and sonicated to a length of 200-350 bp. End-repaired DNA was captured on streptavidin coated beads (DYNAL™ MyOne™ Dynabeads™ Streptavidin C1, Invitrogen 65001). A-tailing and illumina adaptor ligation were performed with the beads-bound ligation products, and samples were amplified using the NEBNext Multiplex Oligos for Illumina (Dual Index Primers) (NEB, E7600S). Unligated adaptors were removed using 1X AMpure beads and final libraries were sequenced PE50 on a NovaSeq6000 Illumina instrument.

### Imaging

#### Immunofluorescence (IF)

The IF staining process was performed as previously described^51^ with minor modifications. Briefly, cells are seeded at sterile coverslips (ThorLabs, CG15NH) and harvested when cells were ~70% confluent. Cells were washed with DPBS before 4% paraformaldehyde fixation at room temperature for 10 mins. The fixed coverslips were washed 3X with DPBS at room temperature for 5 mins. Cells were then permeabilized with 0.2% Triton-X-100 at room temperature for 10 mins. Coverslips were washed 3X with DPBS at room temperature for 3 mins. 2% BSA was added to block background binding at room temperature for 30 mins, and then primary antibody in the block buffer was added to each coverslip. Coverslips were incubated overnight at 4°C. Coverslips were washed 3x 3 min in 1x PBS, and then a secondary antibody in the block buffer was added and coverslips were incubated for 60 minutes at room temperature. Coverslips were washed 3x 3 min in 1x PBS. Cells were stained with 1 µM DAPI and incubated at room temperature for 10 mins, and then washed 1x 5 min in 1x PBS. Coverslips were mounted with ProLong Diamond mounting media either with DAPI (Invitrogen, P36962) or without DAPI (Invitrogen, P36961) and allowed to cure for at least 24 hours in the dark before imaging. Antibody list: rabbit anti-H3K9me3 (Abcam ab8898, 1:200 dilution), ^4^ anti-H3K27ac (Active Motif 39085, 1:200 dilution), SOX17 (R&D Systems, AF1924), FOXA2 (EMD Millipore, 07-633), TBX3 (Santa Cruz Biotechnology, sc-166623), ALB (Sigma, A6684), AFP (Santa Cruz Biotechnology, sc8399), HNP4a (Abcam, ab41898), SOX2 (Santa Cruz Biotechnology, sc365823), OCT4 (Santa Cruz Biotechnology, sc5279). Secondary antibodies used: goat anti-mouse IgG H+L Alexa Fluor 568 (1:1,000, Abcam ab175473), goat anti-rabbit IgG H&L Alexa Fluo 488 (1:1,000, Abcam ab15007).

Images were acquired in Galvano mode on a Nikon A1 point-scanning confocal microscope equipped with 405 nm, 488 nm, and 561 nm excitation lasers and controlled via Nikon NIS-Elements software (v4.4). Fluorescence was captured through an Apo TIRF 60X oil immersion objective (N.A. 1.49; Nikon) using GaAsP detectors for the 488 nm and 561 nm channels, and a high-sensitivity MultiAlkali PMT detector for the 405 nm channel. All images were collected at a resolution of 1024/1024 pixels.

### Immunofluorescence and DNA FISH co-staining

#### Immunofluorescence staining

Cells were grown on a coverslip till they reached ~70% confluency. Cells were then fixed with 2% paraformaldehyde at room temperature for 10 mins. After fixation, cells were washed 1x with DPBS and then resuspended in 1ml DPBS. Fixed cells were stored at 4°C. Cells were permeabilized with 0.2% Triton-X-100 at room temperature for 10 mins, then washed 3X with DPBS at room temperature for 3 mins. 2% BSA was added at room temperature for 30 mins to block background binding, and then CDT1 (abcam, ab202067) antibody in the block buffer was added to each coverslip, which were incubated overnight at 4°C. Coverslips were washed 3x 3 min with DPBS, and then a secondary antibody in block buffer was added followed by incubation for 60 minutes at room temperature, followed by washing 3x 3 min in DPBS. Cells were fixed again with 2% paraformaldehyde at room temperature for 10 mins and then washed 3x 3 min in 1x PBS. Coverslips can be stored in DPBS at 4°C in the dark.

#### DNA FISH

Coverslips were incubated with 1µg/µL Rnase A in 1XDPBS for 60 mins at 37°C and then rinsed with nuclease free water for 1 min. Coverslip were dehydrated through a gradient of 75%, 85%, and 100% ethanol for 3 mins at room temperature, after which the coverslips were briefly immersed in 100% ethanol. The coverslips were taken out of the 100% ethanol solution and dried (1-2 mins). In case coverslips did not dry in a short amount of time, the 100% ethanol treatment was repeated and dried once more. 15 µL FISH probes were denatured at 70°C for 10 mins [10 pM genomic loci probes (synthesized by Arbor company) and 20 or 50 pM (H1-hESC stages: 50 pM, DE onward stages: 20 pM) whole chromosome painting probe (synthesized by Arbor company) in hybridization buffer (50% formamide; 10% Dextran sulfate; 5X Denhardt’s media)]. Next, the 15 µL probe was spotted on a piece of parafilm spreading on a thin glass and a coverslip with cells treated as above was placed face down on the probe. Coverslips with fixed cells and mounted with probes were denatured by placing the glass carrying coverslip directly on the heat source at 80°C for 10 mins followed by incubation at 37°C in the oven overnight. To prevent evaporation, the chamber was sealed with parafilm during incubation. The next day, the coverslips were placed in a well containing pre-warmed washing buffer (0.5x SSC + 0.1% tween-20) at 37°C for 30 mins, followed by washing coverslips at 42°C with pre-warmed 0.5XSSC + 0.1% Tween 3 × 5 mins on a shaker at room temperature. Coverslips were washed with 2xSSC + 0.1% tween-20; 1xSSC + 0.1% tween-20 for 10 min each time on a shaker at room temperature. Coverslips were rinsed with 1xPBS before DAPI staining. For DAPI stained coverslips: these were incubated for 10min at room temp with 3 µM DAPI in DPBS. Finally, coverslips were mounted with Vectashield (Vector Labs, H-1000-10) and sealed with nail polish. Coverslips can be stored in the dark at 4°C.

### Confocal Imaging

Multiplex imaging was performed on a Leica Stellaris 8 confocal laser scanning microscope (Leica Microsystems) using a HC PL APO CS2 100x/1.40 OIL objective. Excitation was done with the 405 nm laser and a white light laser tuned to 499 nm, 553 nm, 653 nm and 752 nm. Fluorescence emission was collected using detector ranges paired across three sequential acquisition passes to minimize spectral overlap as follows. Setting 1: HyD S 1(415-476nm) and HyD X 4 (665-752nm); Setting 2: HyD X 2(512-553nm) and HyD R 5 (762-829nm); Setting 3: HyD S 3 (570-706nm). The confocal pinhole was set to 1.0 Airy Unit (152.7 um) for each channel. Images were acquired at a resolution of 512 x 512 pixels with a pixel size of 45nm and 3-line averages applied to reduce noise. Optical sections varying in total depth were taken where z-stacks had a z-step size of 0.3 µm. All acquisition parameters were controlled using Leica Application Suite X (LAS X 4.6.1.27508).

### Analysis and quantification

#### Integrated Genome Modeling

Bulk Hi-C data for 5 differentiation stages used to generate pseudo single-cell models were obtained from Huo 2026^44^.

The genome is represented as an ensemble of *s* = 1000 three-dimensional (3D) structures, *x*= {*x*_1_,*x*_2,_ …,*x*_s_}. Each genome structure is defined as a collection of 3D vectors specifying the spatial coordinates of chromatin domain centers, 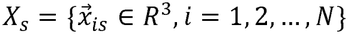. Here, *N* denotes the total number of 200-kb chromatin regions in the diploid genome. For autosomes in the diploid genome, each genomic region is represented by two homologous chromatin domain copies, resulting in a total of ~*N* = 29,838 diploid chromatin domains at 200-kb resolution. Chromosomes were modeled as polymer chains, subject to a combination of physical and data-driven restraints, including chain connectivity, excluded-volume interactions, chromatin contact restraints, and nuclear volume confinement. The nuclear space was modeled as an ellipsoidal confinement defined by three semi-axes, representing experimentally measured average nuclear dimensions. Cell-type–specific nuclear geometries were incorporated into the modeling framework by assigning distinct semi-axis lengths for each cell type. Specifically, the semi-axes (in nanometers) were set to (4368, 3369, 2889) for ES cells, (7570, 5124, 1537) for DE cells, (6309, 4270, 1793) for HB cells, (6822, 4698, 1577) for iHLC cells, and (8056, 5854, 1331) for HLC cells. Each 200-kb domain was modeled as a spherical bead with excluded volume. Bead radii were set separately for each cell type so that the diploid genome occupies approximately 60% of the nuclear volume (PMID: 26951677), i.e. r = (0.6 · abc / N)^(1/3), where a, b and c are the semi-axes of the stage-specific ellipsoidal nucleus and N = 29,838 is the number of beads. This gives a bead radius r = 94.9 nm (ES), 106.2 nm (DE), 99.0 nm (HB) and 108.1 nm (HLC); within each model all beads have the same radius.

Ensemble Hi-C data are represented by a contact probability matrix *A*= (*a*_*l]*_)_*H*x*H*_, where *a*_*l]*_ is the contact probability between the genomic regions *I* and *J*, with *H* as the number of genomic regions when homologous copies are not distinguished. Capital indices (e.g., *I,J*) indicate genomic regions without resolving homologous copies, whereas lowercase indices (e.g., *j, j′* and *j, j′*) explicitly distinguish between the two homologous chromatin domains corresponding to regions *I* and *J*, respectively. The contact probability matrix *A* does not specify which particular pairing of homologous chromatin domains contributes to an observed contact. To account for this missing information at the single-cell level, we introduce a latent contact indicator tensor *W*= (*w*_*ijs*_)_*N*×*N*×*s*_, which is a binary-valued third-order tensor specifying the presence or absence of contacts between chromatin domains *i* and *j* in structure *s*,. Specifically, *w*_*ijs*_ = 1 indicates that chromatin domains *i* and *j* are in contact in structure *s*,, whereas *w*_*ijs*_ = 0 indicates the absence of such a contact.

The objective of the modeling procedure is to infer a population of diploid 3D genome structures that collectively reproduce the ensemble Hi-C contact information encoded in the matrix *A*. This is achieved by jointly estimating the structural ensemble *X* and the associated latent contact variables *W*, such that both are statistically consistent with the observed data. Following the IGM framework, this inference problem is formulated as a maximum-likelihood estimation task.

Model parameters are optimized through an iterative procedure based on a variant of the expectation–maximization (EM) algorithm, combined with scalable optimization strategies previously described^33,34,73^.

Each iteration consists of two alternating steps: an assignment step (A-step) and a modeling step (M-step). During the A-step, the optimal allocation of chromatin contacts in the latent tensor *W* is inferred by maximizing the log-likelihood of all possible contact assignments, while keeping the genome structures *X* fixed to those obtained in the previous iteration. This step determines which homolog-specific domain pairs in a structure can form contacts that are most consistent with the ensemble Hi-C data (see Yildirim et al^32^; for methodological details).

In the subsequent M-step, the genome structures *X* are updated while holding the contact assignments *W* fixed. Physical contact restraints corresponding to the allocated contacts are imposed to generate 3D structures using a combination of molecular dynamics–based simulated annealing^74^ and conjugate-gradient energy minimization (DOI: 10.6028/jres.049.044), as implemented in the IGM modeling framework^33^. This optimization enforces the physical realization of all contacts specified in *W*, while simultaneously satisfying polymer chain connectivity, excluded-volume interactions, and nuclear confinement constraints.

To improve robustness and reduce sensitivity to experimental noise, Hi-C contact restraints are incorporated progressively using a stepwise strategy. Modeling begins with the most confident interactions, ranked by ensemble contact probability *P*(*I, J*). At each optimization stage *t*, only contacts with probabilities *P* ≥. θ _*t*_ (denoted as 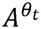)are included. The A/M optimization cycle is then performed iteratively until convergence is achieved for the current threshold. Subsequently, the probability cutoff θ _*t*_ is lowered (e.g., from 1.0 down to 0.002), allowing additional contact restraints to be introduced in successive rounds. Convergence at each stage is assessed using normalized restraint residuals (PMID: 35817938). A restraint is considered violated when its residual exceeds 0.05, and an optimization stage is considered as converged when the global violation rate falls below 0.5%. The procedure is repeated until the final probability threshold is reached and all structures in the ensemble exhibit full convergence. As demonstrated previously, gradually increasing optimization stringency in this manner improves exploration of the solution space, and facilitates the identification of cooperative chromatin interaction patterns^32–34^.

### Calculation of structural features

#### Average radial distance (RAD)

Mean radial distance (RAD) of a chromatin region *I* from the nuclear center is calculated as follows: for each structure *s*,, we compute the normalized radial distance *r*_*is*_ of each locus copy *i* ∈ *I* using the nuclear semi-axes (*a, b, c*) and then average across loci in *I* and across all *S* structures in the population:

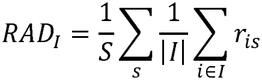

Where *r*_*is*_ is given by

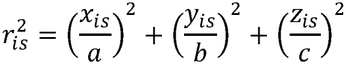

Where (*x*_*is*_, *y*_*is*_, *z*_*is*_) denoting the 3D coordinates of locus *i* in the structure *s*,.

### Average distance to the lamina (LaD)

Average distance to the lamina (LaD) of a chromatin region *I* is calculated as the ensemble averaged, normalized distance of loci *I* from the nuclear lamina.

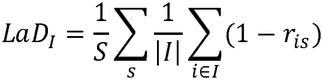

### Average distance to the closest speckle (SpD)

Speckle positions and average distance to the closest speckle (SpD) of a chromatin region *I* are determined following the procedure described in Yildirim et al^32^. SpD is defined as the ensemble-averaged minimal Euclidean distances from for each locus *i* and *i’* to their respective nearest predicted speckle location in each structure.

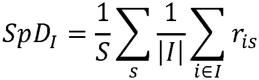

Speckle positions were predicted using the Markov Clustering algorithm (MCL), as described previously^32,33^. Briefly, we used A1-subcompartment regions, which show a strong tendency to localize near nuclear speckles^9^, as an enriched input set for speckle detection. A1 regions were defined at 200-kb resolution as the A1 interaction profile group (IPG), identified by spectral clustering of trans Hi-C contact profiles as described by Spracklin et al^75^. An interaction network was then constructed among A1 regions, and densely connected subgraphs were identified by MCL clustering, requiring a minimum cluster size of five genomic bins. Putative speckle locations were then determined as the geometric centers of these chromatin clusters. Thus, A1 subcompartment identity was used to guide detection of speckle positions but did not itself determine whether individual genomic regions were classified as speckle-associated.

### Cis-to-total contact ratio from 3D models

Each chromatin bead is defined by (*i, a*), where *i* is the genomic locus and *a* ∈ {1,2} denotes the homolog allele. Likewise, a neighboring chromatin bead is (*j, b*) with *b* ∈ {1,2}. For allele *a* in structure *s*,, we define its 3D neighborhood list as

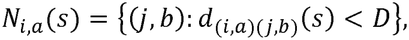

excluding the particle itself (*j, b*)=(*i, a*), and with *d*_(*i,a*)(*j,b*)_ as the center-to-center Euclidean distance between bead particles (*j*, *b*) and (*i, a*) and *D*= 2.5*r* with *r* as the bead radius.

We then divide the neighborhood list of *N*_*i,a*_ (*s*) into cis (intra-chromosomal) and trans (inter-chromosomal) interaction sets:

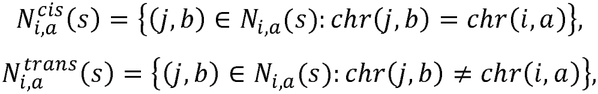

Let 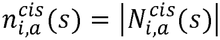 and 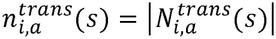 denote the corresponding cis and trans neighbor counts for chromatin region (*i, a*)in structure *s*,.

For each allele *a* ∈ {1,2} of locus *i* in structure *s*, we define the cis-to-total contact ratio as:

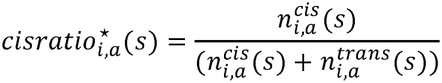

For each genomic locus, the two homologous copies in every diploid structure were treated as independent instances, yielding 2S cis-to-total ratios across S structures. The mean of these values defined the population-average cis ratio of the locus, whereas their standard deviation quantified locus-specific folding variability.

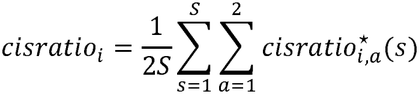

### Median A/B ratio (AB)

For each allele *a* ∈ {1,2} of genomic locus *i* in structure *s*, we partitioned its neighborhood list *N*_*i,a*_(*D* = 4*r* for calculating neighborhood A/B list) into A- and B-associated subsets using the compartment label of each locus (assigned to A or B derived from Hi-C):

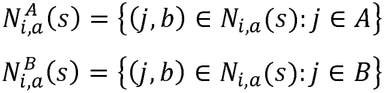

For each allelic locus instance, the corresponding numbers of A- and B-compartment neighbors are

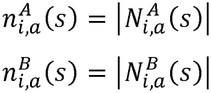

For each allelic locus instance (*i, a*) in each structure *s*,, the A/B ratio is

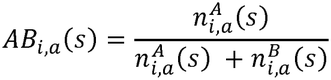

Thus *AB* =1 indicates exclusively A-compartment neighbors and AB=0 exclusively B-compartment neighbors.

Because both homologues allele copies are treated independently, each genomic locus *i* has 2*S* observations across *S* diploid structures.

The median AB fraction per genomic region is then

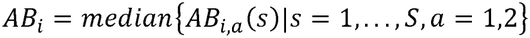

The mean *<u>AB</u>* _*i*_ fraction per genomic locus is defined as:

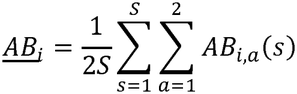

### Chromosome intermingling entropy (Shannon index)

For each locus *i* in structure *s*,, we quantified the diversity of chromosome identities in its 3D neighborhood 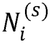 using Shannon entropy. Let 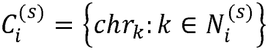 denote the set of chromosome identities represented among the neighbors of locus *i*, where *chr*_*k*_ is the chromosome identity of neighboring locus copy *k*. For each chromosome identity 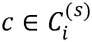 we calculated its relative frequency within the neighborhood as

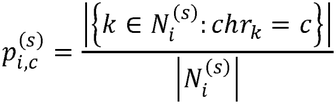

The chromosome-intermingling entropy (i.e. Shannon entropy) of locus *i* in structure *s*, is then defined as

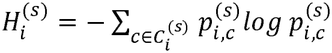

For example, if a locus on chromosome 2 has ten neighbours, six on chromosome 2, three on chromosome 5 and one on chromosome 11, then *C* = {chr2, chr5, chr11}, *p* = (0.6, 0.3, 0.1) and *H* = 0.90.

### Distance to chromosome territory boundary

To define chromosome territories in individual IGM pseudo-single cell structures, we mimicked the procedure of chromosome painting experiments and identified a compact chromosome core based on loci with high cis-to-total contact ratios, which preferentially interact with other regions in the same chromosome. This provides a geometric representation of the densely occupied chromosome territory analogous to that detected by chromosome-painting experiments. For each chromosome copy *m* in structure *s*,, loci were ranked by their cis-to-total contact ratio 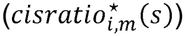. The chromosome territory core of chromosome copy *m* was defined as the set of loci within the highest 30% of this distribution:

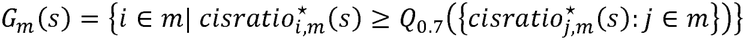

Where *Q*_0.7_ denotes the 70th percentile. The 30% threshold was chosen because it best reproduced the fraction of loci inside their chromosome territory measured by DNA FISH.

The core chromosome territory was then defined as the convex hull of the 3D coordinates of all loci in *G*_*m*_(*s*), with its surface boundary represented as a triangular mesh. d

For each locus *i* in chromosome copy *m* and structure *s*, we calculated the shortest Eucledian distance *d*_*i,m*_(*s*) from its 3D coordinate (*x*_*i,m*_(*s*)) to the convex-hull surface of the core chromosome territory. The distance was assigned a sign according to whether the locus was inside or outside the core chromosome territory: Distances were defined to be negative for loci with coordinates outside the territory convex hull volume and positive for loci with coordinates located inside the core chromosome territory volume.

To make values comparable across chromosome types, single-cell structures and cell types, each signed distance was normalized by the size of the corresponding territory, defined as the distance from the centroid of all core loci coordinates to the chromosome core convex hull surface 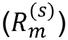 :

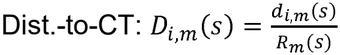

For each genomic locus *i*, CT-based folding variability was quantified as the standard deviation of its normalized signed distance to the chromosome-territory boundary across both homologous copies and all *S* structures, yielding 2*S* observations per locus. Higher values indicate greater cell-to-cell variability in CT embedding and in the extent to which the locus extends beyond the chromosome-territory core.

### Nuclear locale annotation and locale-based metrics

Nuclear locales were annotated across the N = 1,000 pseudo-single-cell 3D genome structures generated by IGM for each differentiation stage. Each 200-kb locus was represented by two homologous copies in every diploid structure, which were treated as independent observations, yielding (2N=2,000 locus instances per genomic region and stage.

For each allelic locus instance in each structure, we calculated its center-to-center Euclidean distance to the nearest predicted nuclear speckle location, the nearest predicted nucleolus location, and the nearest distance to the nuclear envelope. Landmark association was defined using distance thresholds expressed in units of the locus bead radius (*r*). A locus was classified as speckle-associated, and thus occupying the speckle locale, when its distance to the nearest speckle (≤ 5r), nucleolus-associated if its distance to the nearest nucleolus was (≤ 6r).

Envelope association was defined relative to peripheral chromatin distribution within each structure rather than using a fixed absolute distance to the nuclear boundary: The 20% of beads closest to the nuclear envelope were taken as envelope seeds, and every bead that is a seed or lies within 2.5r (centre-to-centre) of a seed was assigned to the envelope locale. This procedure accommodates differences in nuclear size and geometry between structures while extending the envelope locale to chromatin spatially contiguous with the most peripheral regions.

All landmark-distance thresholds were calibrated against independent experimental measurements of subnuclear positioning to maximize agreement between model-derived and experimental localization profiles. Speckle proximity was calibrated against SON-TSA-seq^47^, nucleolar proximity against POLR1 TSA-seq^48^, and envelope proximity against LminaB1 DamID^48^.

Locus instances associated with exactly one landmark were assigned to one of three primary locales: speckle (spe), nucleolus (nuc), or envelope (env). Loci associated with more than one landmark were assigned to the corresponding mixed locale classes (spe-env; spe-nuc; nuc-env). Loci not associated with any landmark were classified according to their local A/B compartment composition of their 3D neighborhood (see section above). For allele (*a*) of locus (*i*) in structure (*s*) the locale A/B ratio was defined as the fraction of A-compartment neighbors among all A- or B-compartment neighbors within a radius of (4r):

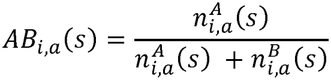

Classification:

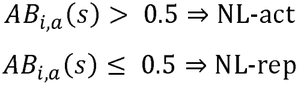

For primary locale analysis, loci simultaneously associated with two or more landmarks were grouped as “others” because of their small sample size.

For each genomic region *i* locale assignments were pooled across both homologues copies and all (*N*) structures for a total of 2,000 observations per locus. The locale occupancy frequency of locale (k) was therefore:

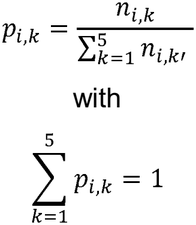

where the five primary, mutually exclusive locales were, env, spe, nuc, NL-act and NL-rep. Instances assigned to “other” were excluded before renormalizing the five primary locale frequencies to sum one.

The resulting five-component locale-occupancy profile (*p*_*i*_ = (*p*_*i,spe*_, *p*_*i,env*_, *p*_*i,nuc*_, *p*_*i,NL*−*act*_, *p*_*i,NL*−*rep*_)) was used to derive three locus-level metrics. First, the “**canonical locale**” (**dominant locale**) was defined as the locale with the highest occupancy frequency:

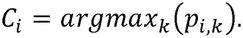

Second, “**canonical strength**” quantified how strongly the most frequently occupied locale was preferred over the second-most frequent locale. If the five locale frequencies are ranked from highest to lowest as

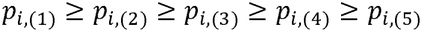

then the canonical strength for locus *i* is

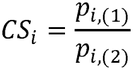

Larger values of *CS*_*i*_ therefore indicate a more pronounced preference for the canonical locale, whereas values approaching 1 indicate similar occupancy of the two most frequent locales.

Finally, the “**locale entropy**” was used to quantify cell-to-cell variability in subnuclear positioning relative to landmarks and functional state:

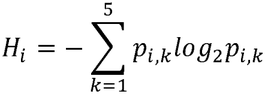

Low locale entropy indicates that a genomic region occupies the same nuclear locale in most cells whereas high entropy indicates that it is distributed across multiple nuclear locales and therefore exhibits greater cell-to-cell variability in subnuclear positioning.

For five possible locales, its theoretical maximum is *H*_*max*_ = *log*_2_(5) = 2.32, indicating even occupancy across the five nuclear locales, while *H*_*min*_ =0 means essentially one fixed locale across cells.

Unless stated otherwise, all locale-based analyses were restricted to autosomal (chr1–chr22) 200-kb regions and excluded centromeric domains according to ES reference annotation.

### Single cell Hi-C analysis

#### Hi-C read mapping and heatmap visualization

##### <Plate-based scHi-C>

Single-cell Hi-C (PE250 FASTQ) sequencing reads were mapped to the human reference genome (Hg38) using Nextflow-based distiller-nf pipeline. The filtering criterion was set to a mapping quality of default-30, followed by removal of duplicate reads. Pseudo bulk Hi-C mapping was performed in the same manner after pooling single-cell Hi-C fastq libraries together. Bulk Hi-C mapped in the same manner, with the filtering criterion set to a mapping quality of default-30.

After mapping, the number of valid pairs in pseudo bulk Hi-C was normalized within each experiment using Cooltools. The data were then binned into .mcool formatted contact matrices at various resolutions, followed by iteratively balancing. Next, .mcool files were loaded in Python-based Jupyter notebooks using the following python packages: cooler (0.9.2), pairtools (1.0.2), pandas (2.3.1), numpy (1.24.4), seaborn (0.12.2), matplotlib (3.10.7), bioframe(0.4.1). The analysis and visualization followed the example scripts provided by cooltools and are available at https://cooltools.readthedocs.io/en/latest/notebooks/viz.html, the python script was available at (https://github.com/dekkerlab/Single_cell_scripts). For single-cell Hi-C, heatmap visualization was performed by loading .mcool files into resgen https://resgen.io.

##### Droplet-based scHi-C

1. **Hi-C Read Processing and Mapping**: Raw sequencing reads from droplet-based Hi-C libraries were processed using a custom pipeline (https://github.com/dekkerlab/Single_cell_scripts). Paired-end reads were aligned to the human reference genome (hg38) with BWA-MEM (v0.7.17), and alignments were converted into BAM format and name-sorted with SAMtools (v1.10). Cell barcodes were extracted from the I2 index read and mapped back to read names. Using a custom Python (pysam v0.16) script, barcodes were assigned as “BX” tags and written into the BAM file.
2. **Pairs File Generation:** BX-tagged BAM files were converted to Hi-C contact pairs using pairtools (v0.3.0). We performed parsing with barcode tags included, followed by sorting and generation of .pairs.gz files.
3. **Merging and Barcode-level Summaries:** Since each 10X index is demultiplexed into four barcodes, we merged all .pairs.gz files into a single dataset for per-barcode summary statistics. Header lines were stripped, and UU (unambiguous–unambiguous) contacts were extracted. For each barcode, we computed the total number of contacts and the number of unique contact pairs, producing a ranked summary of barcodes by interaction count. This step facilitated downstream cell-level quality control and filtering.
4. **Final Deduplication and Sorting:** The merged dataset was further deduplicated based on genomic coordinates and barcodes, followed by coordinate sorting (chrom1, pos1, chrom2, pos2) using GNU sort with multi-threading. The resulting file was compressed into the final .pairs.gz format, which served as the input for downstream analyses, including cis/trans ratio calculation, compartment calling, and variability annotation.

#### Filter interphase cell by Mitosis VS interphase contacts ratio

Single-cell *.pair.gz files were loaded into Python-based Jupyter notebooks, which is available at (https://github.com/dekkerlab/Single_cell_scripts). The contact distance was calculated as df[‘pos1’] – df[‘pos2’], preserving only intra-chromosomal interactions. The ratios of short-range interactions (1.5Kb – 2Mb) and mitotic band (2-12Mb) to total contacts per cell were then calculated. The ranges for Interphase and mitosis cells’ locations in the plot were determined based on these ratios in benchmark assay cells.

#### Single cell compartment strength calculation by referring to bulk annotation

For benchmark, referred compartment profiles were called from pseudo-bulk .mcool files using the scripts in cooltools (https://cooltools.readthedocs.io/en/latest/notebooks/compartments_and_saddles.html) and for differentiation analysis, referred compartment profiles were called from bulk Hi-C .mcool files. Single-cell contacts were filtered based on resolution: contacts with a resolution of < 250,000 bp (for the bin of benchmark pseudo-bulk compartment profiles) or < 100,000 bp (for the bin of differentiation bulk compartment profiles) were excluded. Each contacted locus in single-cell *.pair.gz files was annotated by bulk compartment profiles. The weight of each contact was given from observed counts / expected contacts in geometrical distance. Compartment strength was calculated using the formula:

Compartment Strength = (sum (normalized counts of AA) + sum (normalized counts of BB)) / (sum (normalized counts of AB) + sum (normalized counts of BA)) Permuted single-cell contacts matrix were generated by randomly shuffling the location-ordered single-cell Hi-C contact matrix. The matrix was then re-ordered by sorting the [‘chrom’] column, followed by shifting the [‘pos2’] column by +1 line. Permuted contact matrixes were performed filtering, annotation, and calculation as described above.

#### Density of Interaction distance

Single-cell contacts separation was calculated from single-cell *.pair.gz matrix by [‘pos1’] – [‘pos2’], preserving only contacts that occurred within same chromosome arm and filtering out contacts < 2000 bp. The frequency of single cells contacts separation was then plotted on a density map for each condition.

#### Cell-to-cell variability of genome folding annotation

The single-cell population used to construct cell-to-cell variably folded genome annotation was selected by filtering out cells with fewer than 20,000 contacts. We removed short-range interactions using library-specific distance cutoffs. For plate-based Hi-C, we retained only contacts spanning more than 300 bp across all five stages. When comparing droplet Hi-C with plate-based Hi-C at the ES stage, we applied a >300 bp cutoff to the droplet data as well. When comparing the trend of folding variability across differentiation, we applied a >400 bp cutoff to the droplet Hi-C data from all five stages so that the stages were directly comparable. The cutoffs were chosen based on the average fragment size of each library (**Supplementary Note 2**). The single-cell cis-to-total contact ratio matrix was then generated by counting the cis-contacts to total-contacts per 1Mb in each single cell. The cell-to-cell variability score per bin was calculated using the coefficient of variation across all cells in the matrix. The mean of cis-to-total contact ratio per bin was calculated and compared with the Eigen value of that bin as well.

### Normalization of cis-ratio variability

We modeled an expected standard deviation for each bin, at each of five differentiation stages (ES, DE, HB, iHLC, HLC) profiled by both plate-based and droplet-based single-cell Hi-C, and used the residual between observed and expected standard deviation as a mean-corrected measure of variability. Bins with fewer than 500 (plate-based) or 1,000 (droplet-based) total interaction counts, summed across cells, were excluded prior to fitting to limit the influence of low-depth, high-noise bins.

The functional form used to estimate expected standard deviation was chosen separately for each platform, based on its empirical signal-to-noise characteristics. For droplet-based data, which showed a tight, well-behaved relationship between mean and standard deviation (LOWESS fit R^2^ > 0.9), we used locally weighted scatterplot smoothing (LOWESS; span = 0.4, three robustifying iterations) to empirically estimate expected standard deviation as a smooth, non-parametric function of mean cis-ratio, following standard practice for mean-variance trend estimation in single-cell genomics (e.g., highly-variable-feature detection in scRNA-seq, and DESeq2’s LOESS-based dispersion trend fitting). For plate-based data, which exhibited substantially greater scatter and a less stable empirical relationship, we instead used a theory-grounded model anchored to binomial sampling statistics: expected S.D. = k·√(p(1−p)/n), where p is the bin’s mean cis-ratio, n is its sequencing depth, and k is a single multiplicative overdispersion factor fit by least squares to the platform’s data. This constrained, single-parameter model is less prone to overfitting noisy data than a flexible non-parametric curve, at the cost of assuming the underlying mean-variance relationship follows a scaled binomial shape; k is interpretable as the factor by which true cell-to-cell variability exceeds pure sampling noise.

Both fitting procedures were performed independently for each differentiation stage, rather than pooling stages within a platform, so that each stage’s expected-S.D. curve reflects only that stage’s own mean-variance relationship. For each bin, the residual standard deviation (resid_SD = observed S.D. − expected S.D.) was computed using its own stage’s fitted curve. To account for possible mean-dependence in the spread of the residuals themselves, a locally estimated (quantile-binned) residual standard deviation was computed as a function of mean and used to derive a locally standardized residual (resid_SD_z = resid_SD / local residual S.D.), analogous to a z-score. Fit quality was assessed via the coefficient of determination (R^2^) between observed and expected standard deviation, and residual-versus-mean plots were inspected to confirm the absence of residual mean-dependence after fitting.

### Single cell RNA-seq analysis

#### Mapping

FASTQ reads were trimmed for universal adaptor sequences and the PolyT regions using Trimmomatic-0.36 with the following command “java -jar trimmomatic-0.36.jar PE -phred33 Input_R1.fastq.gz Input_R2.fastq.gz Output_R1.paired.fastq.gz Output_R1.unpaired.fastq.gz Output_R2.paired.fastq.gz Output_R2.unpaired.fastq.gz ILLUMINACLIP:/path/PolyT.fa:2:30:10:8:true”. Then reads were then filtered by length (≥ 150bp) using the following command: “seqkit seq -m 150” before mapping.

Trimmed FASTQ reads were processed using zUMIs version 2.9.7c. Starting from demultiplexing FASTQ files, pseudo barcodes were re-assigned using the scripts as follows: “zUMIs/misc/merge_demultiplexed_fastq.R –dir”. For the *.yaml file setting: reads were filtered for low-quality barcodes (3 bases < phred 20) and UMIs (2 bases < phred 20). UMI were defined from 12 to 19 base in *R1.fastq.gz with detection of pattern “ATGCGCAATG”; cDNA were defined from 23 to 150 base in *R1.fastq.gz; and barcode were defined from 1 to 8 base in *index.fastq.gz. Reads were mapped to the human genome (hg38) using STAR version 2.7.3a with additional STAR parameters ‘--limitSjdbInsertNsj 8000000’. UMI and reads counts were calculated from Ensemble gene annotations “gencode.v29.annotation.gtf”.

### Library quality control and clustering

*dgecounts.rds file generated from zUMIs mapping process, along with barcode file, and gene name file were loaded in R (version 3.6.3) studio. Analysis was performed using Seurat (version 3.2.3) along with other R-based packages: dplyr (version 1.0.10), patchwork (version 3.2.3), ggplot2 (version 3.4.0), biomaRt (version 2.42.1), stringr (version 1.5.0). Library quality control was checked by calculating the percentage of contamination (“percent_mito” and “percent_ribo”) using “PercentageFeathreSet”. Cells with more than 700 transcripts detected were selected, and genes detected in more than 2 cells were preserved. Then data were normalized (‘LogNormalize’), scaled to 10,000 and total number of counts. The top 2000 variable genes were considered and 15 principal components for neighborhood construction (“FindNeighbors”) base on shared nearest neighbor (SNN), cluster classification (“FindClusters”, resolution = 0.5), and UMAP dimensionality reduction (“RunUMAP”).

Batch effect removal was performed using the Harmony package. Merged data from different batches underwent log normalization (‘LogNormalize’), feature selection (top 2000 variable genes), and SCT normalization (‘SCTransform). Batch affection correction was applied using the “RunHarmony” function with “theta = 8 or 10”, followed by the use of five principal components for neighborhood construction (“FindNeighbors”), cluster classification (“FindClusters”, resolution = 0.5), and UMAP dimensionality reduction (“RunUMAP”).

### Expression heterogeneity analysis

To quantify transcriptional heterogeneity while controlling for technical noise, we applied regularized negative binomial normalization using Seurat’s SCTransform function (Hafemeister & Satija, 2019). All genes were retained by setting return.only.var.genes = FALSE, as *“ differentiation_all*.*filt*.*SCT <- SCTransform(differentiation_all*.*filt, return*.*only*.*var*.*genes = FALSE, verbose = FALSE)”*.

SCTransform fits a regularized negative binomial (NB) model for each gene *g* across cells *c*:

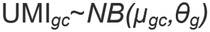

where *μ*_*gc*_ is the expected expression (based on total UMI per cell and gene-specific trend) and *θ*_*g*_ represents the overdispersion parameter of gene *g*.

From this model, Pearson residuals were computed:

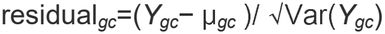

where *Y*_*gc*_ is the observed UMI count, and Var (*Y*_*gc*_) is the variance implied by the fitted NB model. This normalization centers each gene–cell pair relative to its expected value and scales it by its expected variance, placing all values on a comparable scale akin to a z-score. Under a well-calibrated model, residuals driven purely by technical noise should approximate a standard normal distribution (*N* (0,1)), with variance close to 1. Genes with residual variance significantly greater than 1 were thus interpreted as biologically variable.

The resulting Pearson residual matrix—representing normalized and variance-stabilized gene expression values—was extracted from the *scale*.*data* slot of the SCT assay using Seurat’s *GetAssayData*, as*” residual_matrix <-GetAssayData(differentiation_all*.*filt*.*SCT, assay = “SCT”, slot = “scale*.*data”)”*. In this matrix, rows corresponded to genes and columns to individual cells. The matrix was then exported for downstream analysis in Python, where residual variance for each gene was computed directly from the exported residual matrix using: “*residual_variance = residual_matrix*.*var(axis=1)”*.

To quantify gene expression variability across developmental stages, stage labels were assigned to cells based on their identifiers. For each stage, we extracted the corresponding subset of cells and calculated the variance of Pearson residuals across cells for each gene, producing stage-specific measures of expression variability. These gene-level variance scores were subsequently aligned with structural variability annotations (derived from single-cell Hi-C and integrative genome modeling) by matching gene identifiers, enabling integrated analysis of transcriptional and structural dynamics during differentiation. In addition, for the integrated analysis, we filtered out genes with single-cell dropout ratio > 0.85 and bulk RNA-seq (see below) TPM <1.

### Bulk omics data analysis

#### Bulk Hi-C

Bulk Hi-C-seq data for 5-stages were obtained from (Huo 2026^44^) and ES cell phase data were generated in this study. Analyses were performed as described below. Read mapping for bulk Hi-C data was performed the same as described as single-cell Hi-C with minor change, which the reads filtering was set as mapping quality (MapQ) of 30 or higher. MultiQC was used for quality control of the Hi-C library. After mapping, .mcool files generated from binned contact matrices at 1kb, 5kb, 10kb, 25kb, 50kb 100kb, 250kb, 500kb, and 1Mb resolutions were iteratively balanced (ICEed). Then the following Hi-C analysis was performed using python-based Jupyter notebooks using python packages as follows: cooler, cooltools, bioframe, pairtools, pandas, numpy, seaborn and matplotlib. For ES in different cell cycles stages, heatmaps are plotted the region Chr 5: 120,000,000 – 170, 000,000 at 250kb resolution, with Eigenvector 1 plot above each heatmap.

For the compartment analysis in bulk Hi-C, the active and inactive compartment regions were identified using eigenvector decomposition on 250kb binned Hi-C .cool files using cooltools. The positive eigenvalue was given to the more GC rich region of the genome as compartment A (active regions), and the negative eigenvalue was given to more AT rich regions of the genome as compartment B (inactive regions). Saddleplot analysis was followed by the scripts in cooltools (https://cooltools.readthedocs.io/en/latest/notebooks/compartments_and_saddles.html) as described above. The strength of compartmentalization of each distance was calculated as the ratio of (AA+BB)/(AB+BA), where the distance (number labeled in x-axis) represents how many corner bins were used in calculation.

### Bulk RNA-seq

Bulk RNA-seq data were obtained from (Huo 2026^44^), with reanalysis performed here. Paired-end RNA-seq reads were aligned to the reference genome using STAR (v2.X) with a two-pass mapping strategy to improve splice junction detection. Reads were mapped to a genome index with accompanying GTF annotations (genes.gtf), and alignments were generated for both genomic and transcriptomic coordinates (--quantMode TranscriptomeSAM). Key parameters included “--outFilterMultimapNmax 20” and inclusion of SAM attributes for downstream quantification. Unmapped reads were saved for quality control, and software versions (STAR, samtools, gawk) were logged automatically.

Transcript-level quantification was performed using “Salmon” integrated into the mapping pipeline. The resulting gene-level count matrix was imported into DESeq2 (v1.X) in R for normalization and variance stabilization using the “vst” method. The transformed expression matrix (vst) was extracted from the “DESeqDataSet” object and saved as a .tsv file for downstream analyses.

### Gene expression fold change plot and GO analysis

Differential expression between stages was quantified using bulk RNA-seq, and genes were mapped to genomic bins. Genes were stratified into deciles based on residual variance generated from scRNA-seq (above), with the top 20% designated as “high variability.” Volcano plots highlighted these genes, and log2 fold changes (log2FC) were compared between high- and low-variability genes. To control for group-size differences, low-variability genes were randomly downsampled to match high-variability genes.

Gene Ontology enrichment was performed with g:Profiler5. Genes were classified by significance, regulation (up/down), and variability (high/low residual variance). Categories with ≥5 genes were tested against the expressed gene background. Enrichments were filtered to exclude broad terms, and the top 15 GO:BP terms were reported per category.

### Imaging data quantification

#### Compartment score from Immunofluorescence

Confocal images of H3K27ac (euchromatin) and H3K9me3 (heterochromatin) localization were analyzed using Nikon confocal based software: NIS-Elements. Objects corresponding to H3K9me3, H3K27ac and nuclei were identified in each image using the masking function. Briefly, the General Analysis window was opened, and appropriate thresholds were set for individual or combined channels, including single_channel DAPI, combined_channel H3K27ac (red) with DAPI, combined_channel H3K9me3 (green) with DAPI and combined_channel H3K9me3 (green) with H3K27ac (red) with DAPI. The same analysis setting was applied across multiple images to generate a summary table containing information such as “fraction tag”, “coordinates” and “area”. Statistical analysis and plotting were performed in Jupiter Notebook (https://github.com/dekkerlab/Single_cell_scripts).

### DNA FISH

Maximum projection z-stacks from .lif file format to .tif format was performed in Fiji (https://imagej.net/software/fiji/downloads). To enable Fiji to open .lif files, the Bio-Formats Java package was installed from the official website (https://www.openmicroscopy.org/bio-formats/), and the *bio-formats_plugin-8*.*x*.*x*.*jar* file was placed into the *Fiji*.*app/plugins/ folder*. The macro used for batch conversion was downloaded from the GitHub repository *“ImageJ-Export-LIF-as-individual_images*”(https://github.com/pmascalchi/ImageJ_Export-LIF-as-Individual-Images) as the file Export-as-individual-images.ijm. In Fiji, the macro was accessed via Plugins - Macros - Run, where the downloaded file was selected. During execution, the following settings were applied: “Yes” was chosen for batch processing, .lif was entered as the file extension filter, and “Yes” was selected to save maximum projections for Z-stack images. The resulting .tif files were automatically saved in a subfolder within the input file directory.

The maximum projected images were performed using a custom Python script (https://github.com/dekkerlab/Single_cell_scripts) in a Jupyter Notebook to quantify chromatin territories and nuclear loci positioning from maximum-projected DNA FISH confocal images. Nuclear territories were first identified by segmenting the Nucspot 750 (nuclear staining) channel using intensity thresholding and morphological operations to create binary masks. Chromatin territories—representing chromosome territories—were defined based on signal localization in the 488 (chr 11) or 550 (chr 2) DNA FISH channels corresponding to distinct chromosomal probes. Each territory was refined by filtering small objects and applying size and intensity thresholds to remove noise and non-specific signals. For spatial coordinate extraction, the nuclear envelope and chromatin territory boundaries were determined by computing the contours of their respective binary masks using OpenCV. The centroid coordinates of each region were also recorded for subsequent spatial analysis. The contours were used to calculate convex hulls representing the envelope of each region. Individual loci within each chromosome territory were detected based on signal intensity peaks, and their centroid coordinates were recorded.

To quantify spatial relationships, the relative radial distance of each locus from the nuclear center was calculated by normalizing the shortest distance between the locus to the edge of chromosome against the shortest distance from the centroid to the nuclear boundary. To distinguish whether a locus was localized inside or outside its corresponding chromatin territory, a positive or negative sign was assigned firstly. Then the relative position of each locus in chromosome territory was calculated from the distance from the locus to the nearest territory’s boundary dividing the shortest distance from the centroid to the boundary of territory. For the last three developmental stages (HB, iHLC, and HLC), the relative position in territory was manually set to **1** for the radial distance of loci beyond 0.8 of the nuclear radius, which the 0.8 was derived from the mean radial distribution threshold of H3K9me3. Intermingling between two chromosome territories was quantified by calculating the ratio of the overlapping area between their respective binary masks to the total area covered by both territories (i.e., the union of the two masks), providing a measure of spatial overlap.

To quantify the relative volume, radial distance, and sphericity of each chromosome territory, we adopted the established “3D Object Analysis - Cellpose” pipeline available in the Leica-compatible Aivia software (https://www.aivia-software.com). Appropriate thresholds were set for the NucSpot 750 signal to segment the 3D-reconstructed nucleus, and for the 488 nm (chromosome 11) and 550 nm (chromosome 2) DNA FISH channels to identify individual chromosome territories. The same analysis recipe was applied across multiple image stacks to generate a summary table containing measurements such as “Sphericity,” “Volume,” and the “Centroid coordinates” of each detected object (e.g. nucleus and chromosomes). Statistical analysis and plotting were performed in Jupiter Notebook

## Data availability

Any requests regarding cell lines should be directed to Dekker’s laboratory.

## Code availability

Scripts and notebooks used in this study are publicly available in GitHub: https://github.com/open2c and https://github.com/dekkerlab/Single_cell_scripts. IGM software and scripts used for modeling are available in GitHub: https://github.com/alberlab/igm and https://github.com/alberlab/sc-genome-folding-variability-IGM-figures. No other customized codes were developed for this study.

## Acknowledgement

We thank all the members of the Dekker lab and Alber lab for helpful discussions, Y. Li for advice on plate-based technology development, and L. Chang for suggestions on droplet Hi-C. We thank L. Tan, L. Mirny, A. Galitsyna for useful discussions. We thank Core Facilities at the University of Massachusetts Chan Medical School for their assistance: the Deep Sequencing Core, the Flow Cytometry Core, and Sanderson Center for Optical Experimentation (SCOPE) Core. Imaging data was acquired with support from C. Baer and J. McConnell on the Leica STELLARIS 8 STED confocal microscope. Imaging analysis using the Aivia software on the Aivia workstation was funded by a Massachusetts Life Science Center Research Infrastructure grant award to Drs. K. Fitzgerald and C. Baer. The SCOPE RRID is SCR_022721. Diagrams included in figures were created with BioRender.com.

## Funding

This work is supported by the 4DN Center for 3D Structure and Physics of the Genome, funded by NIH UM1 HG011536 (J.D.), NIH UM1 HG011593 (F.A.) and NIH R01 GM162977 (F.A.). J.D. is an investigator of the Howard Hughes Medical Institute.

## Author Contributions

Conceptualization, X.H., J.D., Y.W., and F.A.; single-cell omics and imaging, X.H.; integrative genome modeling, Y.W.; differentiation, X.H., L.Y., J.M., and J.H.; bulk Hi-C experiments, L.Y.; bulk Hi-C analysis, J.G. and X.H.; writing – original draft, X.H., Y.W.; writing – review and editing, X.H., J.D., Y.W., and F.A.; visualization, X.H. and Y.W.; supervision, J.D., F.A. and J.G.; funding acquisition, J.D., F.A., and R.M.

## Declaration of interests

J.D. is on the scientific advisory board of Arima Genomics, San Diego, CA, USA. F.A. is a shareholder of EarlyDiagnostics.

---

**Extended Data Figure 1.**
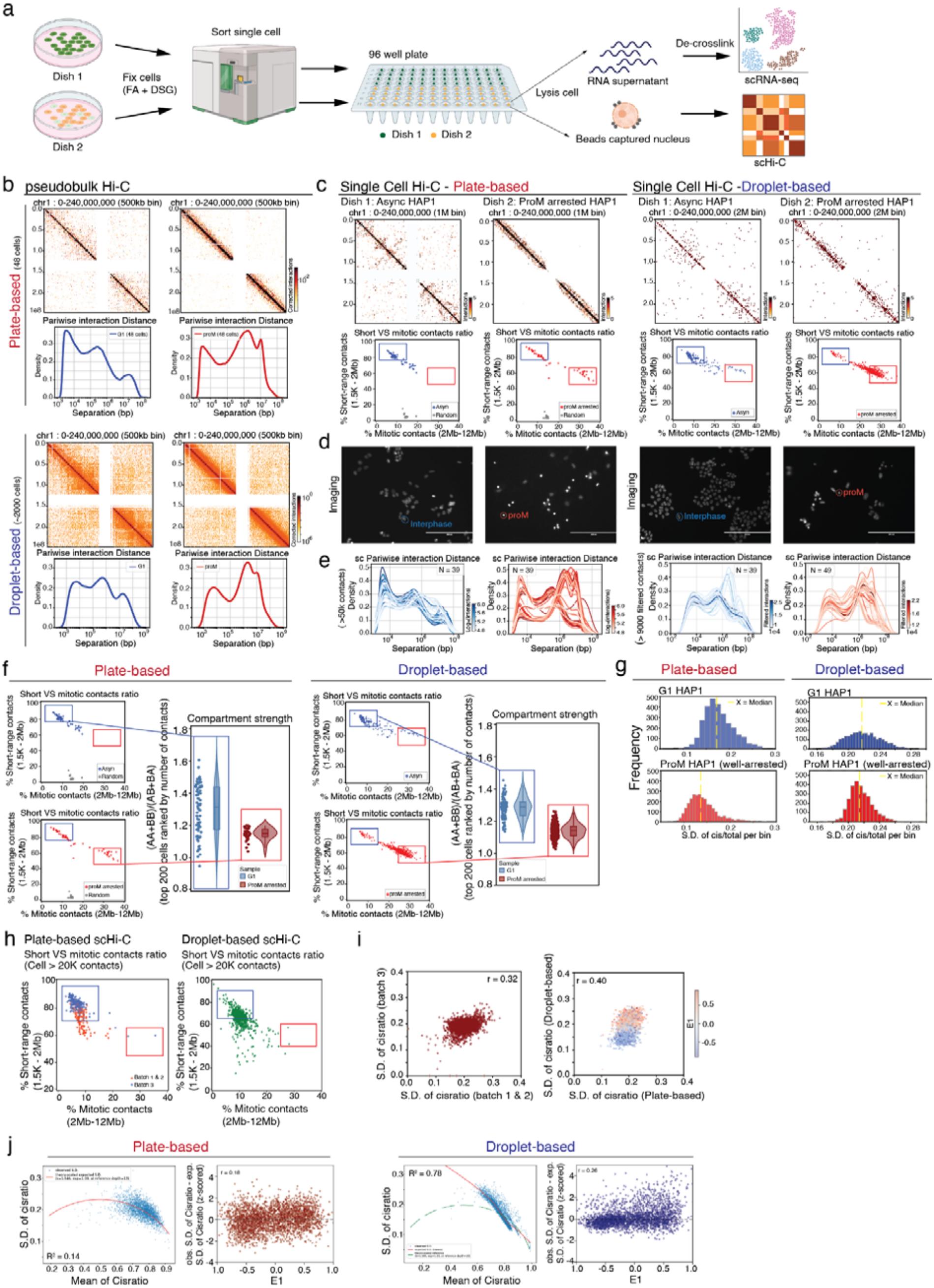
Statistics summary and quality control of single-cell technologies. **(a)** Schematic of the plate-based single-cell co-assay workflow integrating scHi-C and scRNA-seq. Individual cells were fixed (FA + DSG), FACS-sorted into 96-well plates, and processed to separate RNA (for scRNA-seq) and nuclei (for scHi-C) using Nesprin coated beads for nuclei capture during cell lysis; beads captured nuclei processed Hi-C steps in this study; supernatant can be used to get transcripts profile after decrosslink. **(b-e)** Comparison of interphase and prometaphase genome folding states captured by single-cell Hi-C in HAP1 cells. (b) Pseudo-bulk Hi-C contact maps (chr1, 0–240 Mb, 500 kb bins) aggregated from single cells in exponentially grown or prometaphase arrested populations in both plate-based (top) and droplet-based (bottom) workflows. Contacts interaction heatmap and corresponding pairwise interaction distance density plots showing characteristic differences in contact separation profiles between interphase (Dish1 - green) and prometaphase (Dish2 - orange). (c) Top: Representative single-cell Hi-C maps (1 Mb bins). Bottom: Short-range (1.5 kb–2 Mb) vs. mitotic-range (2–12 Mb) contact percentage scatter plots for asynchronous (blue), prometaphase-arrested (red), and simulated random control (grey) cells.(d) DAPI-stained images of asynchronous (blue outline) and prometaphase-arrested (red outline) HAP1 cells. Scale bar: 200 µm. (e) Single-cell pairwise interaction distance density plots for individual cells in each condition. Each curve represents each single cell pairwise interaction distance profile in this group. **(f)** Left: Scatter plot of short-range (1.5 kb–2 Mb) versus mitotic-range (2–12 Mb) contact percentages in exponentially grown population (top) and prometaphase arrested population (bottom), where the blue box indicates G1 population selected and red box indicates prometaphase well-arrested population selected for violin plots on right. Right: Violin plot showing compartment strength scores from plate-based (left) and droplet-based (right) scHi-C benchmark datasets, calculated as (AA+BB)/(AB+BA), where compartment identity was assigned using pseudobulk eigenvector profiles. **(g)** Distribution of structural sc-variability score across genomic bins. Histograms show the standard deviation (S.D.) of cis-to-total ratio across single cells for asynchronous interphase HAP1 cells (top, blue) and prometaphase-arrested cells (bottom, red). The yellow dashed lines indicate the median S.D. in each condition. (Plate scHi-C and droplet Hi-C uses filtered contacts distance > 0.3kb) **(h)** Scatter plots of short-range (1.5 kb–2 Mb) versus mitotic-range (2–12 Mb) contact percentages in hES scHi-C. Pate-based scHi-C workflow (right, two batches) and droplet-based scHi-C workflow (left). **(i)** Scatter plots comparing the S.D. of the cis-to-total contact ratio across genomic bins between plate-based batches (left) and between plate-with droplet-based scHi-C datasets, dots colored by E1 (right). (Plate scHi-C and droplet Hi-C uses filtered contacts distance > 0.3kb) **(j)** Right: Scatter plots of the observed S.D. of the cis-to-total ratio as a function of the mean cis-to-total ratio, overlaid with the expected S.D. fit (red curves).Expected S.D. was modeled using negative binomial theory for plate-based scHi-C and LOWESS (Locally Weighted Scatterplot Smoothing) for droplet-based scHi-C.Left: Scatter plots showing the residual S.D. (observed − expected) as a function of E1 for each 1Mb bin.

**Extended Data Figure 2.**
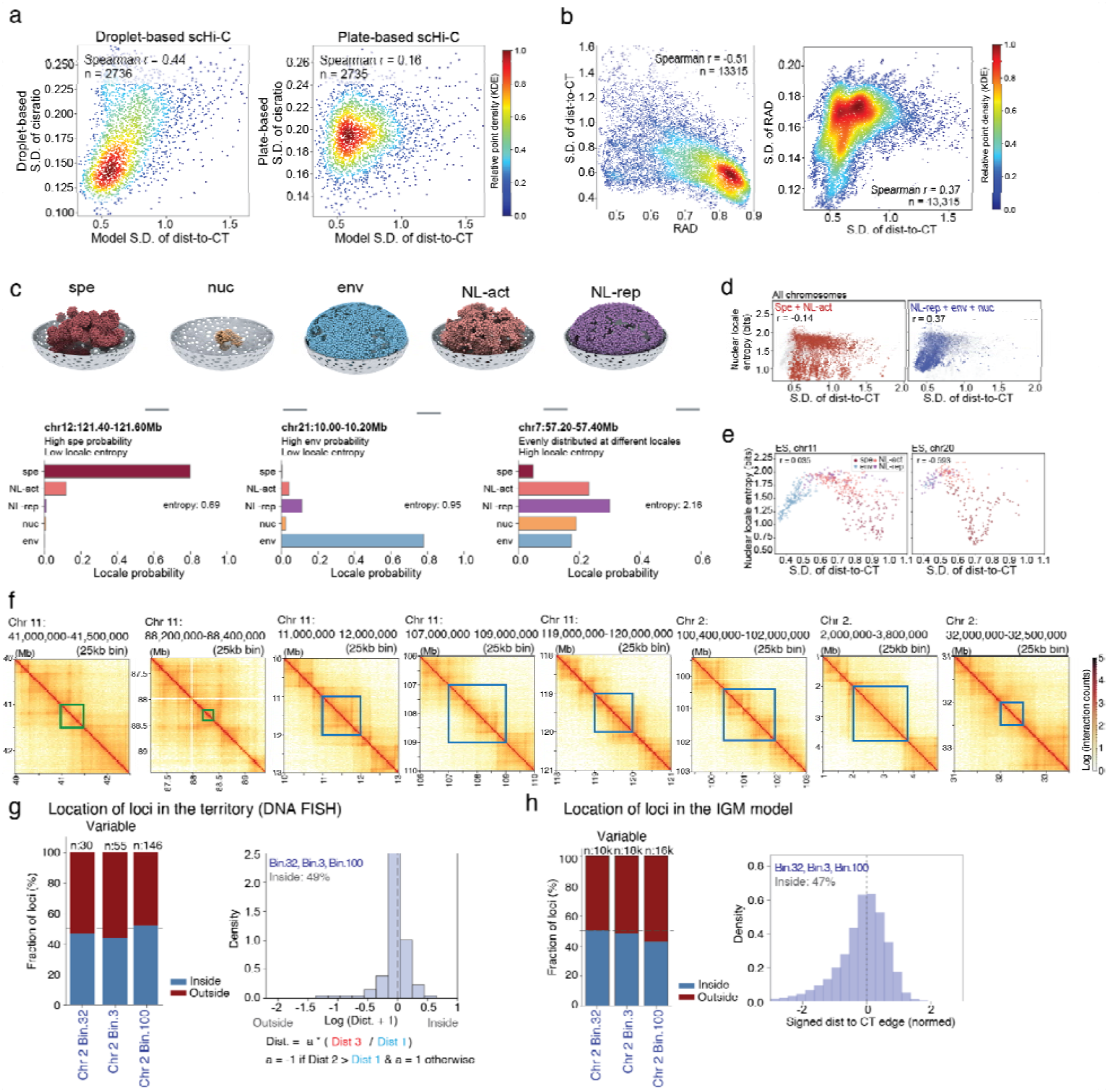
Integrative genome modeling (IGM) and DNA FISH validation. **(a)**Scatter plots comparing the S.D. of cis-to-total across genomic bins from single-cell Hi-C in ES cells with S.D. of dist-to-CT from models. Left: droplet-based scHi-C versus models; right: plate-based scHi-C versus models. Regions are restricted to autosomes and exclude centromeric bins. **(b)**Scatter plot of mean nuclear radial distance versus the S.D. of dist-to-CT (left) and S.D. of dist-to-CT versus S.D. of RAD (right) in the ES-stage from pseudo-single-cell models (200kb bin). Regions are restricted to autosomes and exclude centromeric bins. **(c)**Top: Visualization of the five nuclear locale classes in the ES cells model. Each panel shows one representative 3D structure from the ES cells IGM model, with chromatin beads colored by their assigned nuclear locale. Bottom: Locale probability profiles of three representative loci. For each 200-kb locus, the locale probability is the fraction of structures in the model population in which the locus occupies each locale. Locale entropy is the Shannon entropy of this probability distribution and quantifies how consistently a locus resides in a single locale across the population (low entropy = one dominant locale; high entropy = distributed across many). Three loci illustrate the range: chr12:121.40–121.60 Mb occupies the speckle locale with high probability (high spe probability, low entropy); chr21:5.00–5.20 Mb is predominantly envelope-associated (high env probability, low entropy); and chr7:57.20–57.40 Mb is distributed roughly evenly across multiple locales (high entropy). Entropy values are annotated in each panel. **(d)**Scatter plots of nuclear locale entropy vs. S.D. of dist-to-CT for active locale (red, left) and inactive locale (blue, right). **(e)**Nuclear locale entropy dynamics. Per-bin (200kb) autosomal analyses colored by dominant locale (P argmax). Scatter plots of nuclear locale entropy vs. S.D. of dist-to-CT for chromosome 11 (top) and chromosome 20 (bottom). **(f)**Bulk Hi-C contact maps (1Mb resolution) showing genomic context for the 8 loci selected for DNA FISH validation. Boxes highlight probe regions, where blue indicates relative variably folded regions, green indicates relative consistently folded regions. **(g)**Bar plots showing the relative distance distribution of chromosome 2 loci to CT in the DNA FISH (left). Right: Histograms of Dist. from DNA FISH measurements same as in Fig 2g. Barplots showing the relative distance distribution of chromosome 2 loci to CT in the IGM model (left). **(h)**Histograms of Dist. from IGM structures (right), where Dist. = a × (shortest distance from locus to CT envelope / shortest centroid-to-envelope distance of this CT). a = 1 if the locus is inside the CT; a = −1 if the locus is outside the CT. CT of each chromosome copy per structure is defined as the 3D convex hull of the top 30% cis ratio beads; distances are computed directly on the raw hull. Both homolog copies of each locus are pooled as independent samples across all 1000 structures.

**Extended Data Figure 3.**
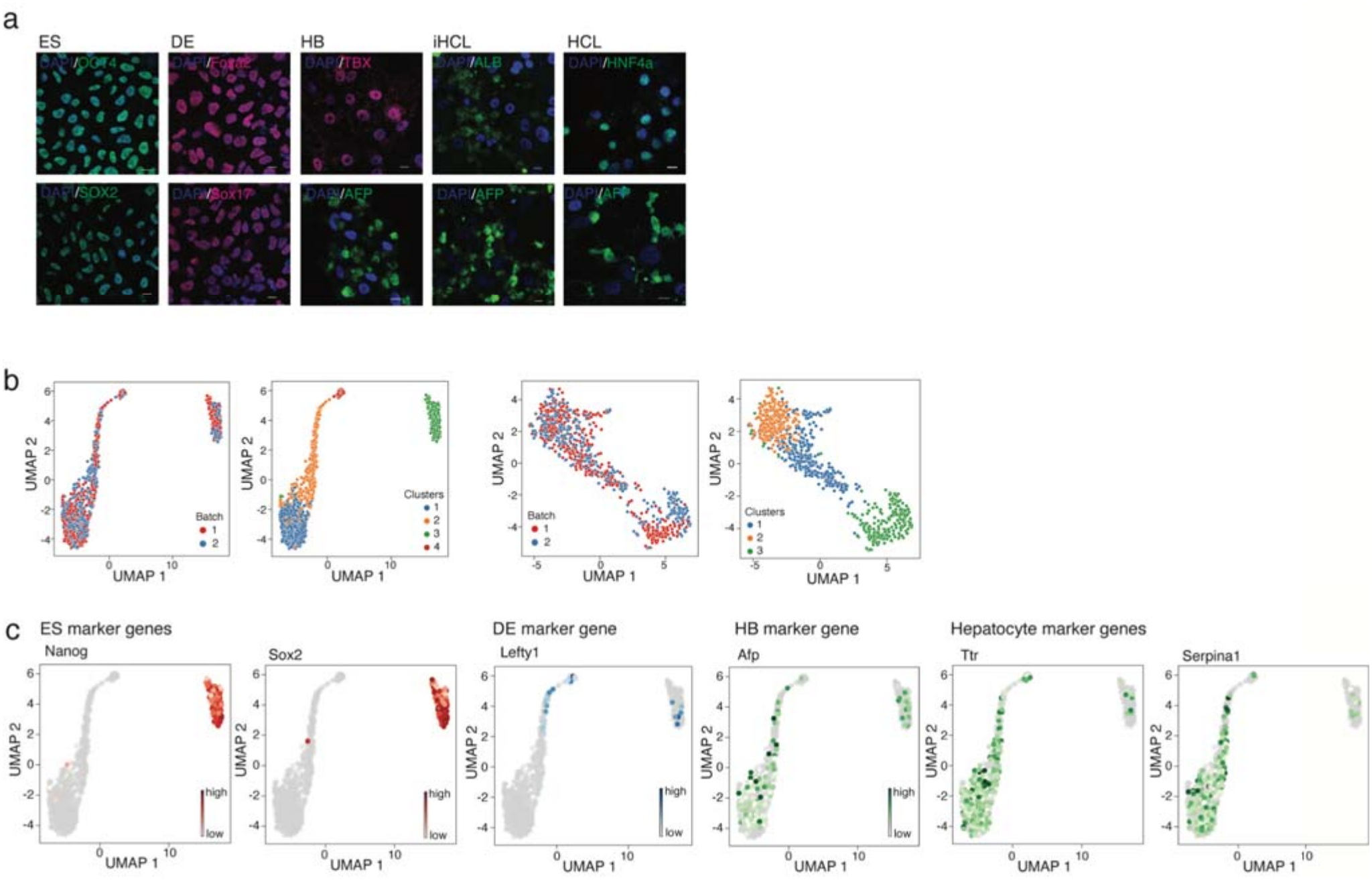
Quality control of differentiation. **(a)** Representative immunofluorescence images showing stage-specific marker gene expression staining across the five stages. Scale bar: 10μm. **(b)** Uniform Manifold Approximation and Projection (UMAP) mapping of single-cell transcriptomes along the differentiation trajectory. Left (first): cells colored by scRNA-seq batch identity across all five stages. Left (second): cell colored by computational defined transcriptional clusters across all five stages. Right (first): UMAP of the four post-ES stages, colored by scRNA-seq batch identity. Right (second): same four stages colored by computational clusters. **(c)** UMAP embedding of single cell along differentiation path, colored by expression of stage-specific marker genes to validate proper assignment of pluripotent and lineage-committed states.

**Extended Data Figure 4.**
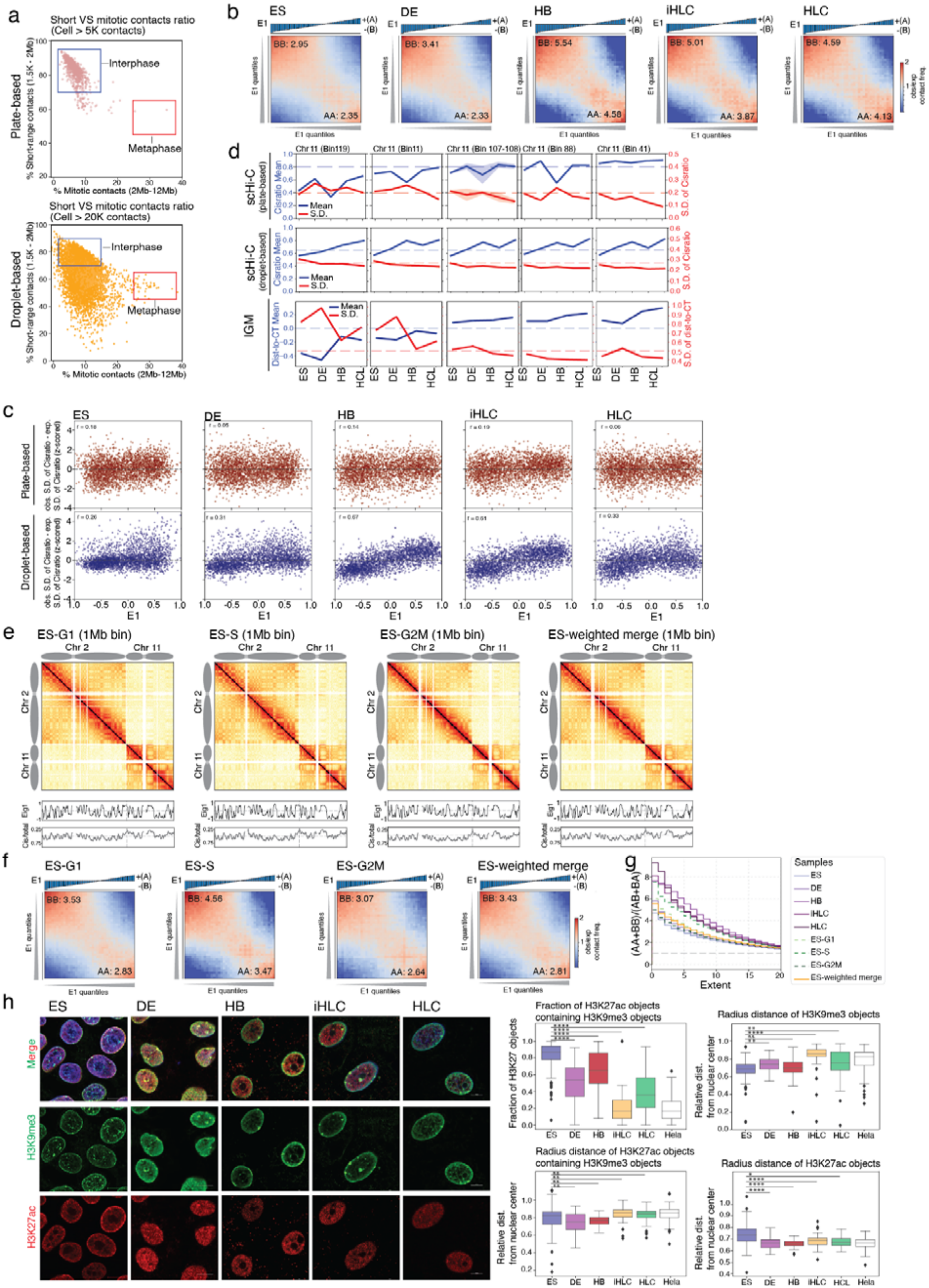
Compartment strength across five stages and H1-hESC cell-cycle phases. **(a)**Scatterplots comparing short-range (1.5 kb–2 Mb) versus mitotic-range (2–12 Mb) contact percentages across individual cells in five stages of differentiation. Top: plate-based scHi-C; bottom: droplet-based scHi-C. Blue boxes indicate G1 phase cells; red boxes indicate metaphase cells. **(b)**Saddle plots from bulk Hi-C showing compartment strength across five stages. The strength of AA compartment and BB compartment are labeled on the right bottom and the left top of the saddle plot (top 20% bins are used to calculate the strength). **(c)**Scatter plots showing the relationship between (observed S.D. − expected S.D.) of the cis-to-total ratio and eigenvector 1 across five stages. Top: Plate-based scHi-C, where expected S.D. is fitted using a negative binomial model based on the mean cis-to-total ratio at the corresponding sequencing depth. Bottom: Droplet-based scHi-C, where expected S.D. is fitted using LOWESS at the local mean cis-to-total ratio at the corresponding sequencing depth. **(d)**Line plots showing the mean (blue) and S.D. (red) of cis-to-total ratio (scHi-C) and dist-to-CT (model) for five chromosome 11 loci examined by DNA FISH. Data used from top to bottom are derived from plate-based scHi-C (top), droplet-based scHi-C (middle) and models (bottom) across five stages. (Plate scHi-C uses batch 1 of 5 stages dataset and filtered contacts distance > 0.3kb; droplet Hi-C uses filtered contacts distance > 0.4kb of 5 stages dataset). **(e)**Top panels (Bulk Hi-C): Contact maps (1 Mb bins) for chromosomes 2 and 11 across four cell cycle stages in ES (G1, S, G2M, pooled by cell phase ratio weighted). Below each map: Compartment eigenvector (Eig1) tracks and cis-to-total contact ratio profiles derived from bulk Hi-C. **(f)**Saddle plots from ES bulk Hi-C showing the interactions strength of different compartments across four cell cycle stages (G1, S, G2M, pooled by cell phase ratio weighted). The strength of AA compartment and BB compartment are labeled on the right bottom and the left top of the saddle plot (top 20% bins are used to calculate the strength). **(g)**Saddle strength profile of compartmentalization from bulk Hi-C across five stages and four cell-cycle stages in ES state. **(h)**Left: representative imaging showing the H3K9me3 and H3K27ac localization across five different stages. Right: statistic quantification of left imaging, including “fraction of H3K27ac objects containing H3K9me3 objects”; “radius distance of H3K27ac objects containing H3K9me3 objects”; “radius distance of H3K9me3 objects”; “radius distance of H3K27ac objects” across five differentiation states and control as Hela cells. Scale bar: 10μm.

**Extended Data Figure 5.**
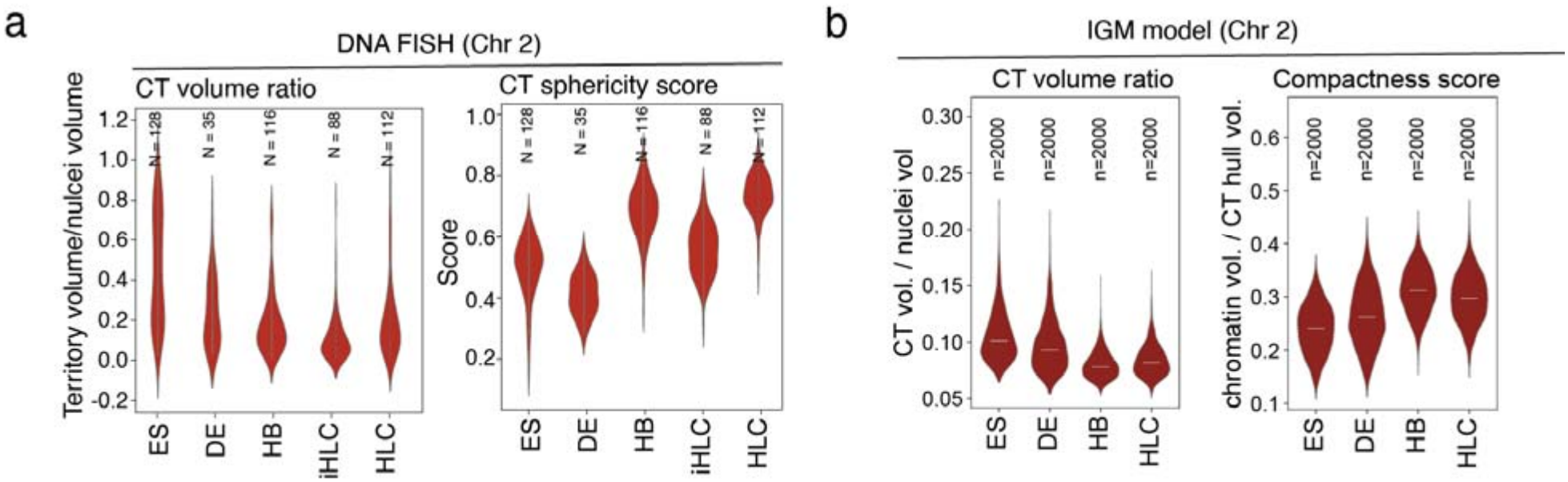
The impact of nuclear structural restraints in nuclear on genome folding variability. **(a-b)** Violin plots show quantified chromosome territory properties for chromosome 2 measured from DNA FISH (a) and from models (b). DNA FISH derived quantification: left: ratio of CT volume to total nuclear volume; right: sphericity scores of the CT in DNA FISH. Model derived quantification: left: ratio of CT volume to total nuclear volume; CT volume here is the volume of convex hull of all the bead from the same copy of chromosome; right: Compactness score (total volume of all chromatin beads in the same copy of chromosome/CT volume).

**Extended Data Figure 6.**
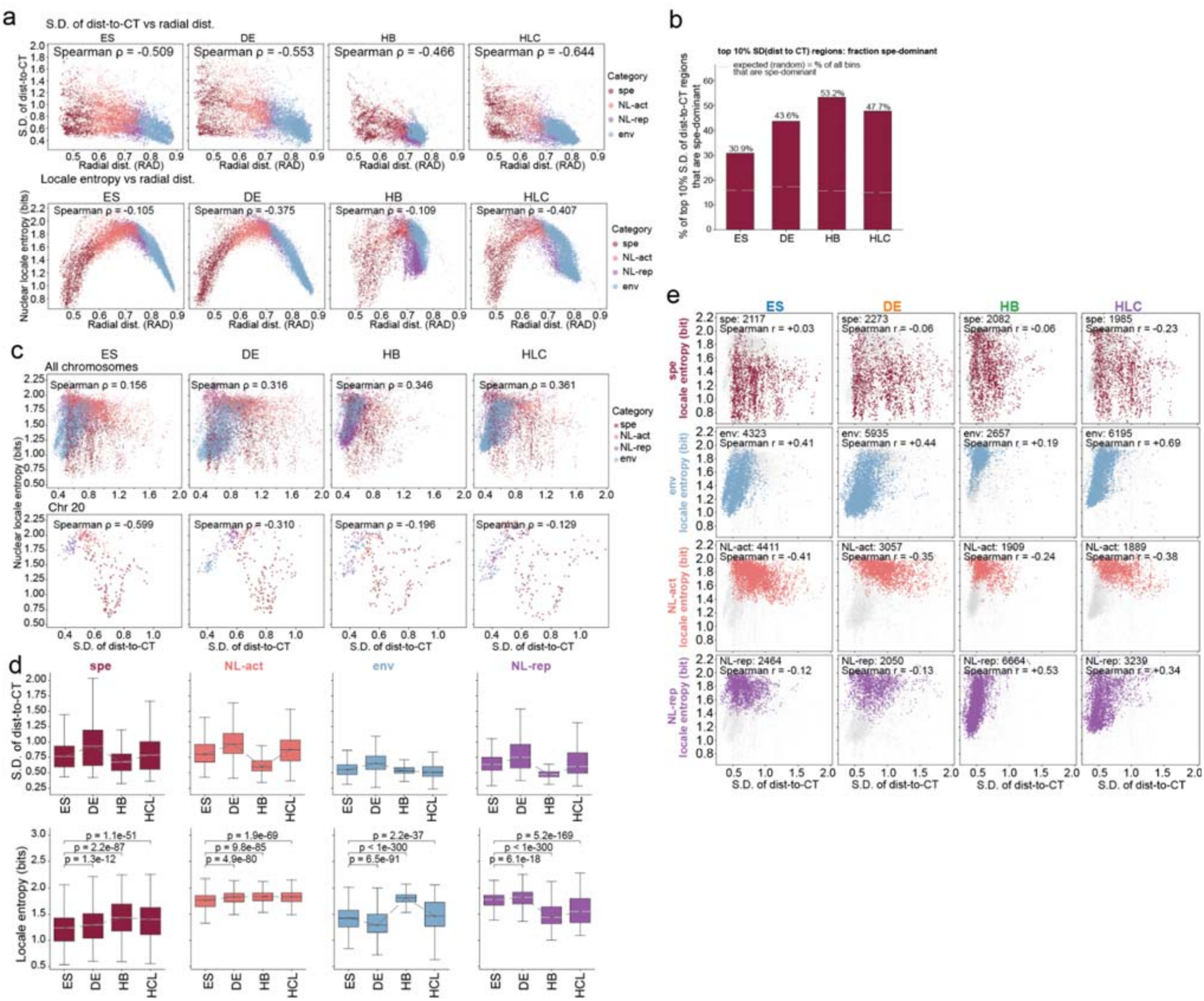
Relocation of genomic regions during differentiation. **(a)**Genome-wide correlation scatter plot of S.D. of dist-to-CT (y-axis) versus normalized radial distance (x-axis) (top) and nuclear locale entropy (y-axis) versus normalized radial distance (x-axis) (bottom)in five differentiation stages. Each dot is one 200-kb genomic region, colored by its dominant locale. Regions are restricted to autosomes and exclude centromeric bins. **(b)**Bar plot indicates the fraction of nuclear speckle locale beads in top10% high S.D. of dist-to-CT genomic regions across five stages. Grey dash indicates the expectation from randomized bins. **(c)**Genome-wide correlation scatter plot of S.D. of dist-to-CT (x-axis) versus nuclear location entropy (y-axis) in each of the five cell lines, colored by its dominant locale. Spearman ρ are calculated between S.D. of dist-to-CT and entropy. Regions were restricted to autosomes and non-centromeric bins. Top: all chromosomes; bottom: chr20. **(d)** Boxplot of S.D. of dist-to-CT (top panel) and nuclear positioning entropy (bottom panel) of regions grouped by dominant nuclear locale across differentiation. Autosomal 200-kb regions (centromeric regions excluded) were assigned a dominant locale in each stage. Boxes are colored by dominant locale; boxes show the median and interquartile range with whiskers, outliers omitted. **(e)** Genome-wide correlation scatter plots of S.D. of dist-to-CT (x-axis) versus nuclear locale entropy (y-axis), separated by dominant locale (rows) and cell line (columns). Each dot is one region: in each panel, regions’ dominant locale. Dominant locale is defined as the argmax over the five normalized association probabilities. Regions were restricted to autosomes and non-centromeric bins.

**Extended Data Figure 7.**
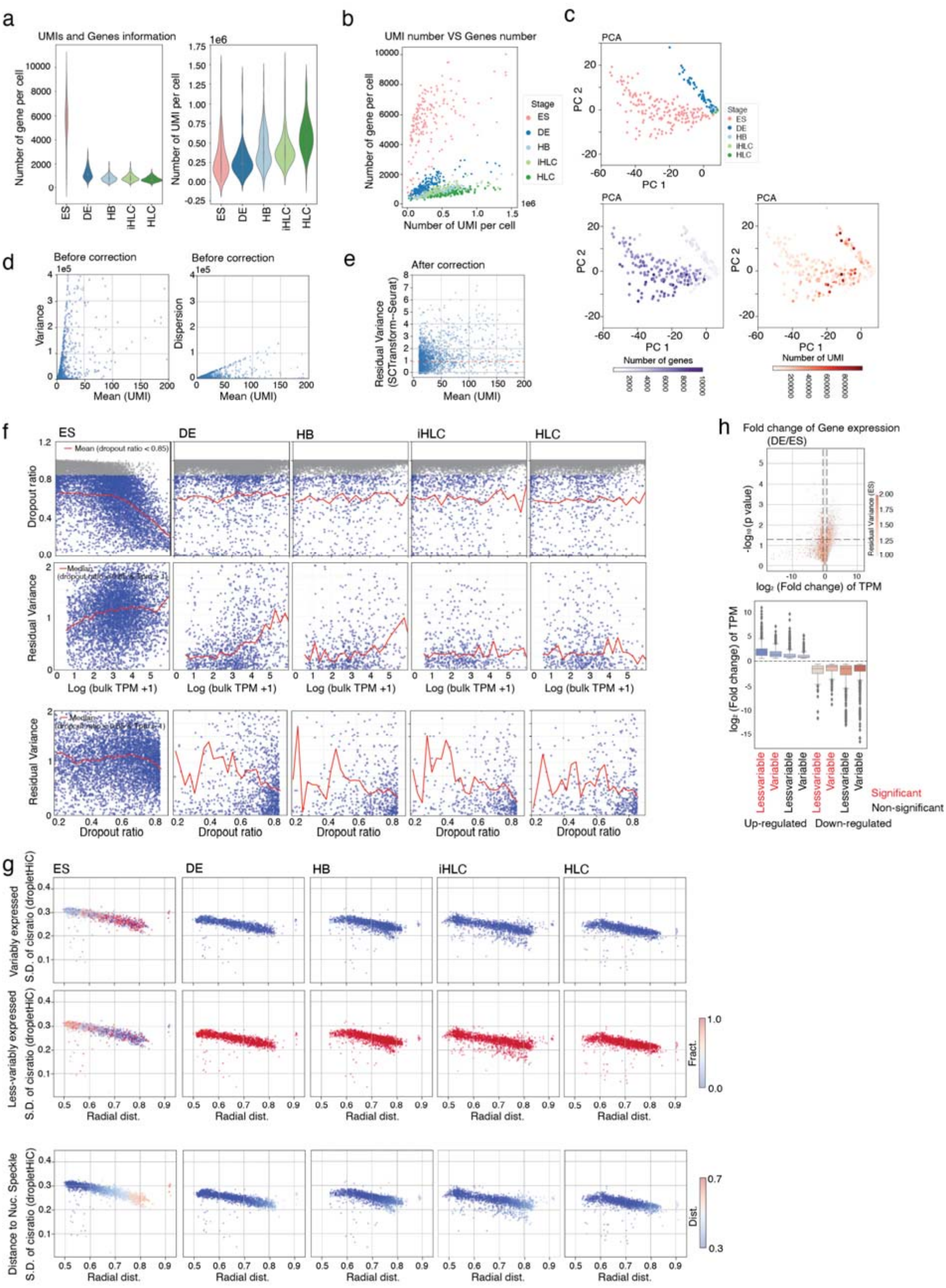
Genome folding variability and gene expression heterogeneity. **(a)** Violin plots showing the number of detected genes (left) and unique molecular identifiers (UMIs; right) per cell from scSmart-seq across five differentiation stages. **(b)** Scatterplots showing the counts of UMI detected per cell with counts of genes detected per cell across five stages. Color marks different cell stages. **(c)**Scatterplots of principal component 1 versus principal component 2 from scRNA-seq. Color marks different cell stages (top), number of genes detected per cells (bottom right) and number of UMI detected per cells (bottom left). **(d-e)** Scatterplots showing metrics of gene expression variability before (d) and after (e) statistical correction. (d) Left: variance of gene across cells versus mean of UMI counts; right: dispersion of gene across cells versus mean of UMI counts. Both use raw values before correction. (e) Residual variance of gene across cells versus mean of UMI counts after fitting with a regularized negative binomial regression (SCTransform), which estimates expected variance based on mean expression and computes Pearson residuals. **(f)** Scatterplots show relationships between gene expression level, expression variability, and dropout patterns across five differentiation stages (ES, DE, HB, iHLC, HLC). Top: Scatter plots showing dropout ratio as a function of log-transformed bulk expression (TPM + 1) for all genes; red lines denote the running mean (genes with dropout ratio >0.85 shown in gray). Middle: Residual variance (cell-to-cell expression variability from scRNA-seq) plotted against log-transformed bulk expression; genes plotted are filtered dropout ratio >0.85 and bulk expression level TPM >1. Red curves indicate running medians, revealing the characteristic dependence of variability on gene expression level. Bottom: Residual variance plotted against dropout ratio; genes plotted are filtered dropout ratio >0.85 and bulk expression level TPM >1, with running medians indicated in red. **(g)** Top: Scatterplots showing relationship between nuclear radial distance and structural variability for less-variable and variable gene groups across differentiation. Genes classified as variably expressed in each stage plotted by their model-derived structural variability (S.D. of cis-to-total contact ratio) versus radial distance of their genomic bins. Middle: Same analysis for less-variably expressed genes. Points are colored by normalized gene expression fraction (Frac.). Bottom: Scatterplots showing relationship between nuclear radial distance, structural variability and nuclear speckle distance of gene loci across differentiation. **(h)** Volcano plot (top) showing differential expression between DE and ES cells, where genes with fold change >2 are colored by their residual variance score in ES. Bottom: boxplot summarizing fold changes across eight categories of genes. Stratified by (significant or non-significant), direction (up-regulated VS down-regulated), and variability (variable vs less-variable)

## References

1. Misteli, T. The Self-Organizing Genome: Principles of Genome Architecture and Function. Cell 183, 28–45 (2020).

2. Dekker, J. & Mirny, L. A. The chromosome folding problem, and how cells solve it. Cell 187, 6424–6450 (2024).

3. Bonev, B. & Cavalli, G. Organization and function of the 3D genome. Nat Rev Genet 17, 661–678 (2016).

4. Bickmore, W. A. & van Steensel, B. Genome architecture: domain organization of interphase chromosomes. Cell 152, 1270–1284 (2013).

5. Falk, M. et al. Heterochromatin drives compartmentalization of inverted and conventional nuclei. Nature 570, 395–399 (2019).

6. Belaghzal, H. et al. Liquid chromatin Hi-C characterizes compartment-dependent chromatin interaction dynamics. Nat Genet 53, 367–378 (2021).

7. Belmont, A. S. Nuclear Compartments: An Incomplete Primer to Nuclear Compartments, Bodies, and Genome Organization Relative to Nuclear Architecture. Cold Spring Harb Perspect Biol 14, a041268 (2022).

8. van Steensel, B. & Belmont, A. S. Lamina-Associated Domains: Links with Chromosome Architecture, Heterochromatin, and Gene Repression. Cell 169, 780–791 (2017).

9. Chen, Y. et al. Mapping 3D genome organization relative to nuclear compartments using TSA-Seq as a cytological ruler. J Cell Biol 217, 4025–4048 (2018).

10. Bhat, P. et al. Genome organization around nuclear speckles drives mRNA splicing efficiency. Nature 629, 1165–1173 (2024).

11. Davidson, I. F. & Peters, J.-M. Genome folding through loop extrusion by SMC complexes. Nat Rev Mol Cell Biol 22, 445–464 (2021).

12. Fudenberg, G., Abdennur, N., Imakaev, M., Goloborodko, A. & Mirny, L. A. Emerging Evidence of Chromosome Folding by Loop Extrusion. Cold Spring Harb Symp Quant Biol 82, 45–55 (2017).

13. Ganji, M. et al. Real-time imaging of DNA loop extrusion by condensin. Science 360, 102–105 (2018).

14. Sabaté, T. et al. Uniform dynamics of cohesin-mediated loop extrusion in living human cells. Nat Genet 57, 3152–3164 (2025).

15. Rao, S. S. P. et al. A three-dimensional map of the human genome at kilobase resolution reveals principles of chromatin looping. Cell 159, 1665–1680 (2014).

16. de Wit, E. et al. CTCF Binding Polarity Determines Chromatin Looping. Mol Cell 60, 676–684 (2015).

17. Vietri Rudan, M. et al. Comparative Hi-C reveals that CTCF underlies evolution of chromosomal domain architecture. Cell Rep 10, 1297–1309 (2015).

18. Gabriele, M. et al. Dynamics of CTCF- and cohesin-mediated chromatin looping revealed by live-cell imaging. Science 376, 496–501 (2022).

19. Valton, A.-L. et al. A cohesin traffic pattern genetically linked to gene regulation. Nat Struct Mol Biol 29, 1239–1251 (2022).

20. Rinzema, N. J. et al. Building regulatory landscapes reveals that an enhancer can recruit cohesin to create contact domains, engage CTCF sites and activate distant genes. Nat Struct Mol Biol 29, 563–574 (2022).

21. Galitsyna, A. et al. Extrusion fountains are hallmarks of chromosome organization emerging upon zygotic genome activation. Nat Commun 17, 2787 (2026).

22. Wike, C. L. et al. Chromatin architecture transitions from zebrafish sperm through early embryogenesis. Genome Res 31, 981–994 (2021).

23. Gibcus, J. H. & Dekker, J. The hierarchy of the 3D genome. Mol Cell 49, 773–782 (2013).

24. Finn, E. H. et al. Extensive Heterogeneity and Intrinsic Variation in Spatial Genome Organization. Cell 176, 1502–1515.e10 (2019).

25. Wang, S. et al. Spatial organization of chromatin domains and compartments in single chromosomes. Science 353, 598–602 (2016).

26. Bintu, B. et al. Super-resolution chromatin tracing reveals domains and cooperative interactions in single cells. Science 362, eaau1783 (2018).

27. Nagano, T. et al. Single-cell Hi-C reveals cell-to-cell variability in chromosome structure. Nature 502, 59–64 (2013).

28. Nagano, T. et al. Cell-cycle dynamics of chromosomal organisation at single-cell resolution. Nature 547, 61–67 (2017).

29. Cao, J. et al. Joint profiling of chromatin accessibility and gene expression in thousands of single cells. Science 361, 1380–1385 (2018).

30. Ramani, V. et al. Massively multiplex single-cell Hi-C. Nat Methods 14, 263–266 (2017).

31. Chang, L. et al. Droplet Hi-C enables scalable, single-cell profiling of chromatin architecture in heterogeneous tissues. Nat Biotechnol 43, 1694–1707 (2025).

32. Yildirim, A. et al. Evaluating the role of the nuclear microenvironment in gene function by population-based modeling. Nat Struct Mol Biol 30, 1193–1206 (2023).

33. Boninsegna, L. et al. Integrative genome modeling platform reveals essentiality of rare contact events in 3D genome organizations. Nat Methods 19, 938–949 (2022).

34. Tjong, H. et al. Population-based 3D genome structure analysis reveals driving forces in spatial genome organization. Proc Natl Acad Sci U S A 113, E1663–1672 (2016).

35. Kalhor, R., Tjong, H., Jayathilaka, N., Alber, F. & Chen, L. Genome architectures revealed by tethered chromosome conformation capture and population-based modeling. Nat Biotechnol 30, 90–98 (2011).

36. Su, J.-H., Zheng, P., Kinrot, S. S., Bintu, B. & Zhuang, X. Genome-Scale Imaging of the 3D Organization and Transcriptional Activity of Chromatin. Cell 182, 1641–1659.e26 (2020).

37. Suter, D. M. et al. Mammalian genes are transcribed with widely different bursting kinetics. Science 332, 472–474 (2011).

38. Chen, K. H., Boettiger, A. N., Moffitt, J. R., Wang, S. & Zhuang, X. RNA imaging. Spatially resolved, highly multiplexed RNA profiling in single cells. Science 348, aaa6090 (2015).

39. Shalek, A. K. et al. Single-cell transcriptomics reveals bimodality in expression and splicing in immune cells. Nature 498, 236–240 (2013).

40. Kolodziejczyk, A. A. et al. Single Cell RNA-Sequencing of Pluripotent States Unlocks Modular Transcriptional Variation. Cell Stem Cell 17, 471–485 (2015).

41. Hagemann-Jensen, M. et al. Single-cell RNA counting at allele and isoform resolution using Smart-seq3. Nat Biotechnol 38, 708–714 (2020).

42. Cremer, T. & Cremer, M. Chromosome territories. Cold Spring Harb Perspect Biol 2, a003889 (2010).

43. Lieberman-Aiden, E. et al. Comprehensive mapping of long range interactions reveals folding principles of the human genome. Science 326, 289–293 (2009).

44. Huo, X. et al. Stepwise reorganization of chromosome conformation and nuclear organization during stem cell differentiation. Preprint at 10.64898/2026.09.16.752023 (2026).

45. Cremer, T. et al. Chromosome territories--a functional nuclear landscape. Curr Opin Cell Biol 18, 307–316 (2006).

46. Croft, J. A. et al. Differences in the localization and morphology of chromosomes in the human nucleus. J Cell Biol 145, 1119–1131 (1999).

47. Zhang, L. et al. TSA-seq reveals a largely conserved genome organization relative to nuclear speckles with small position changes tightly correlated with gene expression changes. Genome Res 31, 251–264 (2021).

48. Kumar, P. et al. Nucleolus and centromere Tyramide Signal Amplification-Seq reveals variable localization of heterochromatin in different cell types. Commun Biol 7, 1135 (2024).

49. Gholamalamdari, O. et al. Major nuclear locales define nuclear genome organization and function beyond A and B compartments. Elife 13, RP99116 (2025).

50. Carpentier, A. et al. Hepatic differentiation of human pluripotent stem cells in miniaturized format suitable for high-throughput screen. Stem Cell Res 16, 640–650 (2016).

51. Hildebrand, E. M. et al. Mitotic chromosomes are self-entangled and disentangle through a topoisomerase-II-dependent two-stage exit from mitosis. Molecular Cell 84, 1422–1441.e14 (2024).

52. Gulati, G. S. et al. Single-cell transcriptional diversity is a hallmark of developmental potential. Science 367, 405–411 (2020).

53. Hafemeister, C. & Satija, R. Normalization and variance stabilization of single-cell RNA-seq data using regularized negative binomial regression. Genome Biol 20, 296 (2019).

54. Meshorer, E. et al. Hyperdynamic plasticity of chromatin proteins in pluripotent embryonic stem cells. Dev Cell 10, 105–116 (2006).

55. Lerner, J. et al. Two-Parameter Mobility Assessments Discriminate Diverse Regulatory Factor Behaviors in Chromatin. Mol Cell 79, 677–688.e6 (2020).

56. Pabba, M. K. et al. Super-Resolution Compatible DNA Labeling Technique Reveals Chromatin Mobility and Organization Changes During Differentiation. Adv Sci (Weinh) 12, e05955 (2025).

57. Dang-Nguyen, T. Q. & Torres-Padilla, M.-E. How cells build totipotency and pluripotency: nuclear, chromatin and transcriptional architecture. Curr Opin Cell Biol 34, 9–15 (2015).

58. Bernstein, B. E. et al. A bivalent chromatin structure marks key developmental genes in embryonic stem cells. Cell 125, 315–326 (2006).

59. Klein, D. C. & Hainer, S. J. Chromatin regulation and dynamics in stem cells. Curr Top Dev Biol 138, 1–71 (2020).

60. Li, F. et al. Bivalent Histone Modifications and Development. Curr Stem Cell Res Ther 13, 83–90 (2018).

61. Harikumar, A. & Meshorer, E. Chromatin remodeling and bivalent histone modifications in embryonic stem cells. EMBO Rep 16, 1609–1619 (2015).

62. Gelali, E. et al. iFISH is a publically available resource enabling versatile DNA FISH to study genome architecture. Nat Commun 10, 1636 (2019).

63. MacArthur, B. D. & Lemischka, I. R. Statistical mechanics of pluripotency. Cell 154, 484–489 (2013).

64. Efroni, S. et al. Global Transcription in Pluripotent Embryonic Stem Cells. Cell Stem Cell 2, 437–447 (2008).

65. Beagan, J. A. et al. Three-dimensional genome restructuring across timescales of activity-induced neuronal gene expression. Nat Neurosci 23, 707–717 (2020).

66. Phillips-Cremins, J. E. et al. Architectural protein subclasses shape 3D organization of genomes during lineage commitment. Cell 153, 1281–1295 (2013).

67. Stadhouders, R. et al. Transcription factors orchestrate dynamic interplay between genome topology and gene regulation during cell reprogramming. Nat Genet 50, 238–249 (2018).

68. Meshorer, E. & Misteli, T. Chromatin in pluripotent embryonic stem cells and differentiation. Nat Rev Mol Cell Biol 7, 540–546 (2006).

69. Kind, J. et al. Single-cell dynamics of genome-nuclear lamina interactions. Cell 153, 178–192 (2013).

70. de Wit, E. et al. The pluripotent genome in three dimensions is shaped around pluripotency factors. Nature 501, 227–231 (2013).

71. Akgol Oksuz, B. et al. Systematic evaluation of chromosome conformation capture assays. Nat Methods 18, 1046–1055 (2021).

72. Scott, S. J., Suvarna, K. S. & D’Avino, P. P. Synchronization of human retinal pigment epithelial-1 cells in mitosis. J Cell Sci 133, jcs247940 (2020).

73. Li, Q. et al. The three-dimensional genome organization of Drosophila melanogaster through data integration. Genome Biol 18, 145 (2017).

74. Kirkpatrick, S., Gelatt, C. D. & Vecchi, M. P. Optimization by simulated annealing. Science 220, 671–680 (1983).

75. Spracklin, G. et al. Diverse silent chromatin states modulate genome compartmentalization and loop extrusion barriers. Nat Struct Mol Biol 30, 38–51 (2023).

