## Supplemental Materials, Notes, and Figures for "Pluripotent and differentiated cells display distinct genomic and nuclear landscapes of genome folding variability"

**Table list:**

| Table 1: | Nuclear dimensions measurement |
| --- | --- |
| Table 2: | Cis-to-total contacts ratio annotation in plate-based single-cell Hi-C |
| Table 3: | Cis-to-total contacts ratio annotation in droplet-based single-cell Hi-C |
| Table 4: | Dist.-to-CT in model |
| Table 5: | Locale entropy in model |
| Table 6: | scRNA expressional residual variance across cells |

**Supplemental note 1**

**Single cell technologies validation.**

To validate these two single-cell Hi-C procedures, we designed a benchmark experiment to test whether the assays could distinguish in a population of HAP1 cells which cells are in mitosis and which are in interphase. To this end, we applied the two assays to two cultures of HAP1 cells: an exponentially growing culture (labeled as dish 1 in **Extended Data Fig. 1a-b**), and a synchronized culture arrested in pro-metaphase (proM) by addition of nocodazole (labeled as dish 2 in **Extended Data Fig. 1a-b**). We generated pseudo-bulk Hi-C results by pooling contacts from all single cells in dish 1 (n = 48 for plate based scHi-C, and n = 3,539 for droplet Hi-C) and for all cells in dish 2 (n = 48 for plate based scHi-C, and n = 3,093 for droplet Hi-C). The results showed that the pseudo-bulk Hi-C map for exponentially growing HAP1 cells exhibits typical interphase genome folding features: a narrow diagonal, a (faint) checkerboard pattern representing compartments, and detectable interactions even for pairs of loci separated by very large genomic distances (10-100 Mb apart). The pseudobulk-Hi-C map for the prometaphase arrested culture shows typical mitotic features: a broader diagonal (for loci separated by up to ~10 Mb) of enriched interactions, and a strong reduction in very long-range interactions. The density distributions of interactions for the two datasets illustrate this clearly (plots below the left two Hi-C interaction maps) (**Extended Data Fig. 1b**).

The difference in the relative abundance of short-range interactions (1.5 kb-2 Mb) vs. longer-range interactions (2-12 Mb) for interphase vs. mitotic cells has been described before and has been used to determine cell cycle stage based on Hi-C maps for single cells ^1,2^. For single-cell analysis we used only cells for which we obtained >20,000 contacts. In addition, for droplet Hi-C datasets we also removed contacts between sites separated by < 400 bp (Methods). We then plotted for each single cell the percentage of short-range interactions vs. the percentage of longer-range interactions (**Extended Data Fig. 1c bottom**). Interphase cells appear on the upper left corner, whereas mitotic cells appear in the lower right corner ^1,2^. In dish 1, containing exponentially growing cells, we did not detect any mitotic cells for the plate based assay, and only a small number for the droplet Hi-C assay. In dish 2, we identified two populations of cells (blue and red boxes). In plate-based assay, around 60% of cells are considered to be in prometaphase, whereas in droplet Hi-C assay, prometaphase ratio is >80%, based on the enrichment of longer-range interactions. This number is similar to the number of mitotic cells in the culture based on DAPI staining of chromosomes (**Extended Data Fig. 1d**).

Another way to identify cell cycle stages for each single cell is by directly plotting the distribution of all interactions as a function of genomic separation. For cells from dish 1, all density distributions display the same shape characteristic of interphase: enrichments of short-range interactions (< 10 kb), enrichment of interactions at the length scale of cohesin-mediated loops (1-200 kb), and very long-range interactions (10-100 Mb). The close alignment of plots illustrates the homogeneity of cell-cycle states in dish 1. In contrast, for cells in dish 2 we detect two distinct shapes of the interaction density plots: one resembling the interphase-like profile, and the second group enriched for interactions at mitotic interaction scales (2-12 Mb) (**Extended Data Fig. 1e**). We conclude that the single Hi-C data obtained with both assays readily distinguishes interphase and mitotic cells.

Single cell Hi-C data also captures compartmentalization, which is strong in interphase cells and absent in mitotic cells ^3^.This is illustrated in **Extended Data Fig. 1f** for single cell data obtained with the plate based assay and droplet Hi-C analysis. We used pseudobulk Hi-C data to determine the pattern of A and B compartments at 250 kb resolution (using eigenvector decomposition). We then quantified compartmentalization strength in single cells in the following way: We determined the number of interactions between pairs of loci (500 kb bins) in the A compartment (A-A interactions), the number of interactions between pairs of loci in the B compartment (B-B interactions), and finally the number of A-B and B-A interactions. We then calculated compartmentalization strength as the ratio of the total number of homotypic interactions (A-A +B-B) divided by the total number of heterotypic interactions (A-B + B-A), for each cell. We selected G1 phase cells from the dish with exponentially growing cells and mitosis of well arrested cells from the synchronized growing cell based on the short VS mitotic contacts ratio, to quantify their compartmentalization strength in single cells respectively. The results showed that G1 phase population shows relatively high levels of homotypic interactions and its compartmentalization strength ratio almost larger than 1.2, while in the group representing mitotic cells, the levels of homotypic and heterotypic interaction are similar, as the compartmentalization strength ratio are 1 to 1.2. From these analyses we conclude that single cell Hi-C data accurately detect presence and absence of compartmentalization observed in interphase and mitotic cells respectively (**Extended Data Fig. 1f**).

**Supplemental Note 2:**

**Distance-based filtering of short-range contacts.**

Our libraries were sequenced on a next-generation sequencer with 2 × 150 bp paired-end reads. Un-ligated DNA fragments longer than 300 bp yield two non-overlapping reads that map close together, and these pairs are incorrectly called as contacts. Because neither the droplet-based nor the plate-based scHi-C protocol includes the biotin fill-in and pull-down steps, such artifacts are not depleted and can account for a large fraction of the called contacts. In Hi-C heatmaps, these false-positive contacts appear as very short-range interactions.

To remove them, we filtered out short-range interactions. This may discard some true contacts, but it substantially reduces false positives. The distance cutoff was set according to the fragment-size distribution of each library. This matters most for droplet scHi-C, in which the DNA is fragmented into very short pieces (~200–400 bp, **Supplemental Fig. 1**).

For the droplet-based libraries across differentiation, we retained only contacts spanning more than 0.4 kb, because one of the droplet libraries had a mean fragment size of ~340 bp. We applied the same cutoff to all stages so that they could be compared directly. For the plate-based libraries, we used a cutoff of 0.3 kb (contact distance > 0.3 kb) to retain as many contacts per cell as possible across all libraries. When comparing the ES stage between the two workflows, we also applied the 0.3 kb cutoff to the droplet-based scHi-C data.

**Supplemental Figure 1:** Single-cell libraries fragmental size detected from bioanalyser

**Droplet Hi-C - hESC stages**

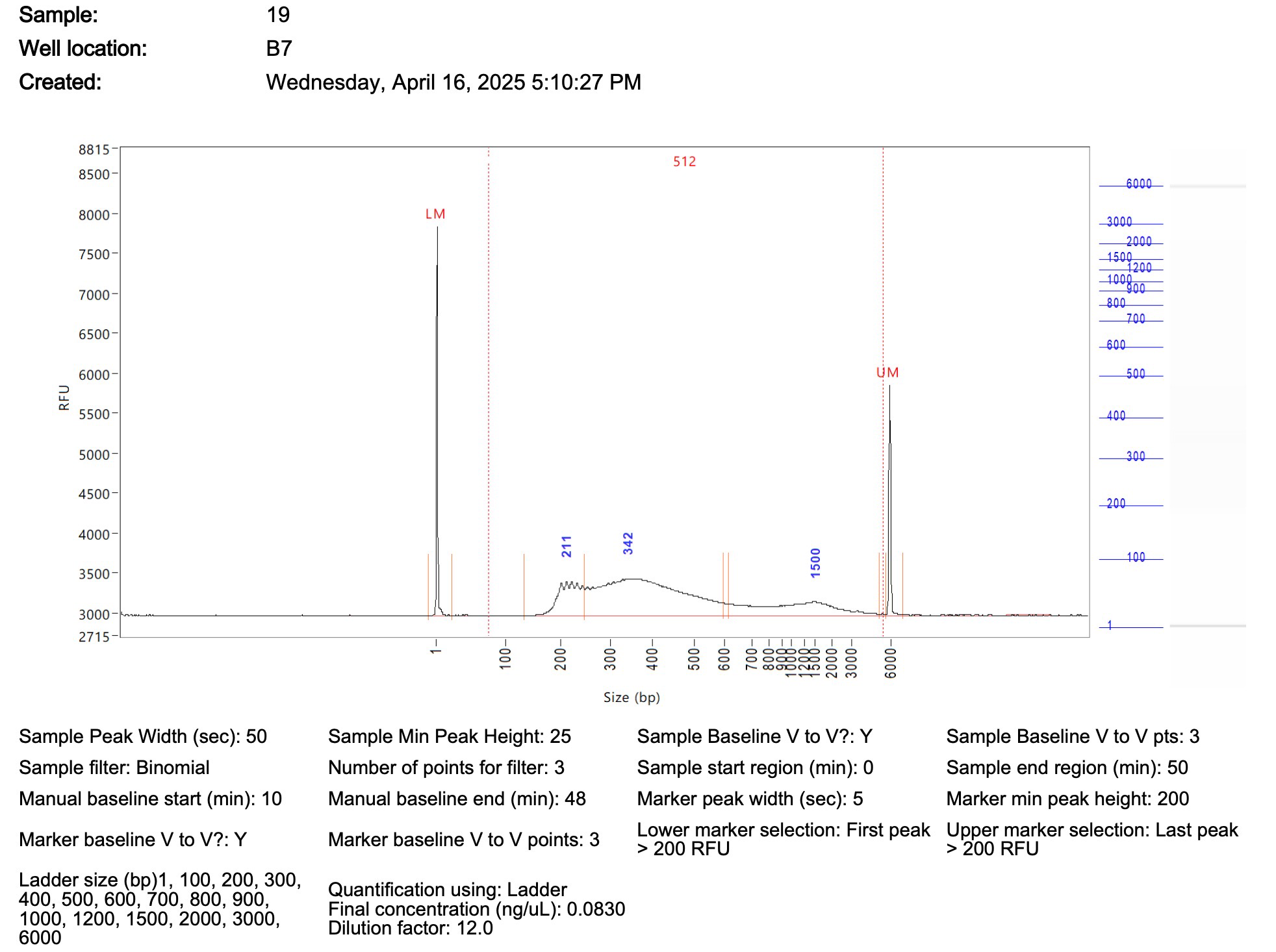

**Droplet Hi-C - DE stages**

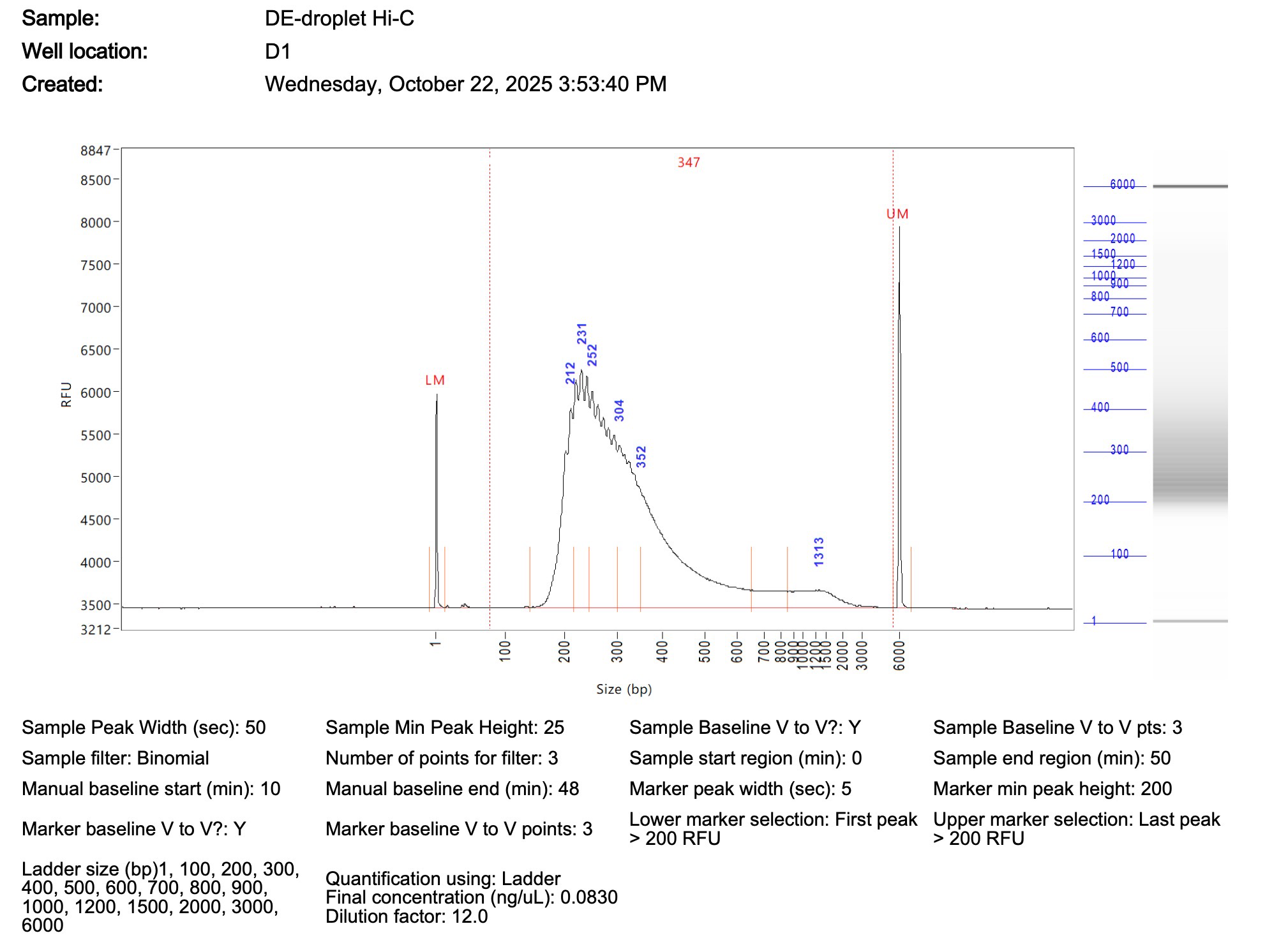

**Droplet Hi-C - HB stages**

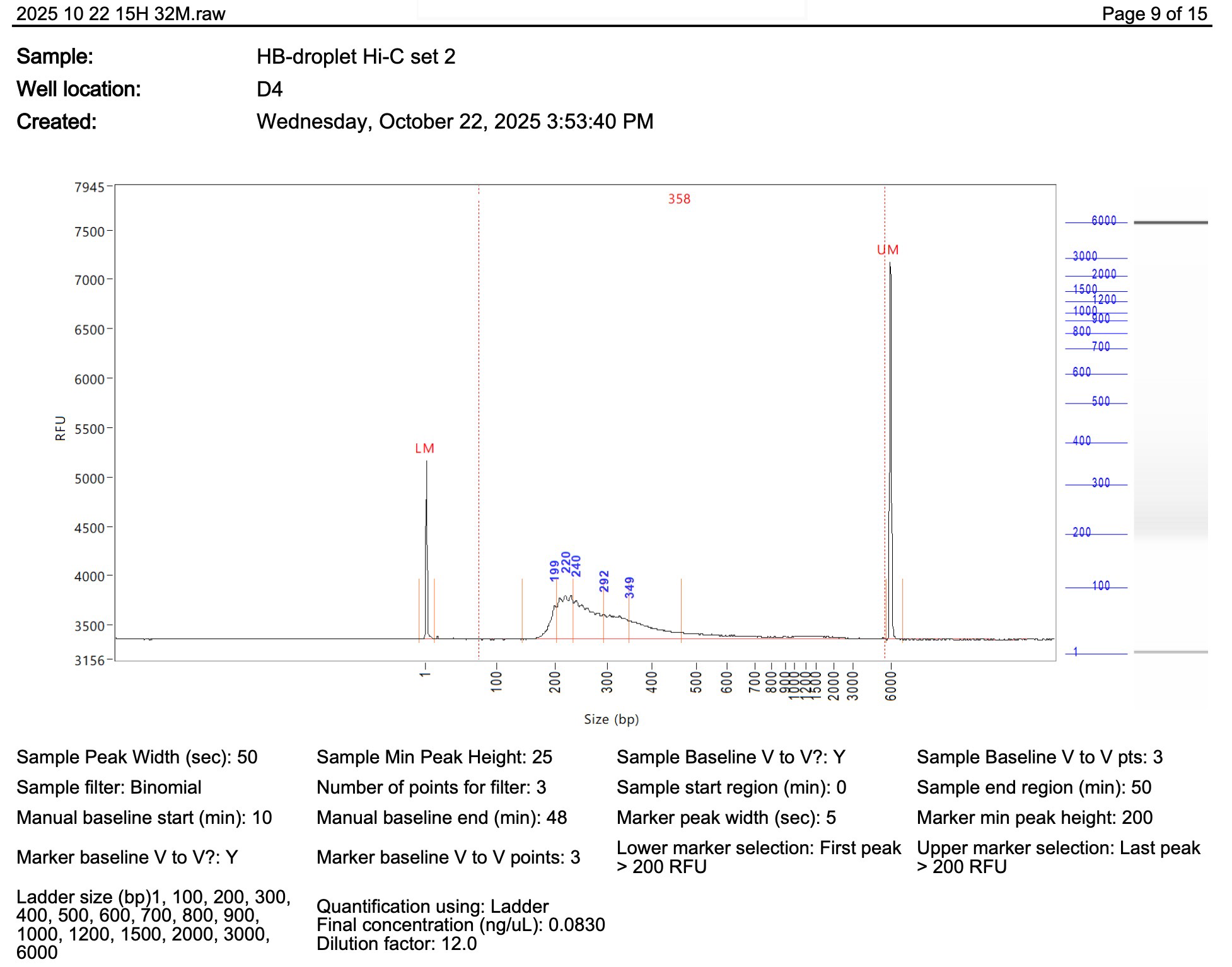

**Droplet Hi-C - iHLC stages**

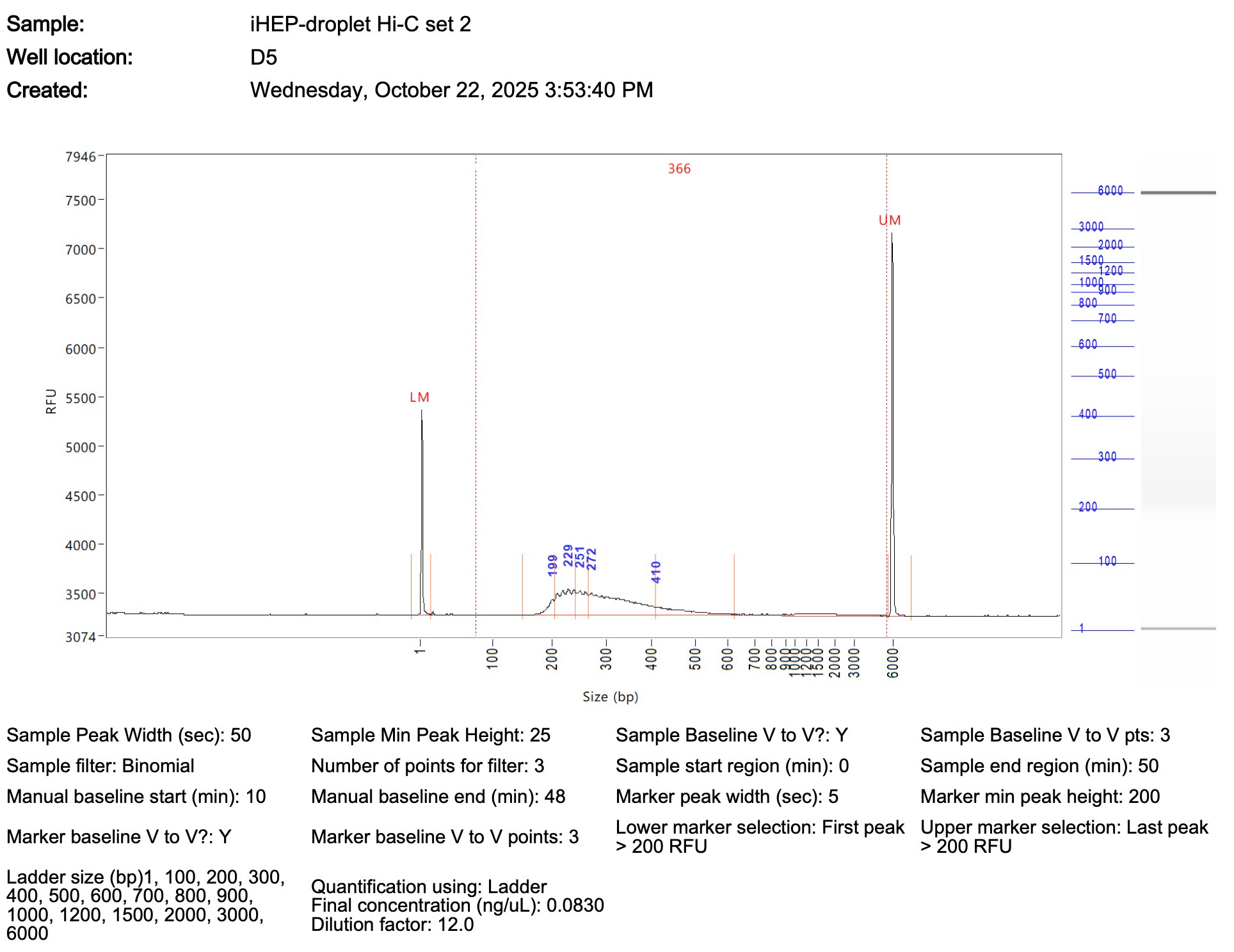

**Droplet Hi-C - HLC stages**

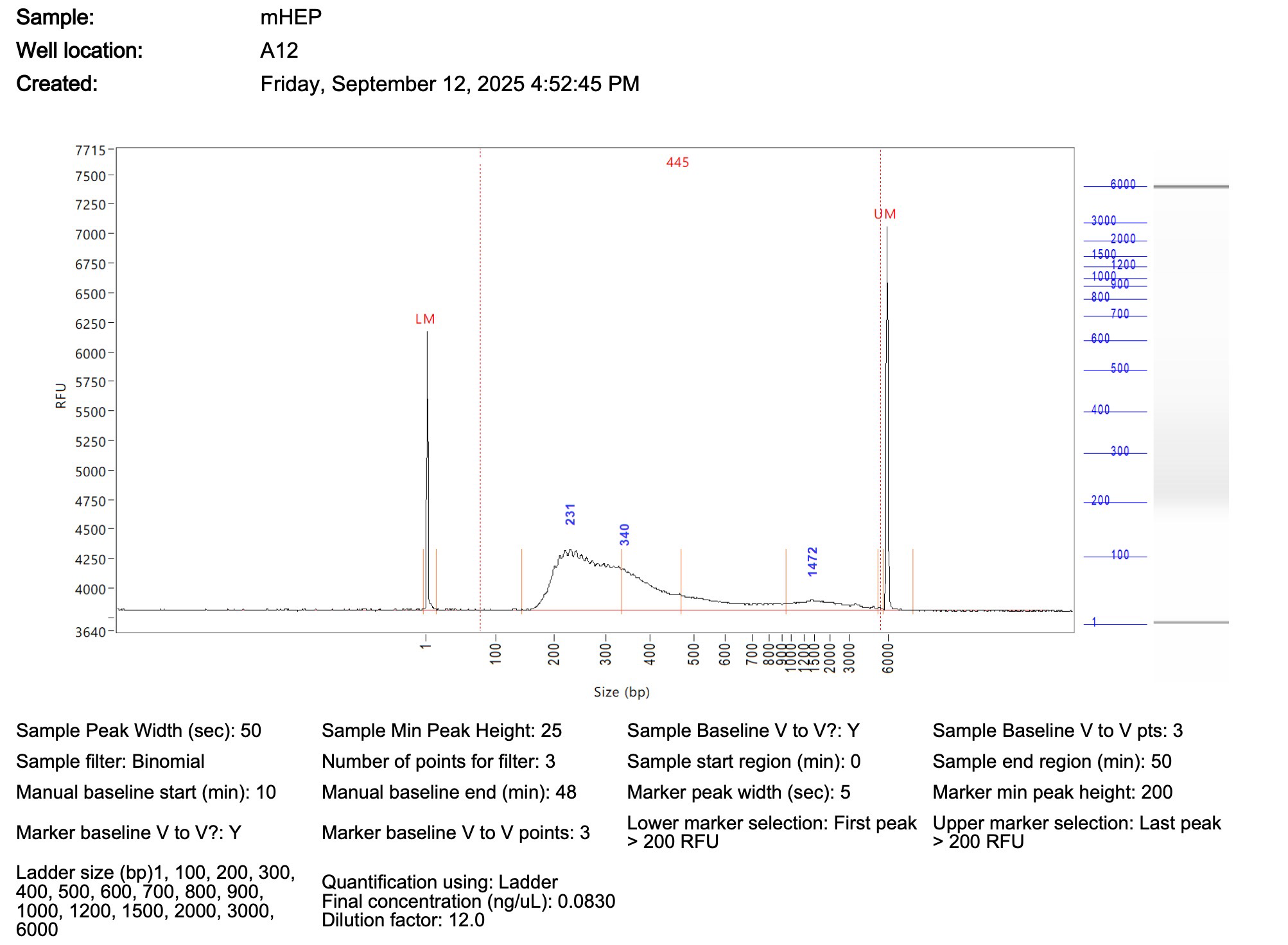

**Plate-based scHi-C - hESC stages- batch1**

<Plate1>

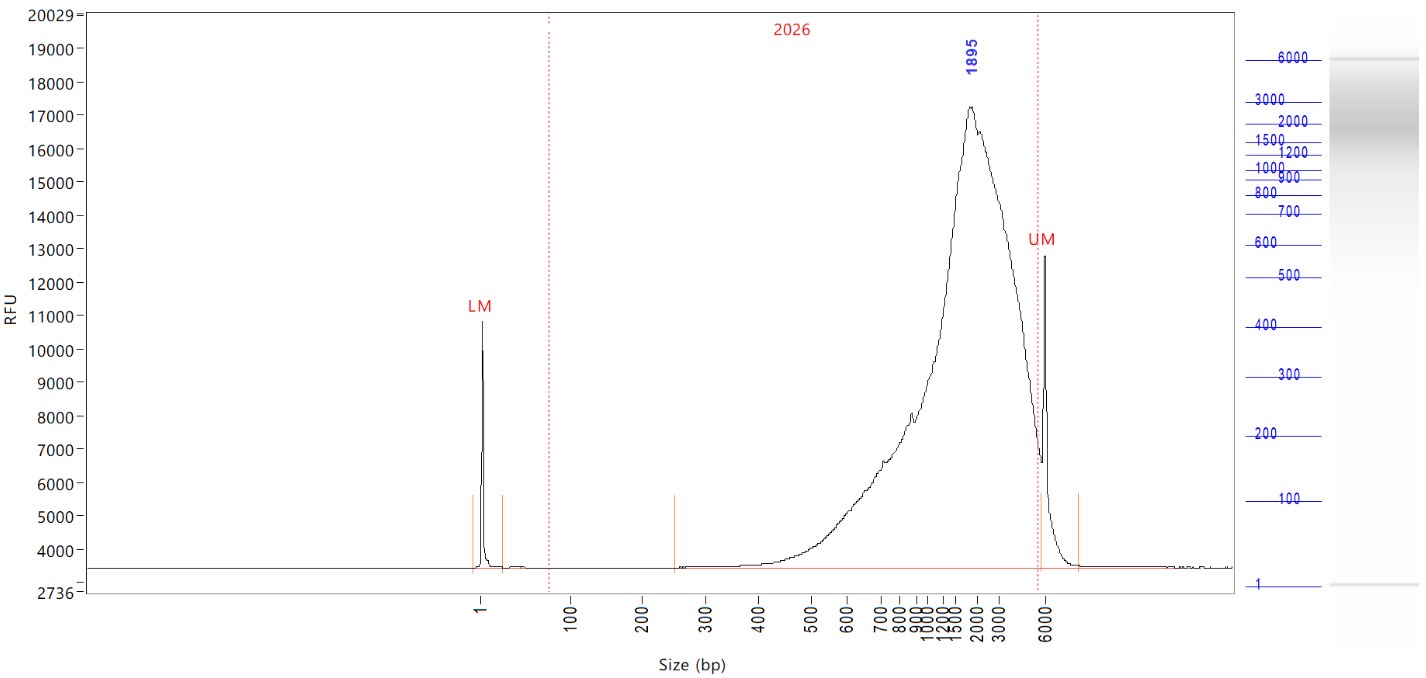

<Plate2>

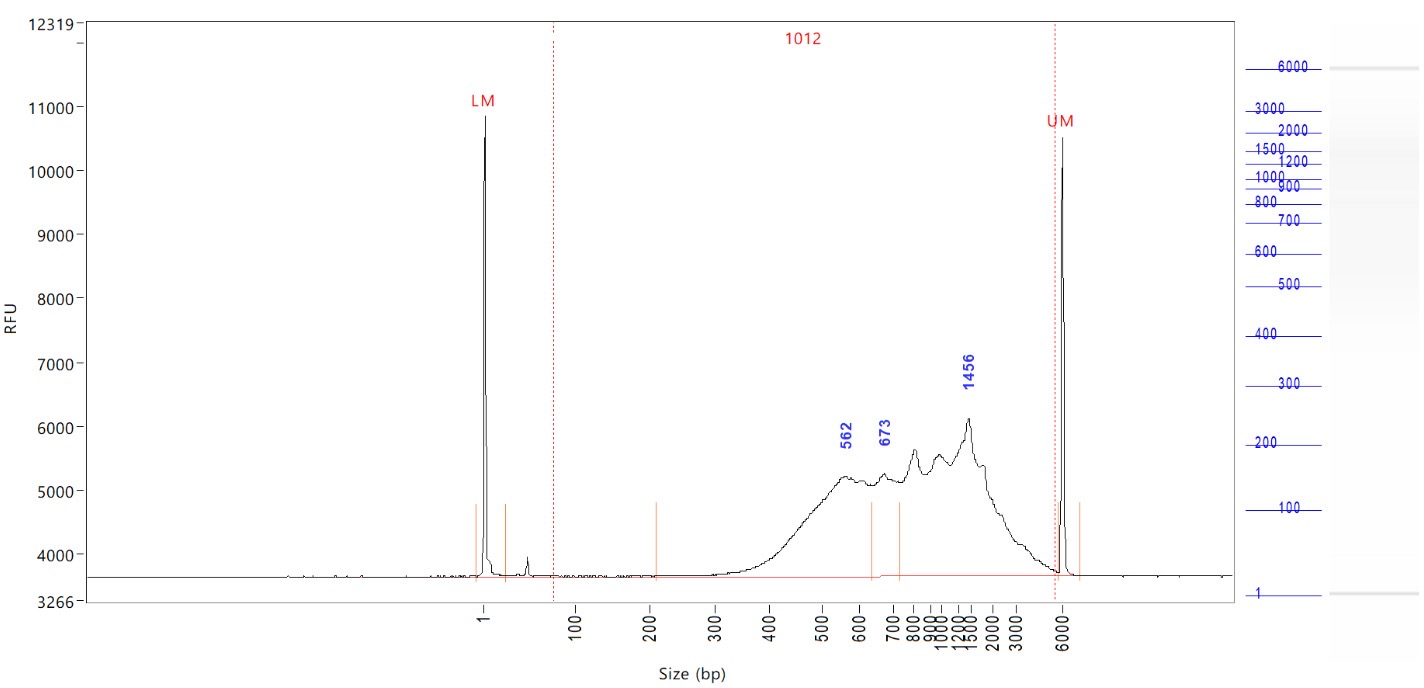

**Plate-based scHi-C - DE stages- batch1**

<Plate1>

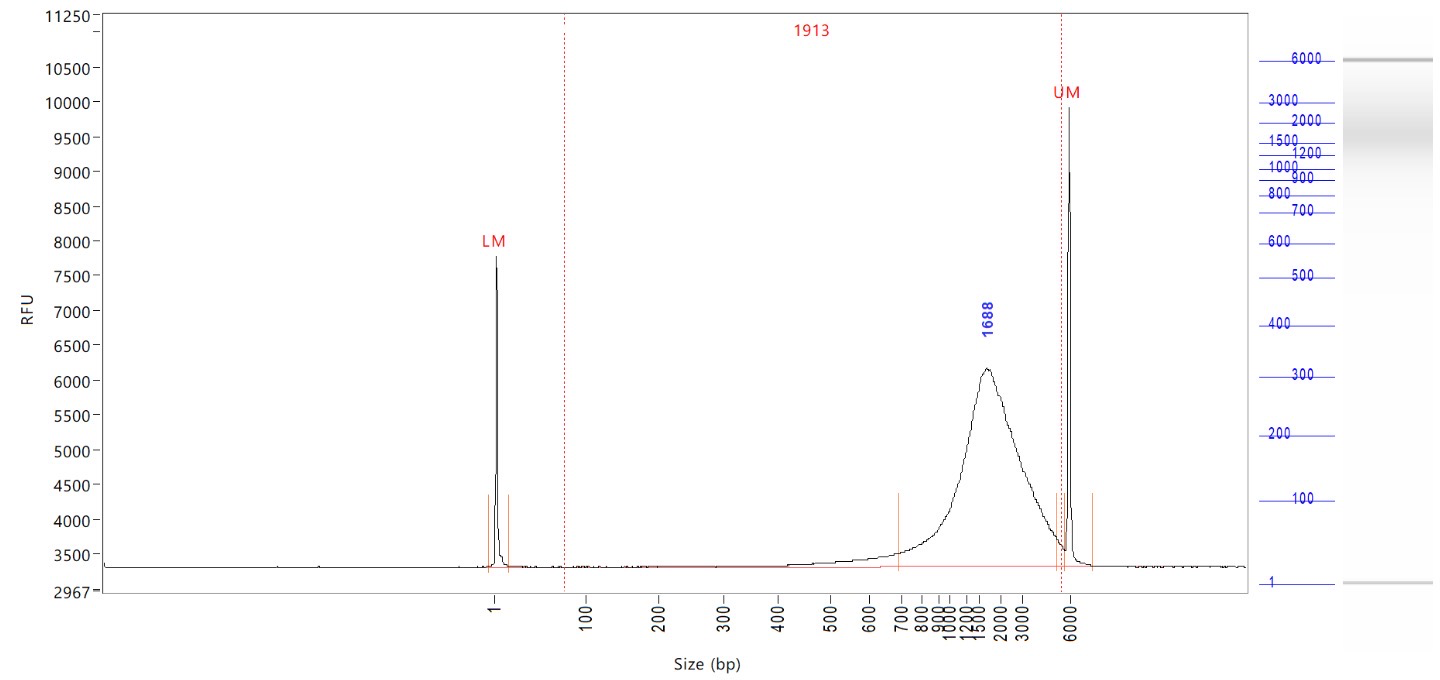

<Plate2>

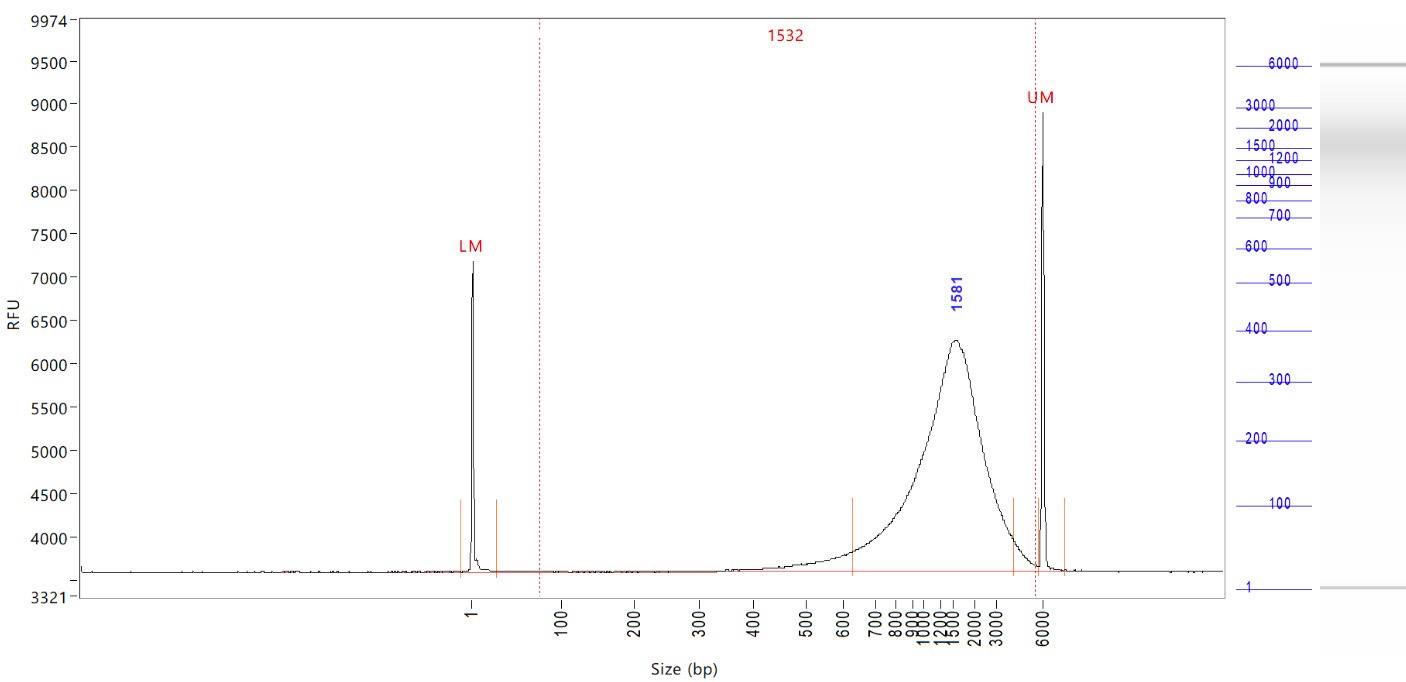

**Plate-based scHi-C - HB stages- batch1**

<Plate1>

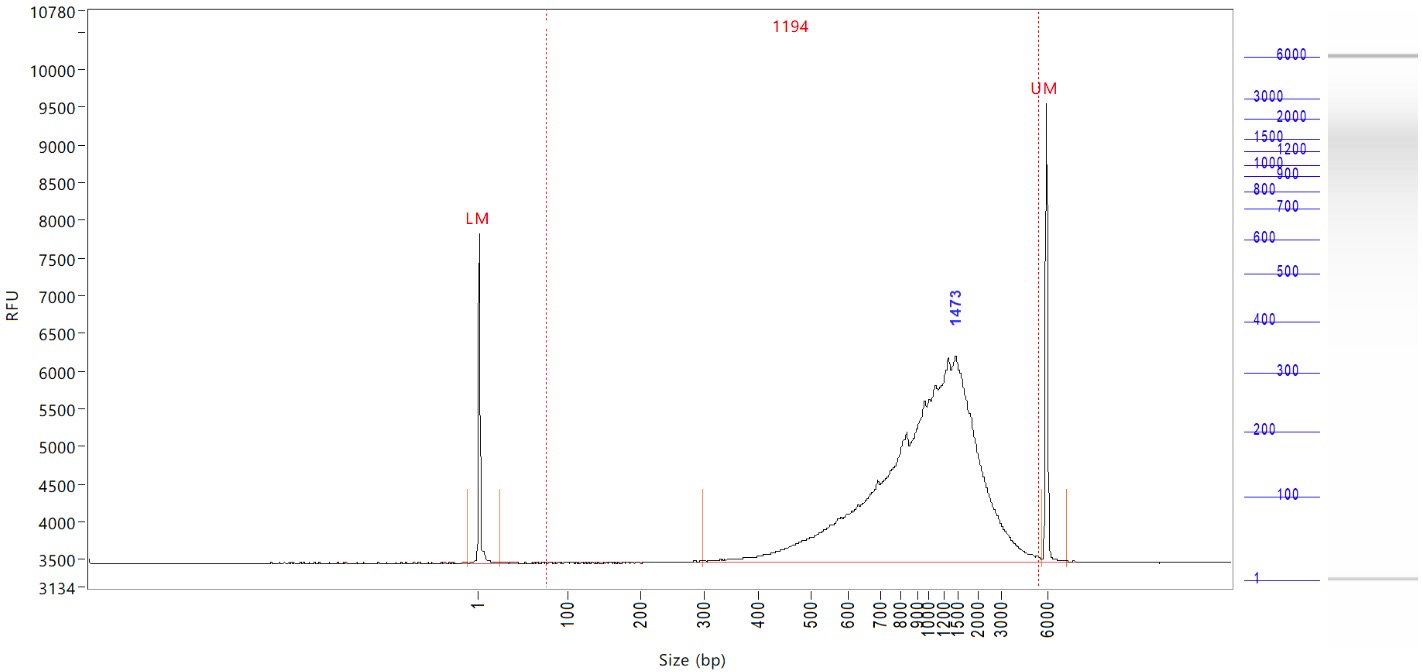

<Plate2>

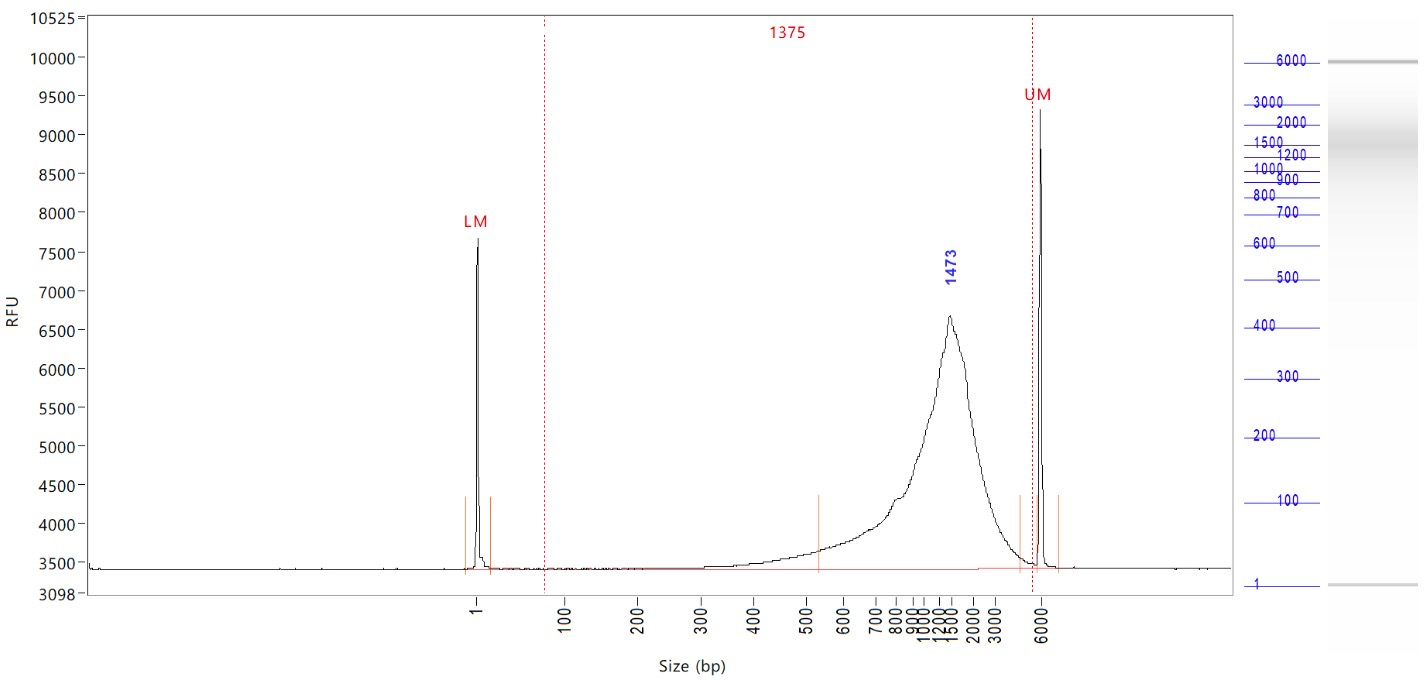

**Plate-based scHi-C - iHLC stages- batch1**

<Plate1>

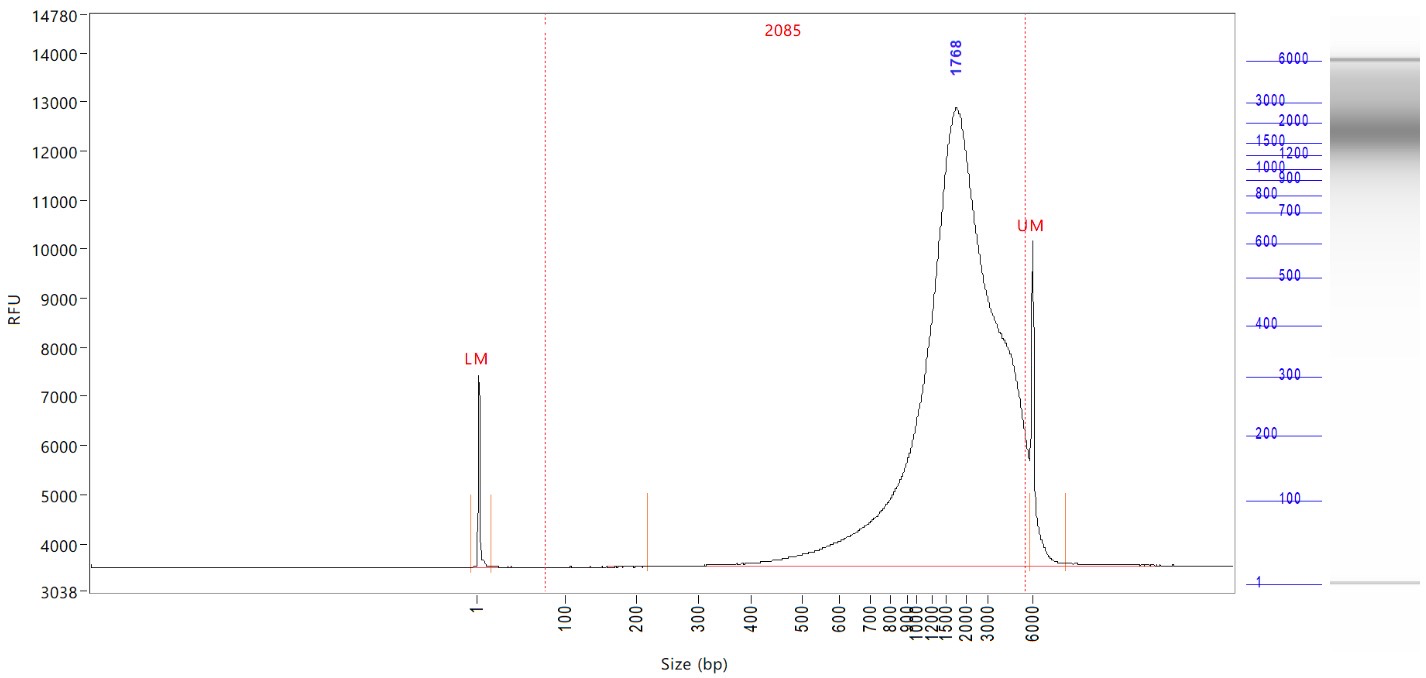

<Plate2>

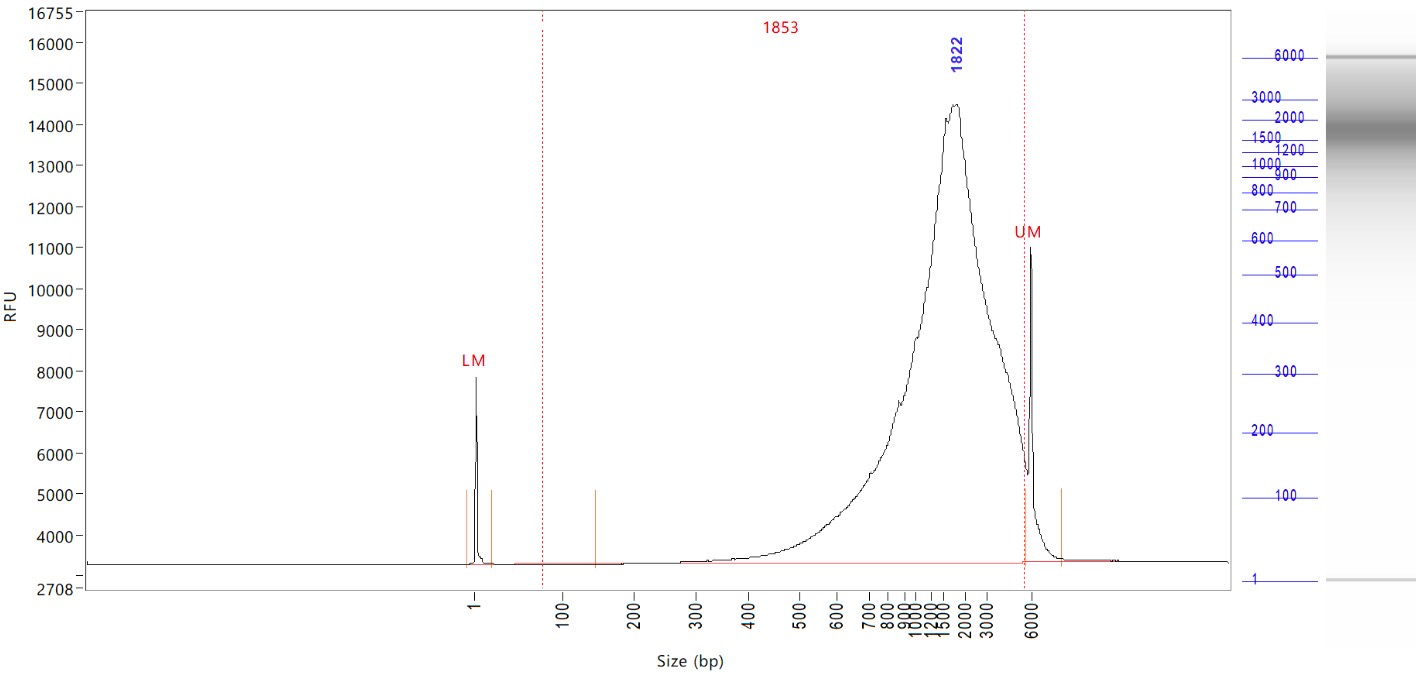

**Plate-based scHi-C - HLC stages- batch1**

<Plate1>

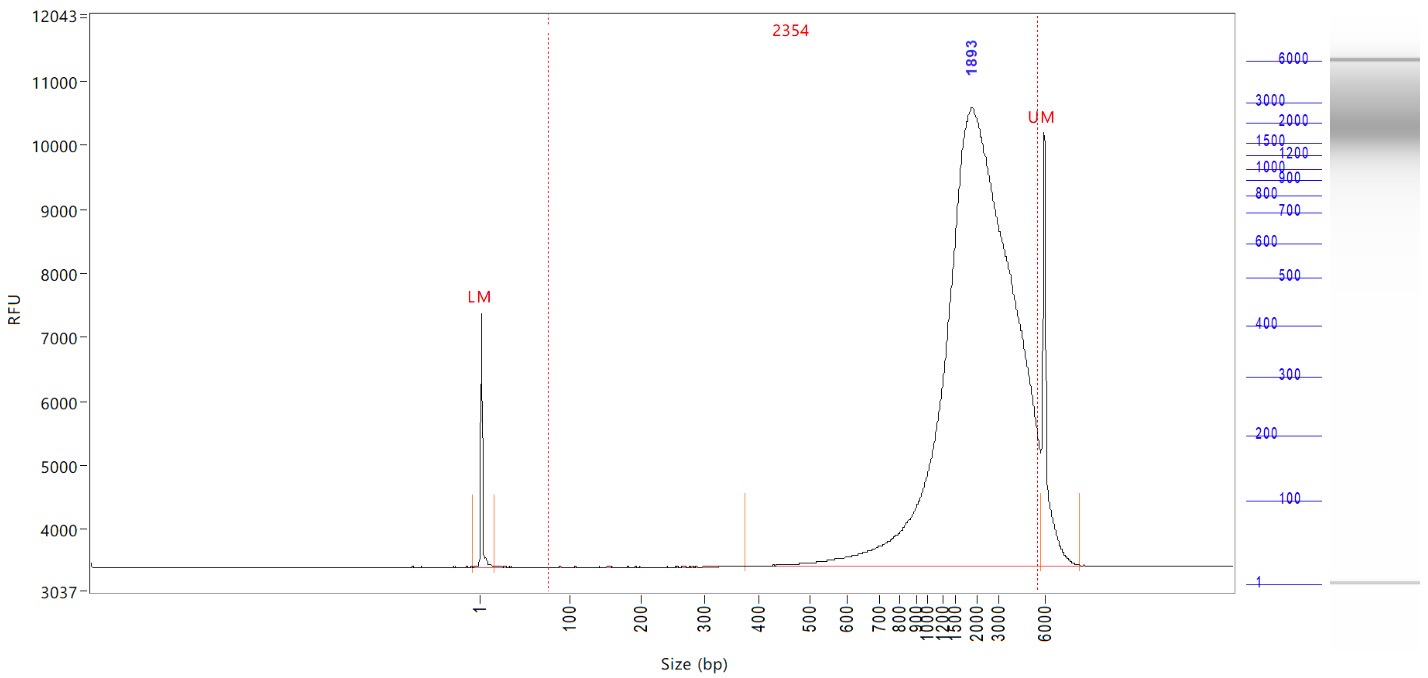

<Plate2>

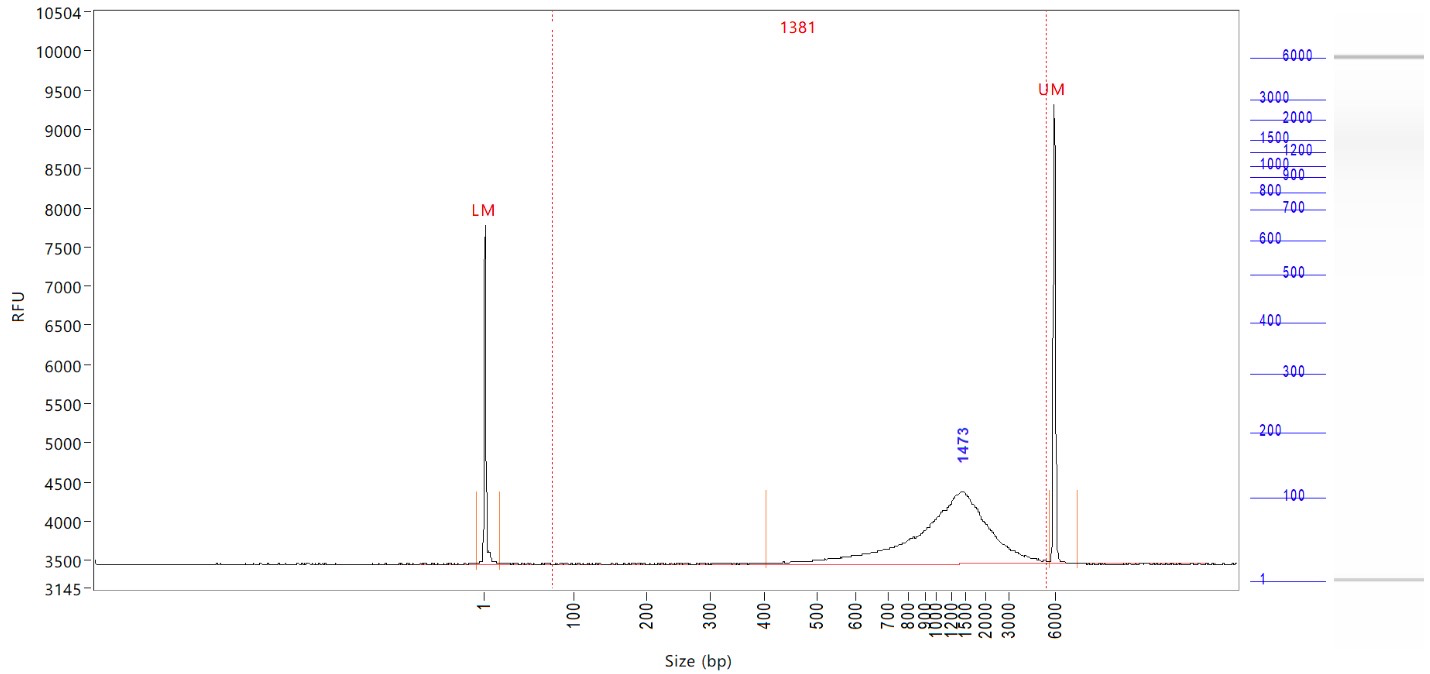

**Plate-based scHi-C - hESC stages- batch2**

<Plate1>

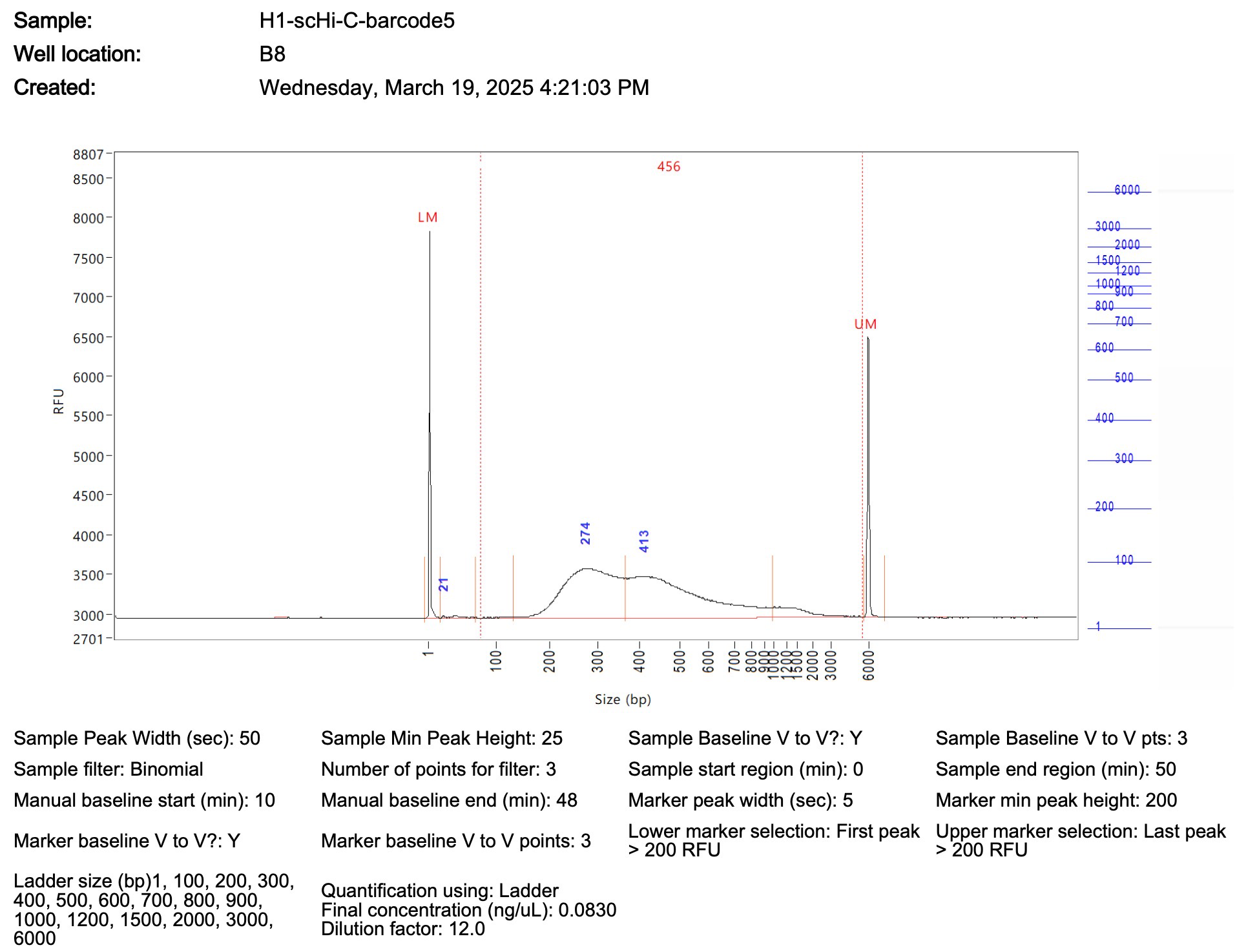

<Plate2>

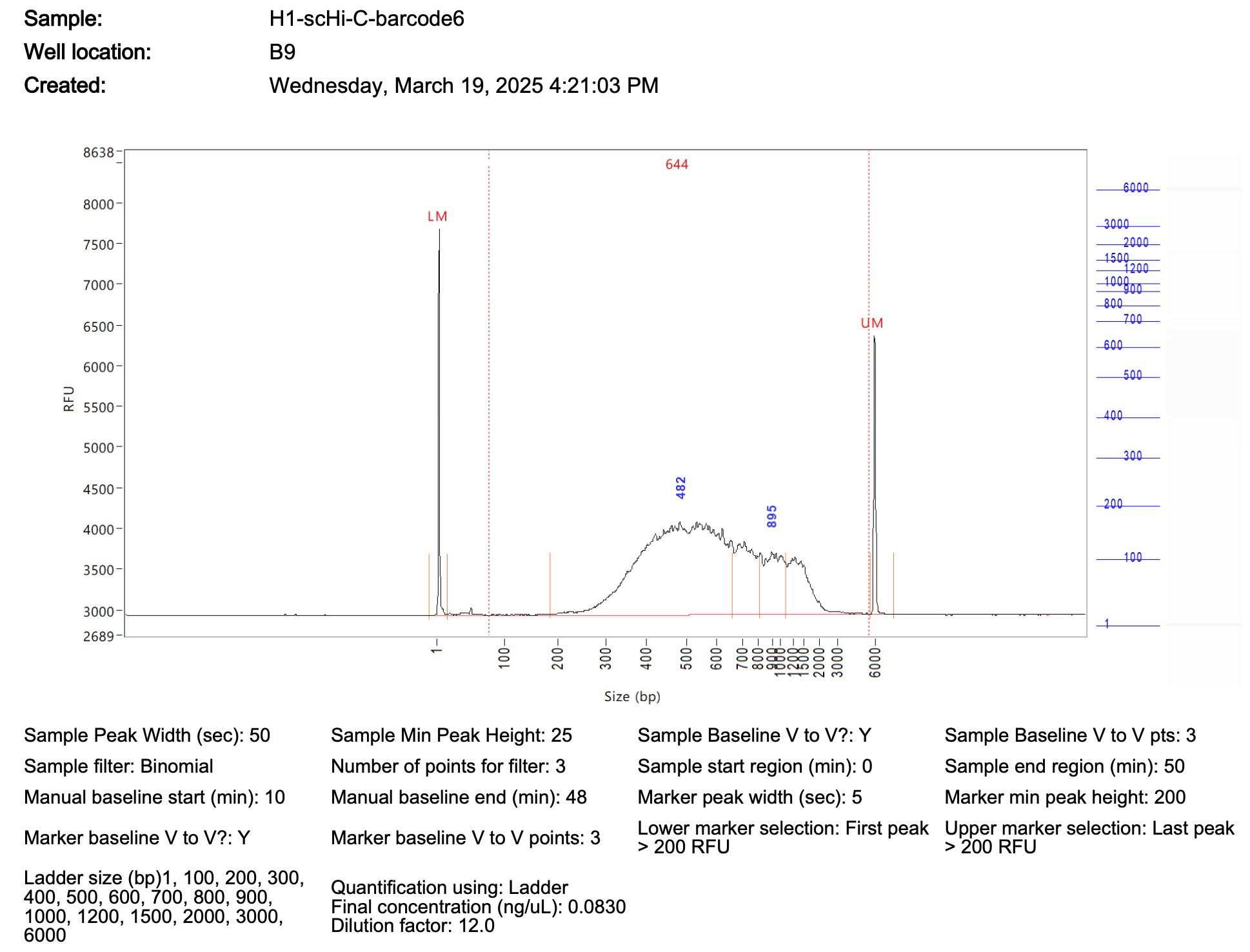

**Supplemental Note 3**

**Separating sampling noise from biological variability**

A further property to validate is how much of the cell-to-cell difference in a single-cell cis/total contact ratio is sampling noise and how much is biological variability. We analyzed droplet-based and plate-based scHi-C of five differentiation stages (ES, DE, HB, iHLC and HLC) in autosomal, non-centromeric 1 Mb bins; for cell *i* and bin *j*, *n_ij_* is the number of contacts touching bin j and *x_ij_* the number of these that are cis, with each contact counted once. Cells passed the quality filters described in Methods. The quantity of interest is the cell-to-cell variability of the cis/total ratio *x_ij_*/*n_ij_* within a stage.

Two features of this quantity complicate its interpretation. First, the variability observed across cells mixes biological variability and sampling noise contributions: when a cell yields few contacts, sampling noise dominates, and *x_ij_*/*n_ij_* can vary widely even if the underlying cis propensity is identical in every cell. Second, *x_ij_*/*n_ij_* is a proportion, and its variance is bounded by *μ_j_*(1 − *μ_j_*) and depends on the mean: bins whose cis fraction is near 0 or 1 have little room to vary, whereas bins near 0.5 can vary more.

To separate the two contributions we used a two-layer model, fitted independently for each bin. Conditioned on its *n_ij_* contacts, a cell's number of cis contacts is binomial with the cell's true cis propensity *p_ij_*; across cells, *p_ij_* scatters around the bin mean *μ_j_* with a precision *κ_j_*, larger *κ_j_* giving a tighter beta distribution:

*x_ij_* | *p_ij_* ~ Binomial(*n_ij_*, *p_ij_*)
 *p_ij_* ~ Beta(*μ_j_* *κ_j_*, (1 − *μ_j_*) *κ_j_*)	(1)

The binomial layer has E[*x_ij_*/*n_ij_* | *p_ij_*] = *p_ij_* and Var(*x_ij_*/*n_ij_* | *p_ij_*) = *p_ij_*(1 − *p_ij_*)/*n_ij_*. Applying the law of total variance and using E[*p*(1 − *p*)] = *μ*(1 − *μ*) − Var(*p*), the observed variance separates into a biological and a sampling term:

Var(*x_ij_*/*n_ij_*) = *μ_j_*(1 − *μ_j_*)[*ρ_j_* + (1 − *ρ_j_*)/*n_ij_*]	(2)

The first term does not depend on *n*, whereas the second shrinks as 1/*n*; for a cell with few contacts most of the observed spread is sampling noise. Dividing both sides by *μ_j_*(1 − *μ_j_*) removes the dependence on the mean and gives the dispersion factor *ρ_j_* = Var(*p_ij_*)/(*μ_j_*(1 − *μ_j_*)) = 1/(*κ_j_* + 1). It is dimensionless and lies between 0 and 1: *ρ_j_* = 0 means that all variability is sampling noise, and *ρ_j_* = 1 means that the true cis fraction varies as widely as the bin mean allows.

*μ_j_* and *ρ_j_* were estimated by the method of moments:

*μ_j_* = Σ*_i_* *x_ij_* / Σ*_i_* *n_ij_*
 *ρ_j_* = Σ*_i_* [(*x_ij_* − *n_ij_* *μ_j_*)² − *n_ij_* *μ_j_*(1 − *μ_j_*)] / Σ*_i_* *n_ij_*(*n_ij_* − 1) *μ_j_*(1 − *μ_j_*)	(3)

Bins were estimated when at least 20 cells had two or more contacts, retaining about 2,650–2,675 1Mb bins per stage.

Because the technical term depends on *n*, stages sequenced to different depth would differ in observed variability even at equal *ρ*. Stages were therefore compared at matched depth: for each stage its N deepest cells were taken, and the stage whose N deepest cells had the lowest median depth served as the reference (HLC, N = 100, for droplet data; ES, N = 35, for plate-based data). Every other stage contributed the N cells that best matched the reference cells one to one on depth, with no cell used twice; contacts were not down-sampled.

In both platforms, the median per-bin *ρ* was highest in ES and lowest in HLC (**Supplemental Fig. 2 in below**): 0.052, 0.033, 0.036, 0.050 and 0.009 for droplet-based data, and 0.140, 0.036, 0.062, 0.057 and 0.034 for plate-based data (ES, DE, HB, iHLC and HLC). The negative correlation between per-bin *ρ* and mean(cisratio) indicating elevated cell-to-cell variability at active chromatin (**Supplemental Fig.3- 4 in below**)

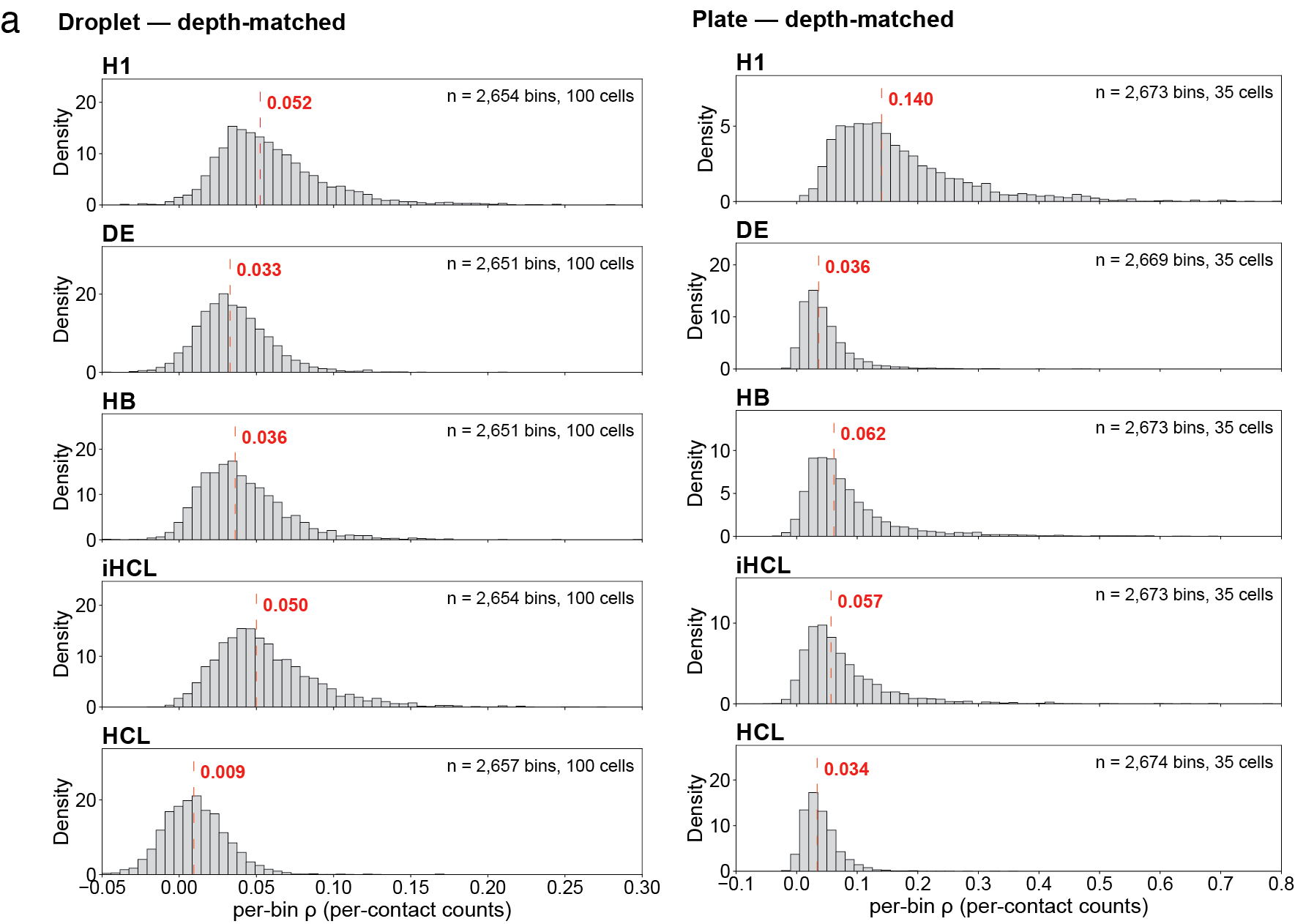

**Supplementary Fig. 2** | **Cell-to-cell variability of the cis/total contact ratio across differentiation stages after correction for sampling noise.** Distributions of the per-bin dispersion factor ρ, estimated by the method of moments from per-contact counts, over 1 Mb bins; one panel per stage. a, Droplet-based scHi-C, 100 depth-matched cells per stage. b, Plate-based scHi-C, 35 depth-matched cells per stage. The dotted line gives the median over all estimable bins; each panel lists bin and cell numbers. Cis contacts ≥0.4 kb (droplet) or ≥0.3 kb (plate) and all trans contacts are counted; a bin requires at least 20 cells with at least two contacts.

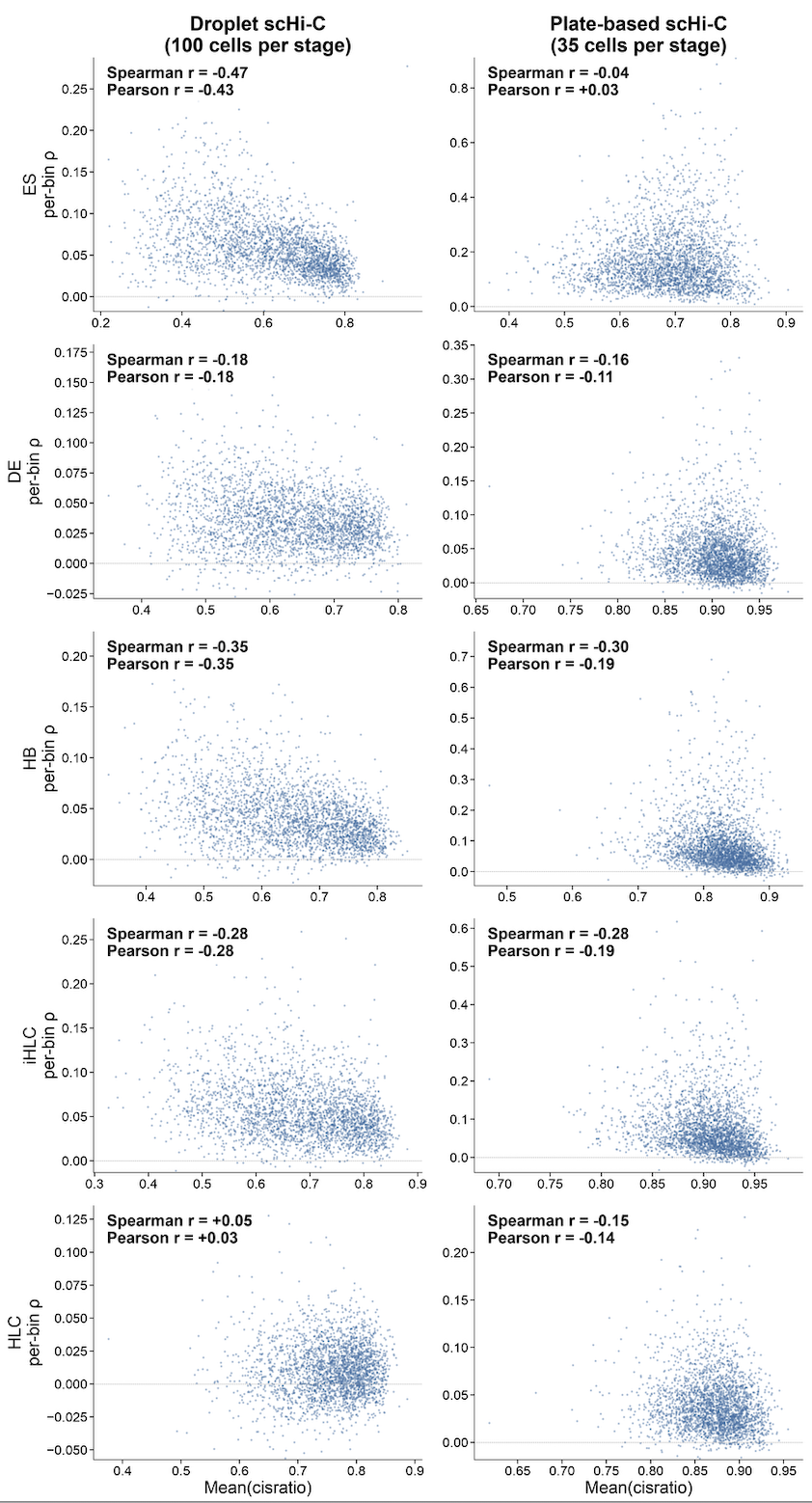

**Supplementary Fig. 3 | Relationship between the cell-to-cell folding variability of the cis/total contact ratio and the mean cis/total contact ratio across differentiation stages after correction for sampling noise.** Per-bin dispersion factor ρ, are plotted against the mean cis/total contact ratio of the same bin for 1 Mb bins; one row per stage (ES, DE, HB, iHLC, HLC). a, Droplet-based scHi-C, 100 depth-matched cells per stage. b, Plate-based scHi-C, 35 depth-matched cells per stage. Each point is one estimable bin; Spearman’s and Pearson’s correlation coefficients are given in each panel, and the dotted line marks ρ = 0. ρ and Mean(cisratio) are computed from the same cells (those with at least two contacts in the bin). Cis contacts ≥0.4 kb (droplet) or ≥0.3 kb (plate) and all trans contacts are counted; a bin requires at least 20 cells with at least two contacts.

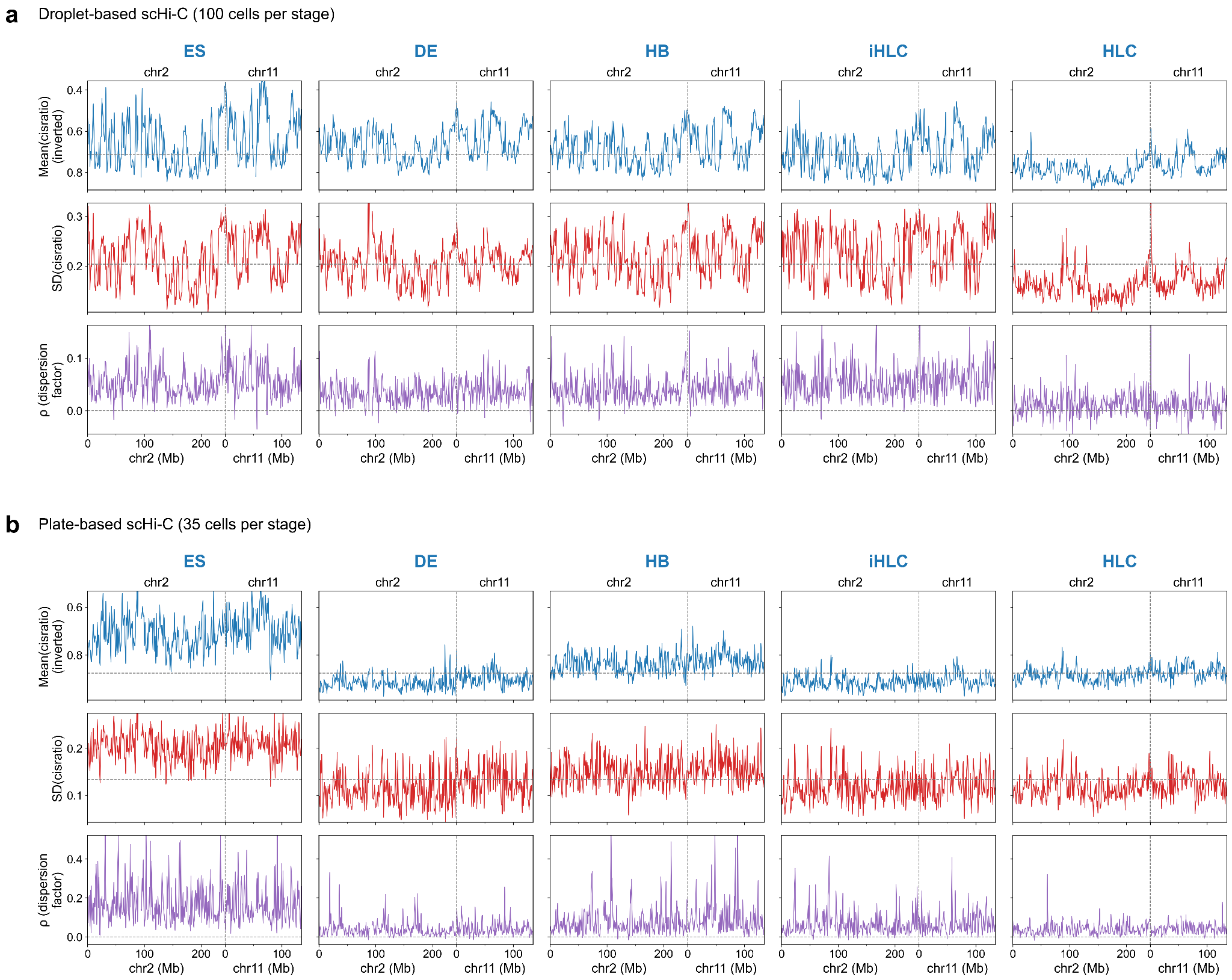

**Supplementary Fig. 4 | Cell-to-cell folding variability of the cis/total contact ratio along chromosomes 2 and 11 across differentiation stages.** Per-bin Mean(cisratio), SD(cisratio) and dispersion factor ρ are plotted along chromosome 2 followed by chromosome 11 for 1 Mb bins; one column per stage (ES, DE, HB, iHLC, HLC). a, Droplet-based scHi-C, 100 depth-matched cells per stage. b, Plate-based scHi-C, 35 depth-matched cells per stage. In each panel, rows show from top to bottom Mean(cisratio) (axis inverted), SD(cisratio) and ρ, the beta-binomial dispersion factor that corrects the cell-to-cell variability for sampling noise. Mean(cisratio), SD(cisratio) and ρ are computed from the same cells (those with at least two contacts in the bin). Grey dashed horizontal lines mark the median over all plotted bins and stages of the same platform (ρ = 0 for ρ); the grey dashed vertical line marks the junction between chromosomes 2 and 11. Cis contacts ≥0.4 kb (droplet) or ≥0.3 kb (plate), with contacts in the same bin counted once; a bin requires at least 20 cells with at least two contacts, and gaps mark centromere/gap bins or bins that fail this requirement.

**Supplemental Note 4**

**Distinct interaction constraints shape folding variability and radial positioning**

To further disentangle the factors contributing to folding variability, we conducted IGM-based simulations in which we selectively left out specific datatypes to assess their contributions to reproducing folding variability. As a baseline, we first generated 1,000 genome structures using an unrestrained polymer model without Hi-C-derived restraints and initialized from randomized compact conformations approximating chromosome territories (Methods) (**Supplemental Fig. 5a, b in below**, left most panels). Under these conditions, genomic regions lost their characteristic differences in both folding behavior and spatial localization, indicating that the observed variability patterns do not arise as an intrinsic property of random polymer chains alone.

We next investigated whether differentiation-associated changes in nuclear morphology and volume could account for the reduction in folding variability. Using the ES Hi-C interaction data, we generated genome structures under two shape–volume configurations: nuclei with an ES-like shape and an ES-like volume and an HB-like shape and volume. Adopting the more flattened nuclear shape and volume of HB cells had only minimal effect, with a very small, although significant increase in overall median CT-based folding variability and no effect on the chromosome compactness within the nucleus. (**Supplemental Fig. 5c**). Thus, global changes in nuclear morphology and volume alone are not the phenomena to reproduce the differentiation-associated decrease in local chromosome folding variability.

Next, to dissect the contributions of trans interactions, cis interactions, and lamina association to folding variability, we applied a computational gain-of-function strategy. Starting from the unrestrained polymer model, we independently reintroduced trans, cis, and LaminB-DamID restraints and assessed their effects on folding variability and locus-specific subnuclear positioning (**Supplemental Fig. 5b**). These simulations test which restraint classes recover features of the fully constrained profile, rather than quantitatively partitioning their causal contributions. Within this framework, our results suggest that global folding-variability patterns emerge from the combined contribution of distinct classes of interaction restraints.

Folding variability was substantially higher in the unrestrained polymer model than in the fully restrained reference model (median S.D., 0.82 versus 0.66) (**Supplemental Fig. 5a the left chart, Supplemental Fig. 5b, the first left model**). Moreover, the locus-specific variability pattern observed in the reference model was completely lost in the unrestrained ensemble, with essentially no correlation between genomic regions in the two conditions ((r=-0.06)). Reintroducing any class of experimental interaction restraints reduced folding variability relative to the unrestrained ensemble (median S.D., 0.55–0.72, labeled as red dash lines) (**Supplemental Fig. 5b, models from second left to right)**, indicating that experimental constraints generally restrict the accessible conformational space of genomic regions.

Cis restraints reproduced the overall magnitude of folding variability observed in the reference model (median S.D., 0.62 versus 0.66) but showed only a small correspondence to the locus-specific pattern (r = 0.27) (**Supplemental Fig. 5a the left chart, Supplemental Fig. 5b, the fourth left model**). This may reflect that cis interactions constraint internal chromosome folding but provide limited information about how far individual regions extend beyond the chromosome territories. In contrast, trans restraints directly constrain loci in interchromosomal space and therefore better specify to what extent specific regions loop out beyond their chromosome territories. Consistently, models from only trans interactions closely reproduced the locus-specific pattern of CT-based folding variability observed in the reference model (r = 0.91), although their overall variability was lower (median S.D., 0.58) (**Supplemental Fig. 5a the left chart, Supplemental Fig. 5b, the third left model**). Thus, both cis- and trans-only models shifted the overall folding variability toward lower values compared to the reference model (cis+trans model, median S.D. 0.66).

The effect of LaminB-DamID restraints on folding variability depended on the interaction context in which they were applied. LaminB-DamID alone reproduced remarkably well the overall folding variability (median S.D. 0.64), as well as the locus-specific variability pattern observed in the reference model (r = 0.86), even though no chromatin-chromatin interactions were introduced (**Supplemental Fig. 5a the left chart, Supplemental Fig. 5b, the second left model**). Combining LaminB-DamID restraints with trans interactions only slightly increased the correlation with the reference model (r=0.86), while decreasing the overall variability measure, indicating that both data types provide complementary information (median S.D. 0.55) (**Supplemental Fig. 5a the left chart, Supplemental Fig. 5b, the first right model**).

Finally, we asked which restraint classes encode locus-specific radial positioning. Trans restraints alone and to a lesser extent LaminaB-DamID restraints alone were sufficient to reproduce the radial localizations observed in the fully constrained model (r = 0.99 and r=0.89, respectively), but did not recapitulate compact chromosome territories. In contrast, cis restraints alone readily produced compact chromosome territories but failed to recover locus-specific radial positioning (r = -0.17). Combining cis with trans restraints or cis with LaminaB-DamID restraints is required to recover both chromosome territoriality and radial nuclear positioning (**Supplemental Fig. 5a, the right chart, Supplemental Fig. 5b**).

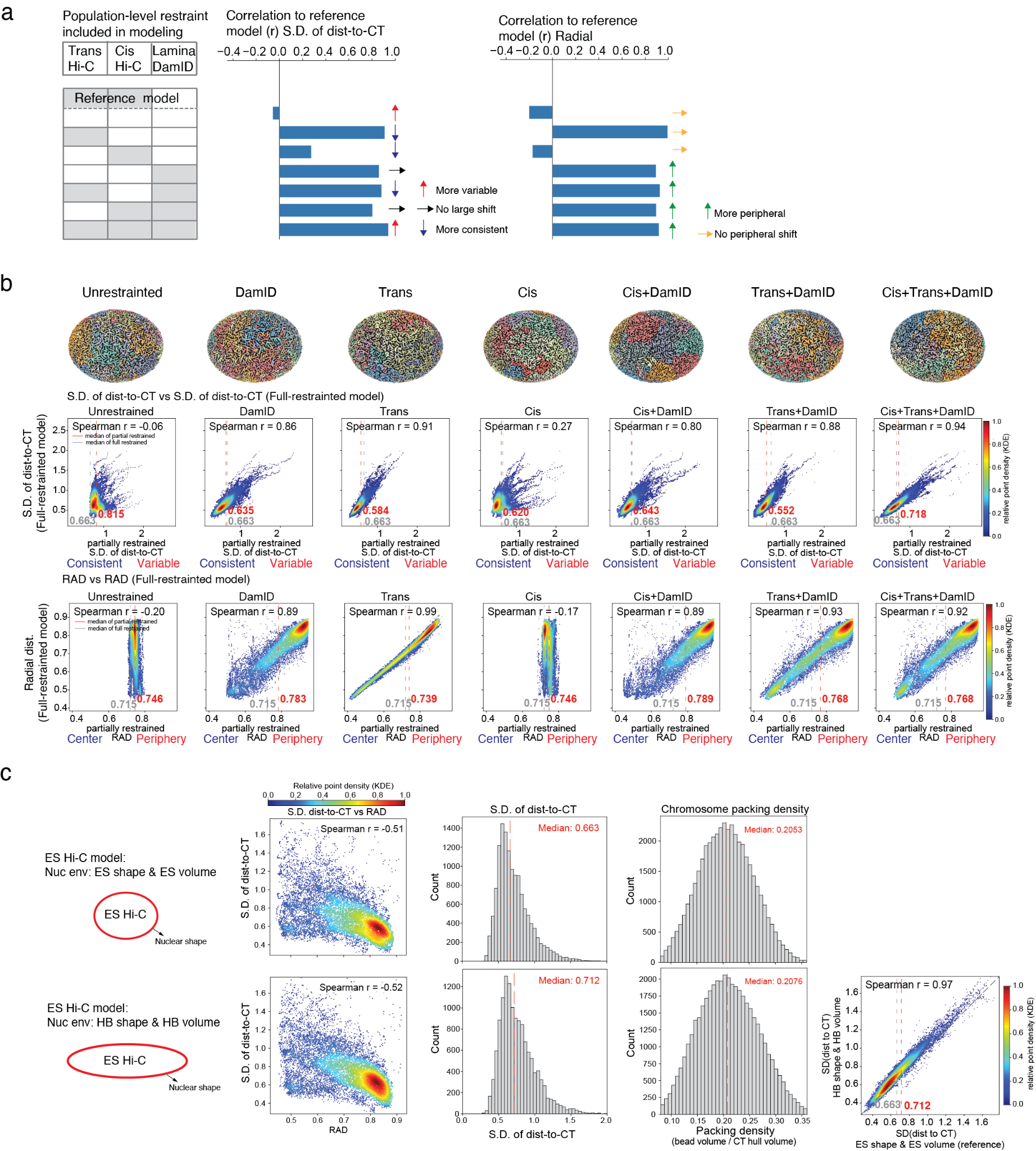

**Supplemental Fig. 5: | Distinct interaction constraints shape folding variability and radial positioning**

**(a)** Summary of simulation results from conditionally restrained models testing contributions of different structural restraints as in Fig S5D. From left to right are: table of the combinations of the restraints used in this simulation; bar plots of the correlations between the S.D. of dist.-to-CT generated from simulated model with that from the full restrained model (cis + trans interactions), where arrows indicate change in direction of S.D. of dist.-to-CT in simulated model from the full restrained model; bar plots of the correlationships between the radial localization generated from simulated model with that from the full restrained model (cis + trans interactions); where arrows indicate change in direction of radial localization in simulated model from the full restrained model.

**(b)** Simulation results from conditionally restrained models testing contributions of different structural restraints. Top: representative genome structure generated under each restraint condition. Middle: scatter plots comparing the S.D. of dist.-to-CT from each conditionally-restrained model (x-axis) versus the S.D. from the full-restrained model (y-axis). Bottom: scatter plots comparing normalized radial position of genomic bins between conditionally-restrained model (x-axis) versus that from full-restrained model (y-axis). Restraint conditions (left to right) include: polymer without Hi-C and DamID restraint, DamID restraints, trans-chromosomal Hi-C restraints, cis-chromosomal Hi-C restraints, cis-chromosomal Hi-C + DamID restraints, trans-chromosomal Hi-C + DamID restraints, and combined cis- and trans- chromosomal Hi-C + DamID restraints. Red dotted lines mark the median S.D. of dist.-to-CT (or radial position) of the partially restrained model shown in each panel, with its value given next to the line; grey dotted lines mark the median of the fully restrained (cis + trans) model (S.D. of dist-to-CT) = 0.663; radial position = 0.719).

**(c)** First column: Diagrams illustrating the nuclear shapes implemented in the conditional models. Second column: Scatter plots of nuclear radial distance versus the S.D. of dist.-to-CT for 200-kb genomic bins in the ES-stage generated from the corresponding conditional models. Centromeric regions were excluded. Third column: Histograms of the S.D. of dist.-to-CT for 200-kb genomic bin across different conditional models. Red dashed lines mark the median S.D. for each condition. Fourth column: Histograms of the radius of gyration for the whole chromosome at 200-kb genomic bin across different conditional models. Histograms show the compactness score computed from each modeled structure, pooled across all autosomes for four ES-condition model variants. Red dashed lines mark the median compactness score for each condition. Fifth column: scatter plots comparing the S.D. of dist.-to-CT from tested model (y-axis) versus the S.D. of dist.-to-CT from the ES model (x-axis)

**References**

1.	Ramani, V. *et al.* Massively multiplex single-cell Hi-C. *Nat Methods* **14**, 263–266 (2017).

2.	Nagano, T. *et al.* Cell-cycle dynamics of chromosomal organisation at single-cell resolution. *Nature* **547**, 61–67 (2017).

3.	Naumova, N. *et al.* Organization of the mitotic chromosome. *Science* **342**, 948–953 (2013).
